# Novel multifunctional glycan probes to elucidate specificities of diverse glycan recognition systems

**DOI:** 10.1101/2023.07.09.548266

**Authors:** Jin Yu, Alexiane C. Decout, Antonio Di Maio, Wengang Chai, Thilo Stehle, W. Bruce Turnbull, Ten Feizi, Yan Liu

## Abstract

Glycan microarrays of sequence-defined glycans are the most widely used approach for high-throughput studies of interactions of glycans with glycan binding proteins. Currently nitrocellulose (NC) coated or N-hydroxysuccinimide (NHS) functionalized glass slides are the two most commonly used surfaces for immobilizing glycan probes as noncovalent and covalent arrays, respectively. The mode of glycan presentation can have an influence on the microarray readouts and is an important consideration in the glycan recognition knowledgebase. Here we present the development of glycan probes, tagged with a novel type of tri-functional Fmoc-amino-azido (FAA) linker, which can be readily converted into lipid-tagged or amino-terminating glycan probes for generating neoglycolipid (NGL)-based noncovalent arrays as well as covalent arrays. The azido functionality in the FAA glycan probes provides a means to ‘light up’ the glycans via biorthogonal ‘Click’ chemistry so that they become ‘scanner-readable’ on the covalent arrays, which represents an advance in array quality control. Here analyses were carried out with a diverse set of 36 glycan binding proteins (GBPs) to compare the performance of 44 glycans presented in a liposomal formulation as noncovalent arrays on NC coated slides and on two types of NHS slides as covalent arrays. With most of anti-glycan antibodies and plant lectins investigated, there were negligible or subtle differences in the binding detected in different arrays. However, there were some striking differences observed between the covalent and noncovalent arrays. These include binding of *Lotus Tetragonolobus* Lectin (LTL) on the covalent but not the noncovalent arrays, and a clear preference observed for glycans on the noncovalent array in the binding of adhesins (VP1 proteins) of two human polyomaviruses JCPyV and BKPyV, and the human immune lectins Siglecs 7 and 9. Subtle yet significant differences in the binding to low affinity glycan ligands were also observed with three bacterial toxins in different arrays. These results revealed interesting insights into the binding behaviour of different GBPs on the noncovalent and covalent arrays and highlight the importance to consider different array platforms in elucidating glycan mediated interactions. The FAA glycan probes can be readily rendered fluorescent via ‘Click’ chemistry in solution, enabling the detection of GBPs at the surface of cells.

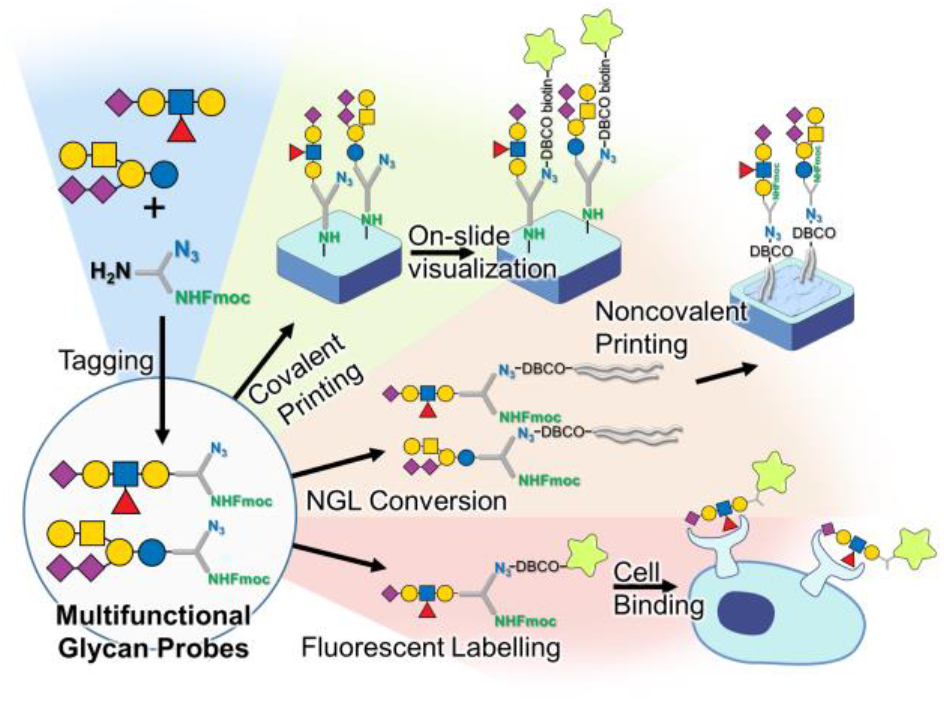

A new trifunctional linker can present glycans on different array surfaces for analyses of soluble glycan binding proteins or as fluorescent probes for binding by proteins at the surface of cells.

## Introduction

Glycans are ubiquitously displayed at the surface of cells and their surroundings in all living organisms. They interact with numerous binding partners, endogenous and exogenous, and mediate many key biological processes, including cell-cell adhesion, signalling, intracellular trafficking and two-way microbe-host interactions^1, 2^. On the one hand, viruses^3^, fungi^4^, bacteria^5^ and parasites^6^ often select and use host cell surface glycans for attachment at the initial stages of infection or colonization. On the other, the host responds to infective agents through interactions of the microbial glycans with carbohydrate recognizing antibodies and proteins of the innate immune system lectins^7^ and receptor like kinases^8^. Elucidation of the specifies of the diverse glycan binding proteins (GBPs) is crucial for our understanding of the glycan-mediated biological recognition events and pathways.

Glycan microarray technologies for sequence-defined glycans, first introduced in 2002^9^, have gained momentum internationally and have revolutionized the molecular dissection of glycan-protein interactions in a miniaturised and high-throughput fashion^10, 11^. Great advances have been made in the expansion of glycan libraries both through chemical synthesis approaches^12-15^ and isolation and characterization of glycan ligands from natural sources^9, 16-18^. To enable glycans to be immobilized on microarray surfaces for binding studies, a variety of linkers or tags have been developed^19-22^. The glycans can be converted into lipid-linked probes, neoglycolipids (NGLs), and immobilized noncovalently on nitrocellulose (NC) coated glass slides^23^, side-by-side with natural glycolipids (GLs). Alternatively, glycans can be covalently attached to microarray slides via various surface chemistries; among these the most popular are amine-terminating glycan derivatives on NHS functionalized glass slides^24, 25^.

In the various chemical derivatization procedures for glycan immobilization, there are factors such as the status of the core monosaccharide, the chemical properties of the linkers used, the density and the mode of display of the glycans^26, 27^. These can influence the microarray readouts and are important considerations in the glycan recognition knowledgebase. So far there are few reports on microarray platform comparisons in respect to binding signals elicited with a given glycan recognition system^28-31^, and such comparisons have been focused on data with plant lectins the majority of which give robust binding signals with arrayed glycan ligands. Comparative data on other, relatively low affinity glycan recognition systems are limited. A limitation thus far in all the slide-based planar array platforms has been the lack of a means to monitor and measure, using microarray scanners, the relative amounts of the glycans immobilized on the array surface. These are parameters to note in the quality control of arrays generated for glycan recognition studies.

Here we present the design of a novel tri-functional, Fmoc-amino-azido (FAA), linker. The FAA-linked glycan probes can be readily prepared from free reducing glycans and converted into amino-terminating as well as lipid-tagged glycan probes for generating covalent arrays and noncovalent arrays, respectively. This provides a unique opportunity for a close comparison of the two array platforms with a variety of glycan recognition systems. The azido functionality of the FAA probes, moreover, provides a means to ‘light up’ the glycans via biorthogonal ‘Click’ chemistry so that they become ‘scanner-readable’, both on array and in solution. This enables probing glycan-mediated interactions at intact cell surfaces.

## Results and discussion

### Design of the trifunctional FAA linker and preparation of FAA-linked glycan probes

The linker was designed with several features in mind: i) efficient conjugation with minute amounts of glycans and give single anomeric products to ensure the applicability of the linker for labelling and resolving heterogeneous natural glycan populations, ‘glycomes’ of cells and tissues; ii) incorporation of a fluorescent label in the linker to assist the purification of the derivatised glycans, enhance the sensitivity of detection during multidimensional chromatographic separations and facilitate quantitation of individual glycans or glycan fractions before arraying; iii) inclusion of an amino functional group to allow arraying of derivatised glycans directly onto functionalized glass slides using the well-established amide-coupling reaction; iv) conferring further versatility by including an additional azido functionality in the linker, to enable azide-alkyne cycloaddition (‘click’ chemistry) to be used for conversion of the glycan derivatives into other forms of functional glycan probes; these include for example lipid-linked, biotinylated, fluorescent forms that could be used in different settings for glycan recognition studies. Details of the syntheses are in the Supplemental Information.

The synthesis of the trifunctional linker was achieved by a two-step procedure using as a precursor, a commercially available Fmoc (fluorenylmethyloxycarbonyl)-protected azido-containing amino acid, Fmoc-L-azidoornithine (**Fig 1A**). The carboxyl group of the amino acid was activated and coupled to the amino group of N-Boc-ethylenediamine to give the Boc-protected amino-functionalized azidoornithine (compound **1**) as the single product in high yield. After purification it was deprotected by acid treatment to give Fmoc azido amino (FAA) linker, in quantitative yield.

**Figure 1.**
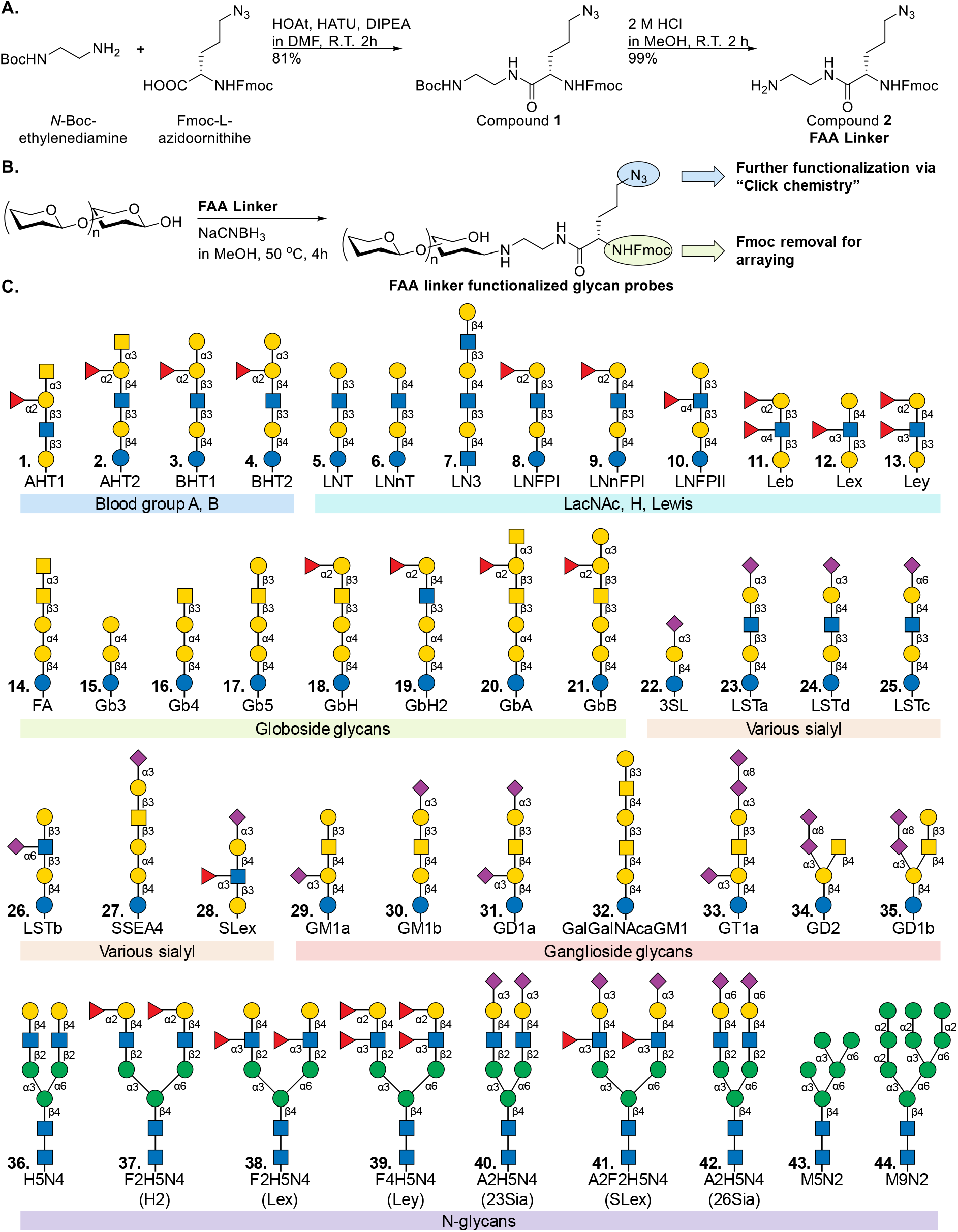
Schematic representation of the generation of FAA linker, the conjugation to glycans and symbolic sequences of forty-four FAA-tagged glycan probes for arraying. **A)** Synthesis of the FAA linker. **B)** Glycan conjugation to FAA linker. **C)** The forty-four FAA-conjugated glycan probes. Details of the linker synthesis, the sources and sequences of the glycan materials, their conjugation to the linker and their characterisation are in the Supplementary Information.

The FAA-linked glycan probes were prepared via microscale reductive amination whereby the reducing ends of glycans were conjugated to the free amine group of the FAA linker (**Fig 1B**). The efficiency of glycan conjugation was initially evaluated using Maltotriose at milligram scale. the starting glycans were fully labelled within 4 hours using 2 equivalents of the FAA linker as shown by thin layer chromatography (TLC). A glycan standard mixture of 6 oligosaccharides (3 nmol of each glycan) was labelled using 15 equivalents of FAA reagent at pH 6 (**Fig S5A**). The yield was determined by quantitation based on the fluorescence intensity of the product on HPLC. Almost full conversion of the starting glycans was achieved, regardless of sizes, neutral or acidic features, into the FAA-linked glycan probes. For labelling oligosaccharide mixtures released from glycoproteins which have the less reactive HexNAc at the reducing end, 20 equivalents of the FAA linker were used to achieve a yield above 80%. To test the efficiency of conjugation of more complex samples to the FAA linker, the *N*-glycans released from RNAse B and bovine fetuin were labelled at a scale of ∼10 μg with FAA, and two widely used aromatic glycan labeling reagents 2-aminobenzamide (2AB)^32^ and 2-amino-N-(2-amino-ethyl)-benzamide (AEAB)^33^. Whilst very similar HPLC profiles in terms of the retention times and peak shapes were observed with the *N*-glycans, the FAA-linked glycans gave much stronger fluorescence signals compared to the 2AB and AEAB derivatives (**Fig S5B-G**), consistent with previous findings with other Fmoc containing linkers^22^. The sensitivity of the FAA glycan labelling method was further demonstrated in a comparative HPLC-based quantitation of maltotriose labeled with FAA and 2AB (**Fig S7**). The FAA-linked maltotriose elicited a fluorescence signal that was 14 times higher than that of the 2AB labelled glycan. Moreover, the detection limit of the FAA-linked glycan was in the low pmol range, rendering the approach suitable for labelling natural glycans that are often available in limited in amounts.

The optimized procedure (20 eq FAA linker, pH 6, 50°C, 4 h) was used to label a total of 39 reducing glycans with the FAA linker; these included different types of blood group antigens, A, B, H and Lewis (Le^a, b, x, y^), sialyl glycans, globoside, ganglioside and *N*-glycans (**Fig 1C**). The FAA-linked glycan probes were purified and quantified by HPLC, the molecular weights were determined by MALDI TOF MS analyses (**Fig S6**).

The FAA glycan probes can serve as glycan acceptors in enzymatic reactions for further elongations or modifications. Starting from the asialo-biantennary *N*-glycan (probe #36), five additional *N*-glycan probes (#37-#41) with different modifications (sialylation and fucosylation) were enzymatically synthesized at 10-100 nmol scale. The presence of the hydrophobic Fmoc moiety facilitated chromatographic purification and monitoring of the synthetic products (**Fig S10**). The HPLC and MS characterization of the purified *N*-glycan probes are in **Fig S6**.

### ‘On-array’ visualization of immobilized FAA glycan probes

A limitation of the existing solid phase glycan array systems, to the best of our knowledge, is the lack of a means to quantify the glycans of different size and charge retained on the microarrays after deposition and the washing steps before analysis. For example the widely used AEAB-tagged glycan probes can be detected by fluorescence in solution, but the wavelength Ex330/Em420 wavelength^33^ is not compatible with the existing microarray scanner instruments for quantifying the arrayed spots using florescence readout.

With the FAA-glycan probes, the Fmoc protecting group in the linker can be readily removed to generate a primary alkyl amine for reacting with NHS functionalized glass microarray slides. The presence of the azido functionality in the FAA linker of the immobilized glycan probe allows introduction of a fluorescent group via the highly efficient copper-free click chemistry strain-promoted azide-alkyne cycloaddition (SPAAC) reaction. We have therefore used a two-step procedure that gives excellent fluorescence scanning results: overnight on array incubation with a dibenzocyclooctyne (DBCO)-biotin reagent, followed by the detection reagent Alexa Fluor 647-conjugated streptavidin. This allows visualization and semi-quantitative measurement by the microarray scanner of the glycan probes attached to the array surface.

Here a ‘dose-response’ array of seven FAA-glycan probes was generated on CodeLink (CL) slides to investigate the relationship between concentrations of printed probes (acidic and neutral) and the amounts detected on array after blocking and washing steps (**Fig 2**). Four concentrations of each probe were printed and the unreacted NHS groups on the arrayed slide were blocked by overlaying with ethanolamine blocking solution. To determine the optimal concentration of DBCO-biotin for detecting the glycan probes printed at the different concentrations, serially diluted DBCO-biotin was used 500 μM to 1 nM. Florescence intensities, after automatic subtraction of the background fluorescence by the scanner software, are shown as histograms (**Fig 2D**). There was negligible fluorescence at 1nM and 10 nM of DBCO-biotin concentration; very weak fluorescence emerged at 100 nM and increased at 1 μM and 10 μM and to a maximum at 100 μM DBCO-biotin. The fluorescence readouts were lower at 250 μM and 500 μM DBCO-biotin due to the increased background fluorescence automatically subtracted. For each of the seven glycan probes the fluorescence signals with 100 μM glycan probe and 100 μM DBCO-biotin were in the region of 40,000 – 50,000 units indicating that the relative amounts of glycans immobilized, whether acidic or neutral, varied by no more than 20%.

**Figure 2.**
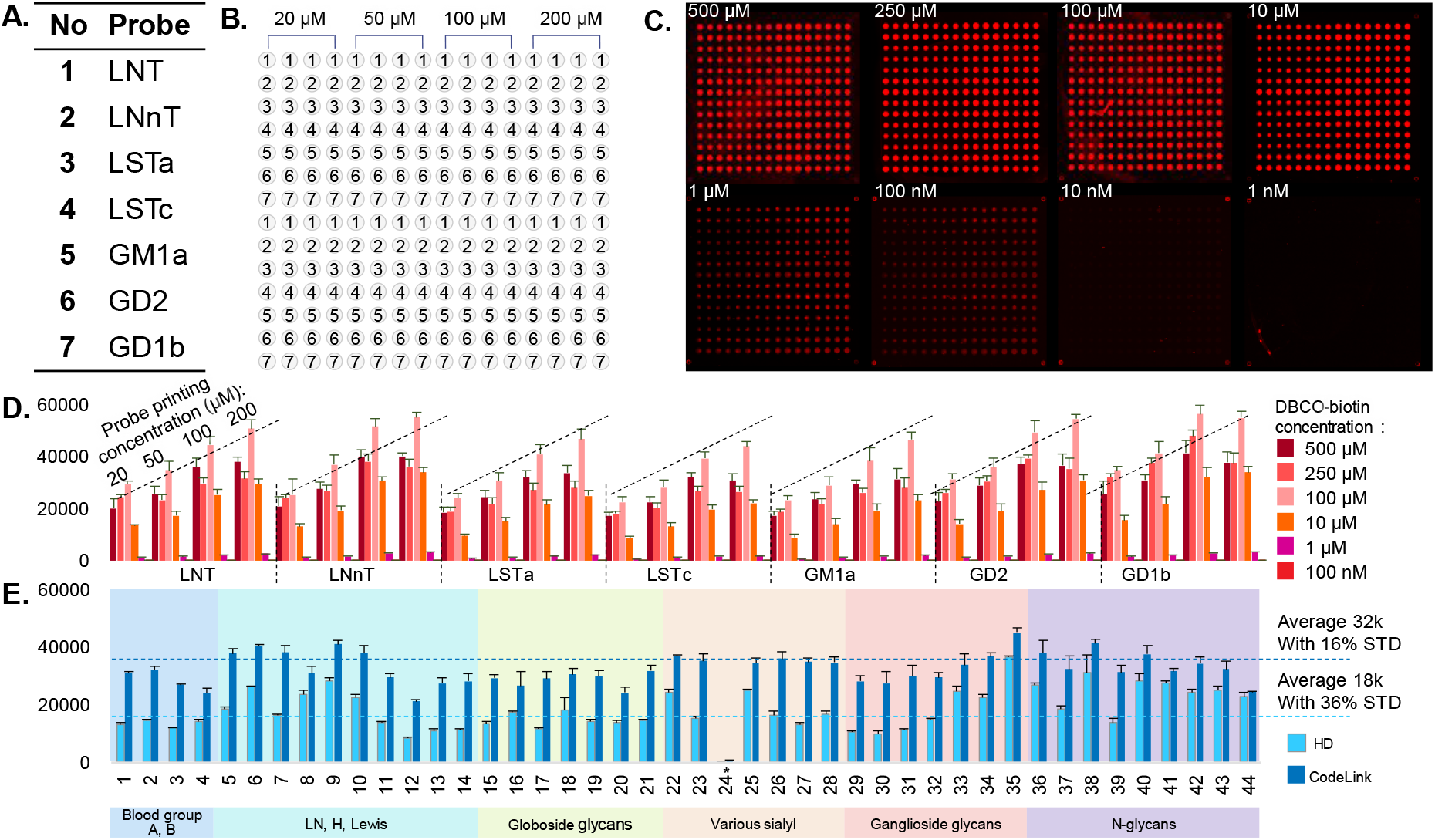
A schema of the ‘on-array’ visualization and quantifying of FAA linked glycan probes. **A)** Seven FAA-linked glycan probes and their designations. The glycan sequences are included in Fig 1C. **B)** Printing layout of the 7 FAA probes. The glycan probes were printed on CL slides at four levels (20 μM to 200 μM) in two rows; thus each probe level was printed as 8 replicates as indicated. **C)** Array images of the 7 FAA-linked probes after on-slide incubation with DBCO-biotin at 1 nM to 500 μM followed by streptavidin Alexa Fluor 647 at 1 μg/mL. **D)** Averages of fluorescence intensities recorded for the 7 arrayed probes after background subtraction. **E)** Averages of fluorescence intensities of the 44 sequence-defined FAA probes arrayed on CodeLink (CL) and high-density Codelink (HD) slides and overlaid with 250 μM DBCO-biotin followed by 1 μg/mL streptavidin Alexa Fluor 647. *Probe 24 was misprinted and did not pass quality control.

### Construction of FAA glycan-based covalent and noncovalent glycan microarrays for cross-platform comparisons

Covalent and NGL-based noncovalent glycan arrays have been widely used for assigning ligands for a diverse range of GBPs^10, 11^. However, systematic comparisons between the two array systems with the same glycan derivatization method have not been carried out so far. This has been the rationale for constructing covalent and noncovalent arrays of FAA based glycan probes to carry out side-by-side comparisons of the performance of the same glycans presented in the two array systems for recognition by different types of GBPs. For the covalent arrays, the density of glycans attached to the functionalized microarray slides is also considered.

#### Covalent array of FAA probes and post-printing ‘on-array’ visualization of arrayed spots

Two covalent glycan arrays were prepared with the 44 FAA glycan probes shown in **Fig 1C** following Fmoc deprotection using the optimal probe printing concentrations determined above (100 μM). Two types of NHS-functionalized glass slides, Codelink (CL) slide and the high-density Codelink (HD) slides were used. This would potentially add additional information on the effect of glycan density on the covalent array surface in glycan recognition. The immobilization efficiency of glycan probes on two types of covalent array slides was assessed using the DBCO-biotin based ‘on-array’ visualization method (**Fig 2E**). The readouts revealed that the various glycan probes were immobilized with a 16% variation on the CL slides and a 36% variation on HD slides. The same volume of printing solution was deposited onto the two types of slides in the same printing cycle. This difference is therefore attributable to the distinct surface properties of the two types of slides: larger and more uniform spots were visualized on the CL slides, and more variable spot sizes on the HD slides.

Of note is the printing of the probe #24 (LSTd) which was depicted as satisfactory by the automatic optical dispense test of arrayer instrument (**Fig S16**). However, the ‘on-array’ visualization method revealed that there was only a negligible amount of this probe immobilized on the array surface (**Fig 2E**). This information is annotated in the subsequent microarray binding results to avoid ‘false negative’ results of this probe in the covalent arrays. This highlights the importance of having the post-printing ‘on-array’ probe visualization and quantitation step as part of the array quality control process.

#### Noncovalent array of FAA-linked glycan probes converted into NGLs

By exploiting the presence of the azido functional group, the FAA-linked glycan probes were converted to NGLs by a single SPAAC reaction step using the lipid reagent DBCO-DHPE (**Scheme S2**). An aliquot (10-20 nmol) each of the 44 FAA glycan probes (**Fig 1C**) were converted into the corresponding FAA-NGL probes, which were purified by HPLC and their molecular weights confirmed by MALDI-MS (**Fig S6**). The resulting NGL probes were quantified based on their lipid content^34^, and arrayed in a liposomal formulation with carrier lipids using the protocol established for conventional NGLs^23^. As standards 11 natural glycolipids (glycosylceramides) were arrayed with the NGLs in arrays designated concovalent set (**Excel Table 2**). The fluorescent dye Cy3 was included in the printing solution of the NGL probes as a tracer to help monitor the printing process^23^.

### Comparisons of the readouts from FAA glycan-based covalent and noncovalent glycan microarrays

A range of glycan recognition systems including 14 anti-glycan antibodies, 9 plant lectins, the adhesins of 3 viruses, 3 bacterial toxins, and 7 lectin-type receptors of the immune system were investigated in a comparative microarray study with the noncovalent on nitrocellulose coated slides and covalent glycan arrays generated on CL and HD slides. The binding signals are detailed in **Figs 3**-**5** and **Figs S11**-**14** in the Supplementary Information. Fluorescence intensities and heat maps of the relative binding strengths of glycans with all of the GBPs investigated are in the **Supplementary Excel Tables 3** and **4**.

**Figure 3.**
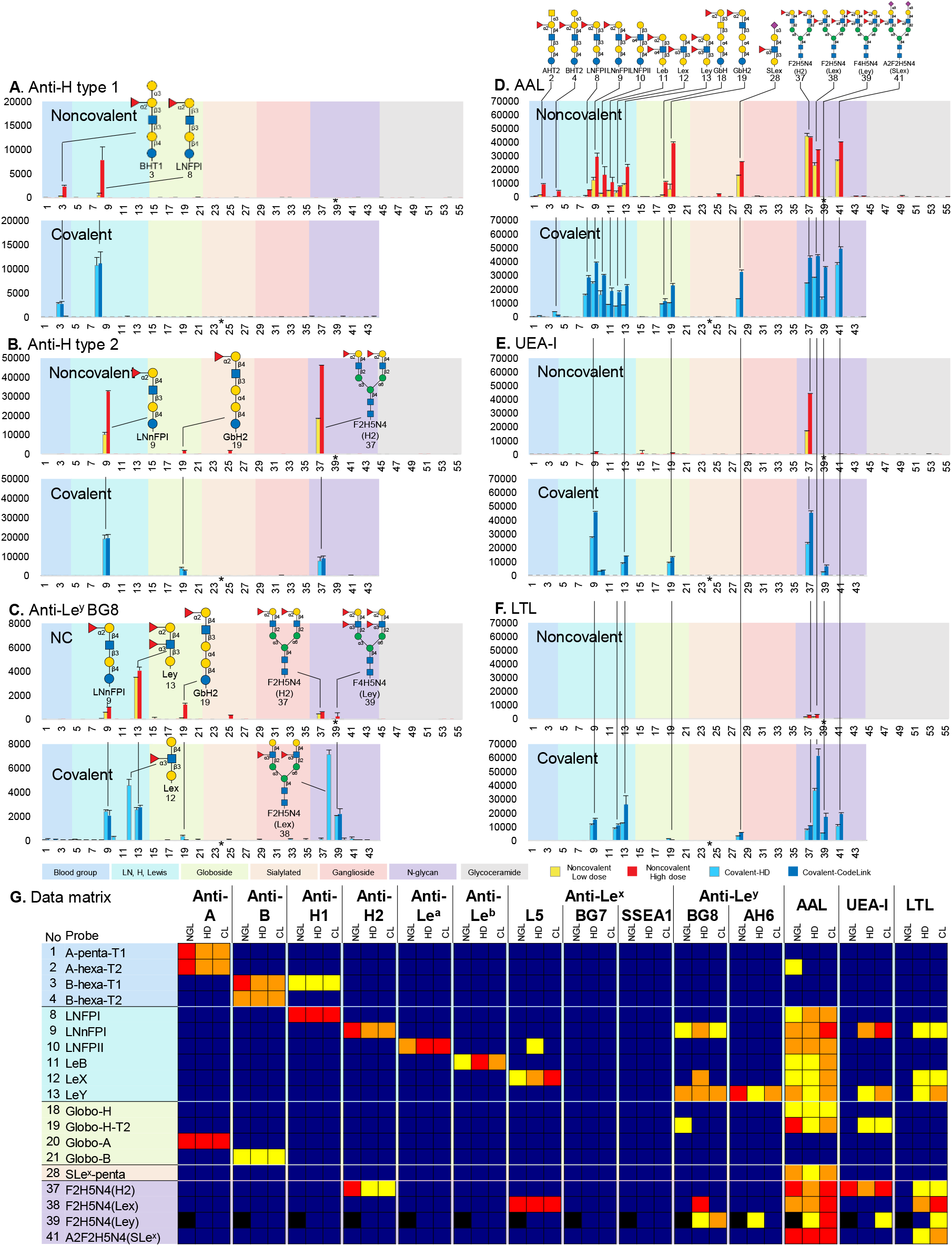
Glycan binding studies with sequence-defined noncovalent and covalent (CL and HD) arrays using anti-blood group A, B, H and Lewis type antibodies, and fucose-binding lectins. Histogram charts showing fluorescence intensities of binding of **A)** anti-H type 1, **B)** anti-H type 2, **C)** anti-Le^y^ BG8, **D)** AAL, **E)** UEA-I, and **F)** LTL. Results are shown as background-subtracted fluorescence intensities of duplicate spots, printed at 2 and 5 fmol per spot level in the noncovalent arrays (error bars represent half of the difference between the two values), and of the averaged fluorescence intensities of quadruplicate spots in the covalent arrays (error bars represent the SD among the four values). **G)** Heatmap comparing the relative binding intensities of the fucosylated glycan probes with the 14 proteins investigated in the three types of FAA arrays: covalent high dose, HD and CL. The highest signal intensity given by each protein across the three types of arrays is 100%. Red >70%, orange 30-70%, yellow 10-30%, blue <10%. * Black, probe 24 on the covalent arrays and probe 39 on the noncovalent array were misprinted and did not pass quality control.

**Figure 4.**
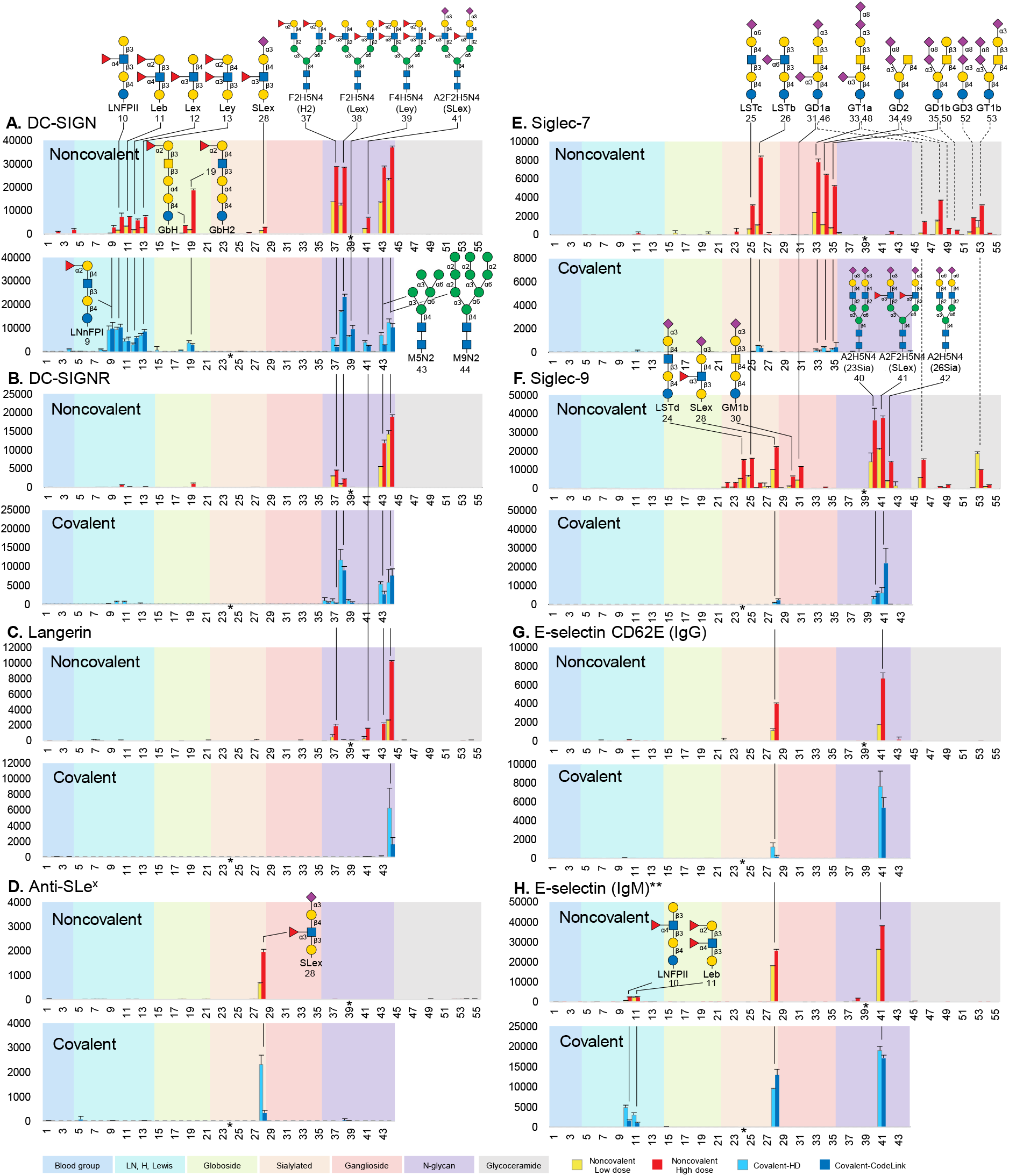
Distinct binding preferences of the innate immune receptors observed in FAA glycan arrays. Microarray analysis of **A)** human Fc-tagged human DC-SIGN, **B)** human DC-SIGNR, **C)** rhesus Langerin, **D)** anti-SLe^x^, **E)** human Siglec-7, **F)** human Siglec-9, **G)** His-tagged human E-selectin, and **H)** murine E-selectin IgM chimera. Results are shown as histogram charts of the background-subtracted fluorescence intensities of duplicate spots, printed at 2 and 5 fmol per spot level in the noncovalent arrays (error bars represent half of the difference between the two values), and of the averaged fluorescence intensities of quadruplicate spots in the covalent arrays (error bars represent the SD among the four values). * Probe 24 on the covalent arrays and probe 39 on the noncovalent array were misprinted and did not pass quality control.

**Figure 5.**
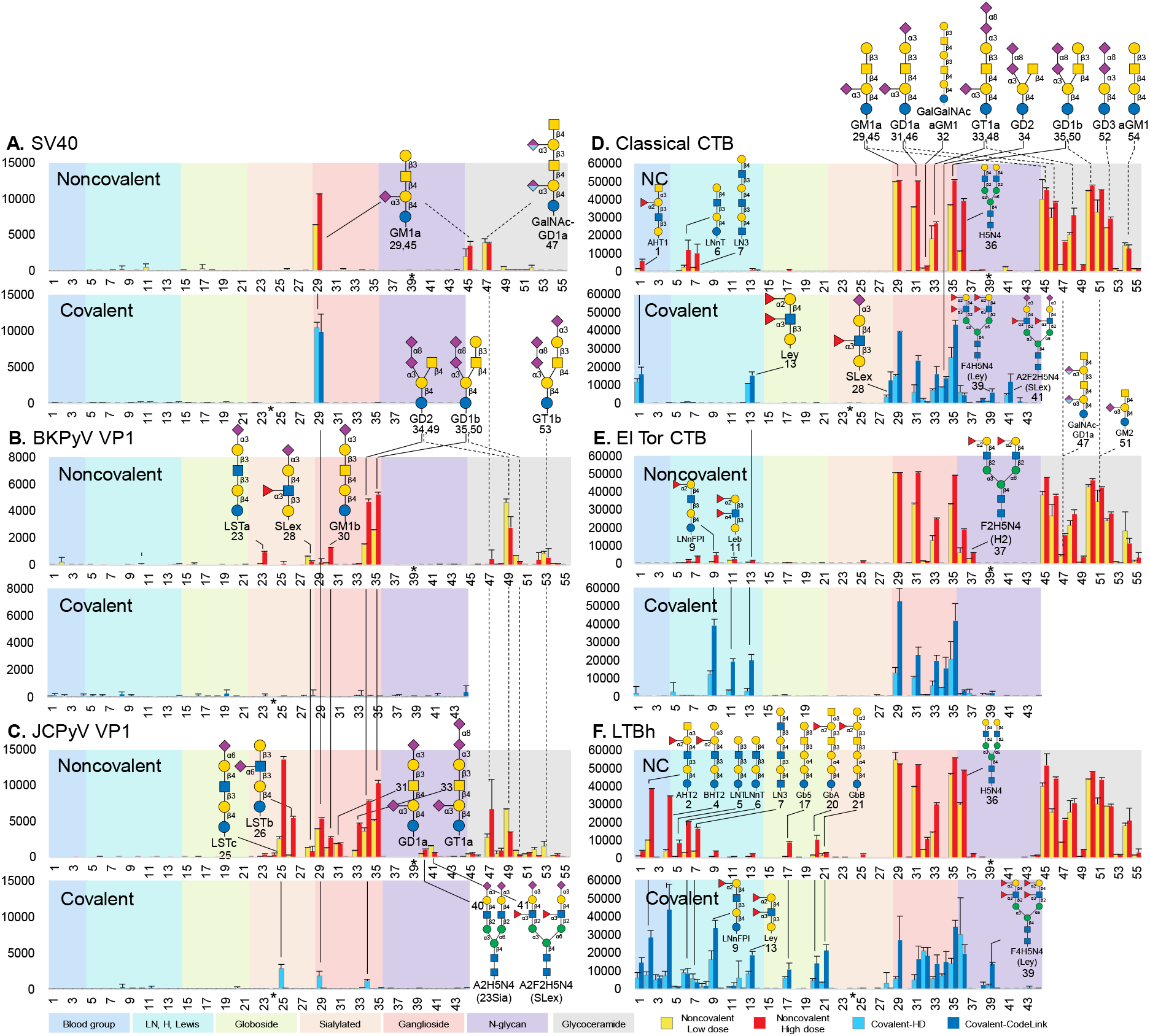
Preferential binding of polyomavirus VPLs in the noncovalent arrays and different glycan binding characteristics revealed for bacterial toxins. Histogram charts showing fluorescence intensities of binding of **A)** his-tagged SV40 VP1, **B)** BKPyV VP1, **C)** JCPyV VP1, **D)** classical CBT, **E)** EI Tor CTB and **F**) LTBh. Results are shown as the background-subtracted fluorescence intensities of duplicate spots, printed at 2 and 5 fmol per spot level in the noncovalent arrays (error bars represent half of the difference between the two values), and of the averaged fluorescence intensities of quadruplicate spots in the covalent arrays (error bars represent the SD among the four values). * Probe 24 on the covalent arrays and probe 39 on the NGL array were misprinted and did not pass quality control.

#### Binding signals given by fucosylated glycans using antibodies and lectins

Fucose-containing glycans are displayed on glycoproteins and glycolipids at the surface of many cell types and their secretions associated, as well as secreted mucins. They are integral parts of the blood group antigens and have important roles in cell recognition and signaling, tumour metastasis and host-pathogen interactions. Lectins and sequence-specific antibodies and have for long been used as histochemical tools as micro-sequencing reagents^35, 36^. Glycan array analyses are a powerful means of assigning and refining knowledge of the specificities of these reagents as more and more sequence defined glycans become available for inclusion in microarrays.

A panel of eleven blood group-related antibodies and three commonly used fucose-binding lectins, *Aleuria Aurantia* Lectin (AAL), *Ulex Europaeus* Agglutinin I (UEA-I) and *Lotus Tetragonolobus* Lectin (LTL), are among the lectins analyzed here using the three types of FAA glycan arrays. This is not only for the purposes of quality control of the 19 fucosylated FAA glycan probes included in the arrays (**Fig 1C**), but also for a better understanding of the fine specificities of the antibodies and lectins towards differing fucosyl linkages and backbone sequences, e.g. type 1, type 2, globo-series, and *N*-glycans.

Overall, the antibodies analyzed recognized their respective glycan antigens as predicted, with comparable signal strengths observed on the noncovalent and covalent CL and HD arrays. Observations are in accord with current knowledge on the proteins investigated. Detailed discussions are in the Supplementary Information. Highlights of observations worthy of specific mention are describes below. The **anti-H type 1** antibody (clone 17-206) could tolerate terminal α-Gal and bound to the blood group B-hexa type 1 probe (#3, **Fig 3A, Fig S10B**). The **anti-H type 2** antibody BE2 bound strongly to the H type 2 pentasaccharide LNnFPI (#9) and biantennary *N*-glycan F2H5N4 (#37) but not the H type 2 globo-sequence GbH2 (#19, **Fig 3B**), indicating the importance of backbone sequence for binding by this antibody.

Of the three **anti-Le**^**x**^ antibodies, only anti-L5 bound to the two sequences (#12, #38) that display Le^x^ motif on a trisaccharide backbone (**Fig S10E**). The BG7 (clone P12) resembles anti-SSEA-1^35^ which was known to require a longer backbone^6^; neither antibody gave binding signals in the covalent or noncovalent FAA glycan arrays (**Fig 3G, Fig S10G&H**). Weak binding of anti-L5 to Le^a^ probe LNFPII (#10) (**Fig 3G, Fig S10E**) was detected as previously noted in the original chromatogram binding study^37^.

The commercial **anti-Le**^**y**^, BG8 (clone F3) gave binding signals with H type 2 probe LNnFPI (#9, **Fig 3C**), in contrast to the anti-Le^y^ monoclonal antibody AH6 from Henrik Clausen Lab^38^, which showed binding exclusively to Le^y^ related probes (#13 and #39, **Fig S11F**). Worthy of note, in the covalent HD array only, the commercial anti-Le^y^ BG8 also gave strong binding to the two Le^x^ probes (#12 and #38) in the absence of the blood group H fucose. Thus the high glycan density of the HD covalent array may increase the likelihood of detecting binding to ‘weak’ binders, reminiscent of findings from an earlier study on anti-HIV antibody recognition^39^.

**AAL** showed binding to blood group H glycans as well as to a broad spectrum of fucosylated glycans. however a lack of significant binding to blood group A and B related sequences was observed in all three types of arrays (**Fig 3D**) This is in accord with information provided in the recently published lectin guide^40^ and data from our conventional NGL-based screening arrays (unpublished).

**UEA-I**, known for its specificity towards α1-2-fucosylated type 2 backbone glycans, indeed showed binding to α1-2-Fuc-terminating biantennary *N*-glycan in both covalent and noncovalent arrays (#37, **Fig 3E**). However, LNnFPI (#9) which elicited strong binding with the anti-H type 2 antibody BE2 in all three arrays (**Fig 3B**), was bound by UEA-I only on the covalent arrays rather than the noncovalent array.

**LTL**, which is known to have 1-3-fucose specificity, showed good binding to the Le^x^ and Le^y^ related probes in the covalent arrays but not in the noncovalent array (**Fig 3F**). Knowing that the Le^x^ and Le^y^ NGL probes were well arrayed and strongly bound by the anti-Le^x^ and Le^y^ antibodies respectively (**Fig 3G**), our findings suggest that these two glycan probes are more favourably displayed to LTL on the covalent rather than the noncovalent array. LTL binding detected here to the sialyl Le^x^ on a short SLe^x^ or the antennae of *N*-glycan (#28, #41) and H type 2 probe LNnFPI (#9) has not been described^40^ to our knowledge and this would be an addition to knowledge on the specificity assignment of this lectin.

Microarray data with other antibodies and lectins analyzed are in the Supplementary Information (Figs **S12-S14**). It is noteworthy that, while the binding signals with ConA were similar in the covalent and noncovalent arrays, several other lectins either exhibited higher overall signal intensities in the covalent arrays, as with PNA, SNA and MAA-I, or showed fine differences in the binding to certain glycan probes, as with RCA120 and WGA (**Supplementary Excel Table 3 and Table S7**). Further details regarding the observations of glycan binding specificities of these GBPs are in the **Supplementary Results and Discussion section**.

#### Binding signals given by innate immune receptors

Using the FAA glycan arrays, we investigated the binding preferences of seven receptors of the innate immune system. These included the closely related C-type lectin receptors (CLRs) DC-SIGN, DC-SIGNR, Langerin, and two siglecs, Siglec-7 and Siglec-9. These are immune regulatory receptors playing important roles in immune cell signalling and modulation^41^. In addition, we analyzed an IgM and a His-tagged construct of E-selectin, an adhesion molecule that mediates the interaction between leukocytes and endothelial cells and during inflammation through binding to sialylated and fucosylated glycans on including sialyl Le^x^ on leukocytes^42^.

### CLRs

Overall with the three CLRs the signal-to-noise ratio was better in the noncovalent compared to the covalent arrays (**Fig 4A-C**). **DC-SIGN** in common with the other two CLRs is known to bind to oligomannose glycans^43, 44^. In addition, it binds to fucosyl glycans including H type 2 and Le^a,b,x and y 45^. Under the three assay conditions, the top binders for DC-SIGN in non-covalent array were oligomannose *N*-glycans (#43, #44) and biantennary H type 2 (#37) and Le^x^-terminating complex type *N*-glycans (#37, #38), followed by H type 2 globo-sequence GbH2 (#19) and the Lewis type antigens (#10-#13). The preference for the oligomannose *N*-glycan probes was less prominent in the covalent arrays (**Fig 4A**). **DC-SIGNR** showed a more restricted binding profile towards *N*-glycan probes including oligomannose (#43, #44) *N*-glycans with all three arrays. Notably, DC-SIGNR exhibited strong binding to the Le^x^-*N*-glycan (#38) in the covalent arrays, whereas only weak binding was observed in the noncovalent array (**Fig 4B**). To our knowledge, this finding represents novel evidence for the binding of DC-SIGNR to glycans beyond oligo/high-mannose *N*-glycans^43^. **Langerin** bound almost exclusively to the high-mannose Man9GN2 probe in three types of FAA arrays. The binding intensity was considerably weaker in the CL array than the HD and noncovalent arrays (**Fig 4C**). The lack of binding to Man5GN2 highlights the requirement of α1,2-linked Man for binding by this CLR, in accord with the recent findings using an oligomannose isomer-focused array^44^.

### Siglecs

With the present probe library FAA glycans arrayed, the two human siglecs showed binding signals mainly in the noncovalent arrays but not the covalent arrays (**Fig 4E** and **F**). **Siglec-7** showed binding to α2,6-sialyl sequence LSTb (#26) and to α2,8-sialylated GT1a, GD2, GD1b (#33-35) in agreement with findings from previous studies^46, 47^. These sequences were included in all three arrays. All of these probes and in addition LSTc (#25) have given binding signals in the noncovalen array, whereas little or no binding was observed in covalent arrays. **Siglec-9** is known to bind to α2,3- and α2,6-linked sialyl sequences including sialyl Le^x^ (SLe^x^) containing probes. There was little overlap with the binding profile of Siglec-7. Except for binding to SLe^x^ *N*-glycan probe (#41), binding to α2,3-or α2,6-sialylated probes (#24-26) were mainly detected in the noncovalent and not the covalent arrays. This contrasts with the binding signals detected with the sialic acid-binding plant lectins SNA and MAA-I which showed stronger binding in the covalent than the noncovalent arrays (**Fig S13**).

### E-selectin

The high avidity IgM construct of murine E-selectin gave strong binding both with the covalent and noncovalent arrays to the potent SLe^x^ terminating probes (#28, #41, **Fig 4H**) and relatively weak binding to asialo-Le^a^, Le^b^ probes (#10, #11) which are low affinity ligands for E-selectin; this is in accord with previous observation in cell based analyses^48^. The His-tagged human E-selectin gave binding signals with the two SLe^x^ probes (#28, #41, **Fig 4G**) but not with the Le^a^ and Le^b^ probes. In contrast, the **anti-SLe**^**x**^ antibody (anti-human CD15S, clone CSLEX1) gave good binding exclusively to the SLe^x^ pentasaccharide (#28) but not the biantennary *N*-glycan probe (#41) in all three types of arrays (**Fig 4D**). The hindrance observed with the biantennary *N*-glycan probe is in reminiscent of observation with a similar anti-SLe^x^ (anti-human CD15S, clone CHO131)^49^.

#### Interactions of microbial adhesins and toxins with their ligands on the different forms of FAA glycan arrays

The interactions between microorganisms and host cells often involve the recognition of specific host glycans by microbial proteins. These include adhesive proteins that mediate the initial host cell attachments, and toxins produced by various pathogenic bacteria which are virulence factors perturbing host physiological processes. Here we have investigated the adhesive VP1 proteins of three polyomaviruses and toxins of three bacteria. One common characteristic among these six proteins is their recognition of sialyl glycans, in particular gangliosides, which facilitates cellular entry or uptake^50^.

##### Polyomavirus VP1 proteins

Previous studies with the conventional NGL arrays have identified specific sialyl glycan ligands for the VP1 proteins of simian virus 40 (SV40)^51^, human JC polyomavirus (JCPyV)^52^, and human BK polyomavirus (BKPyV)^53^. In the present study, **SV40 VP1** showed strong binding to the well-recognized ligand^51^, the ganglioside GM1a in the form of FAA glycan probe (#29) in all three arrays (**Fig 5A**). In the noncovalent array where glycosylceramides were included, binding was detected to the natural GM1a (#45). There was also binding to disialyl glycolipid, GalNAc-GD1a with one Neu5Ac and one Neu5Gc (#47)^54^; the latter was not included in our previous microarray study^51^.

###### BKPyV and JCPyV

There was a striking difference in readouts in the covalent and noncovalent arrays, namely the detection of strong binding in the noncovalent and little or no binding in the covalent arrays (**Fig 5B** and **C**). Here the **BKPyV** VP1 bound mainly to the GD2 and GD1b probes, consistent with the previous study^55^. In an earlier NGL-based array^52^ the **JCPyV** VP1 protein was shown to bind exclusively to the L-shaped α2-6-sialyl sequence LSTc. Here, using the FAA-noncovalent array with our improved liposomal formulation and protein construct, we confirm the binding the LSTc probe (#25) with the strongest signal intensity, but we also detect interactions with various other sialyl glycan probes, in particular gangliosides, GM1a, GD2 and GD1b. These recent results align closely with a comprehensive structural and functional study indicating the VP1s of different JCPyV strains engage multiple gangliosides in addition to LSTc^56^.

##### Bacterial toxins

Three structurally related toxins of the cholera toxin family: *Vibrio cholerae* Toxin Subunit B of the O1 classical biotype (Classical CTB, cCTB) and the El Tor biotype (El Tor CTB), and the enterotoxigenic *Escherichia coli* heat-labile Toxin (LTBh)^57^ were investigated here. In contrast to the findings with BKPyV and JCPyV VP1s, the covalent and the noncovalent arrays exhibited robust binding (**Fig 5D-F**). All three toxins bound strongly to a wide range of ganglioside probes with similar binding patterns in the three types of arrays. Although ganglioside GM1a is known to be the most potent ligand for the cholera toxins^58^, when the toxins were overlaid at 5 μg/mL, the binding signals detected with GM1a, GD1a, GD1b were almost equally strong in the noncovalent arrays, followed by GT1a (#33), GD3 (#53) and asialo-GM1 (#55). The ganglioside binding was also well detected in the covalent CL arrays but less strongly in the HD arrays. There was a lack of binding of the three toxins to GM1b probes (#30) in the three arrays. These findings are in overall agreement with those from earlier reports^59, 60^. An observation of note is the binding to GD2 detectable in the covalent arrays (#34) but not in the noncovalent arrays (#34, #49). When the toxin concentration was reduced to 0.5 μg/mL, a noteworthy decrease in signal intensity was observed in the covalent arrays, amounting to approximately one-fourth of that observed in the noncovalent arrays, suggesting that the noncovalent platform exhibits a higher sensitivity for studying toxin binding.

With the three types of FAA arrays binding of the three toxins could be detected to non-sialyl glycans^61-64^, Greater differences were observed in the different array formats that with the ganglioside related sequences. For example, binding to Le^y^ probe (#13), the known ligand for the three toxins^57^, was well detected in the covalent CL array but only weakly in the HD arrays and almost absent in the noncovalent array. In contrast, the LacNAc-terminating glycans were more prominently bound in the noncovalent array compared to the covalent arrays: the complex type *N*-glycan (#36) was bound by all three toxins, and LNnT (#5) and LN3 (#6) by the cCTB and LTBh. The binding of the three CTB-related toxins to blood group antigens was observed with varying intensities in the three arrays and these are described in detail as below.

**cCTB**, binding was observed to blood group A type 1 (#1) in both covalent and noncovalent arrays. This has not been reported to our knowledge. In the covalent arrays, additional binding was detected to the two Le^y^ terminating (#13, #39) and two SLe^x^ glycan probes (#28, #41) in agreement with the results obtained from inhibition of cell binding studies using soluble oligosaccharides^65^. **El Tor CTB** showed binding exclusively to H type 2 probe LNnFPI (#9) among all the ABO antigens tested. This finding provides an explanation for the blood group O association with the severity of cholera caused by the El

Tor biotype of *V. cholerae*^66^. This toxin also bound to α1-2-fucosylated Lewis antigens Le^b^ (#11) and Le^y^ (#13), particularly with the CL array. **LTBh** is reported to bind to blood group A and B but not H-oligosaccharides^67^. In this study, we were able to detect LTBh binding to H-oligosaccharides in addition to A- and B-(#2, #4, #9), mostly on type 2 sequences. However, little binding was detected to the N-glycan probe with short type 2 H outer arms (#37) and Globo-H analog GbH2 (#19) having the H-type 2 epitope. LTBh also bound well to Globopentaose Gb5 (#17), Globo-A and B probes (#20, #21), but not Globo-H (#18).

The differential binding of the three toxins to blood group antigens can be summarized as follows. **cCTB:** binds A type 1, Le^y^, SLe^x^. **El Tor CTB** binds: H type 2, Le^y^, Le^b^. **LTBh** binds A and B (type 2 more strongly than type 1), also A and B on globo-backbone, H type 2, Le^y^. Thus, it appears that the bacteria have evolved to produce toxins with differential preferences to blood group related glycans and their backbones, the majority of which are found in the intestine.

### Rendering the FAA glycan probes fluorescent for cell recognition assays

To broaden the applications of FAA tagged glycan probes, we have incorporated a red Fluor 545 dye into the probes to render them suitable for fluorescence-based cell binding assays. By a simple SPACC click reaction, the DBCO-PEG4-Fluor 545 (DBCO-F545) reagent has been coupled with the azido group of the FAA glycan probes with high efficiency (**Fig 6A**). This has allowed facile assembly of a panel of seven F545-FAA glycan probes: four sialyl glycans (SLe^x^, LSTc, LSTd and LSTb) as well as three neutral glycans (LNnT, Mal3, and Mal5). The preparation of these F545-labelled probes and their characterization are in Supplementary Information.

**Figure 6.**
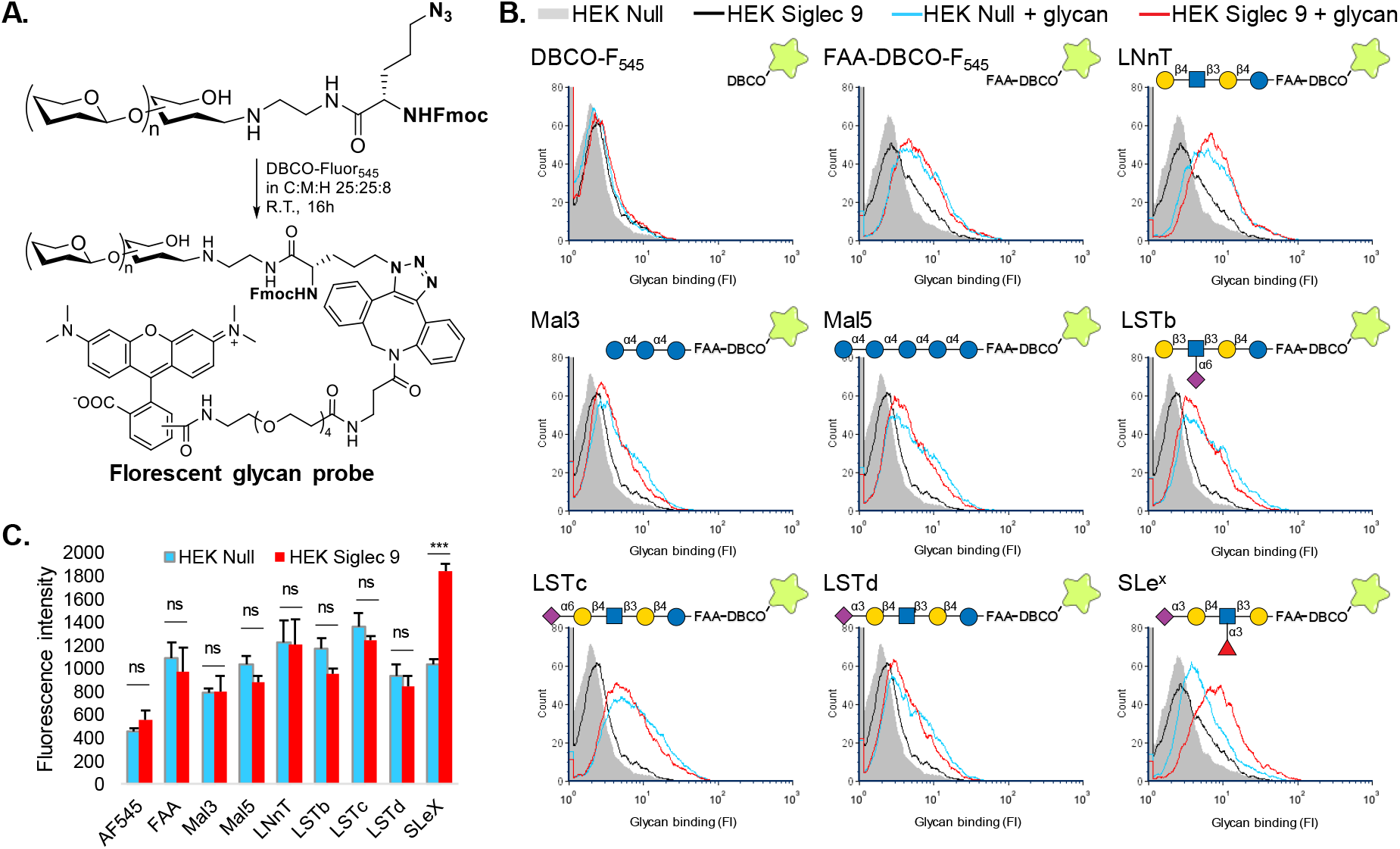
Analysis of HEK Siglec-9 cells incubated with fluorescently labelled glycans. **A)** Scheme for fluorescent labelling of FAA tagged glycan probes. **B)** Flow cytometry-based glycan binding profiles of HEK Siglec-9 and control cell line HEK Null1. **C)** Fluorescence intensities of the HEK Siglec-9 and HEK Null1 cells incubated with the fluorescently labelled glycans in the microwell experiments (mean +/-SD, n=3 independent experiments, ***: p< 0,001).

Cell binding analyses were performed with the fluorescent glycan probes using HEK cells transfected with Siglec-9 and HEK null1 cells as controls. The F545-FAA-glycan probes, the unconjugated dye reagent DBCO-F545 and the F545 labelled blank FAA linker (without glycan) FAA-F545 were also used as controls. The results of the flow cytometry analysis and the corresponding microwell-plate fluorescence readouts are in **Fig 6B and C**. Whereas little or no interaction was observed with DBCO-F545, the FAA-F545 control showed some ‘background’ binding of comparable intensity to both HEK Null1 and HEK Siglec-9 cells. This binding is likely attributable to hydrophobic interactions stemming from the FAA linker.

Cell binding profiles similar to those of FAA-F545 tag were observed with the F545-FAA-glycans tested with the exception of the high affinity Siglec-9 ligand SLe^x^, with which a clear shift in fluorescence intensity was observed with the HEK Siglec-9 cells compared to the control HEK Null1 cells (**Fig 6B**). The microwell assay gave results comparable to those of the flow cytometry read out, a significantly higher binding to the Siglec 9-expressing cells being observed only with the SLe^x^ probe (**Fig 6C**). There was no shift in cell fluorescence intensity with the F545-FAA glycans LSTc and LSTd which are weaker ligands for Siglec-9 than SLe^x^, as observed in the binding experiments with Siglec 9 using noncovalent glycan arrays (**Figure 4F**). The fluorescent glycans are in monomeric state in solution thus only with the high affinity ligand could binding be detected to the Siglec-9 expressing cells.

## Conclusions

Over the past two decades, glycan microarrays have emerged as essential tools in glycoscience, with technological advancements and increasing applications in laboratories worldwide. Glycan microarrays have greatly accelerated the acquisition of data on glycan-mediated interactions. However, interpreting data obtained from various array platforms is not always straightforward, as multiple variables exist throughout glycan microarray experiments^68^, potentially yielding differing results even with the same glycan binding system. The ‘glycan array community’ is recognizing the importance of adhering to the minimal information requirements outlined in the MIRAGE Guidelines for reporting glycan microarray analysis data^69^ to ensure the generation of interpretable data. Nonetheless, significant questions persist regarding the performance of different platforms raising the possibility of standardizing glycan array methodologies.

The newly developed trifunctional FAA linker described in this study imparts functionalities to native glycans whereby they can be immobilized on microarrays by different means, covalently on NHS-functionalized slides or noncovalently as lipid-liked probes in a liposomal formulation on nitrocellulose-coated slides. The ‘on-array’ visualization of the FAA glycan probes via the ‘spare’ azido functional group in the FAA linker, represents a significant advantage in the quality control of arrayed glycans. This approach enables the accurate assessment of immobilized glycans on arrays and mitigates false negative results.

A focus of this study has been to closely compare the presentation of 44 FAA-linked glycans on the covalent and noncovalent arrays, for binding by 36 GBP samples using identical assay conditions and data analysis procedures. Salient observations here on glycan presentation on the different array platforms are that glycan binding signals similar or equivalent for the majority - 18 out of 36 GBPs investigated (**Table S7**). There are however striking differences: plant lectin LTL showed binding in the covalent arrays only; in contrast, with 2 of the 7 lectins of the innate immune system, Siglec-7 and Siglec-9, and 2 of the 3 viral adhesins, VP1s of polyomaviruses JCPyV and BKPyV, binding signals were detected with the noncovalent arrays only. Subtle but significant differences were also observed with the remaining GBPs tested, including the three structurally related toxins which showed similarly strong binding to ganglioside related probes in the covalent and non-covalent arrays but distinct binding preferences with the low affinity glycan ligands such as blood group and galactose-terminating sequences between the different types of arrays. The preference for distinct glycan ligands on different array platforms is likely influenced by the specific requirements regarding ligand density and presentation, necessary for binding by multivalent binding sites of GBPs. Observations worthy of note for users of the GBPs as research tools are highlighted in **Table S7**.

Thus, the lack of binding signals on a particular platform does not necessarily indicate the absence of the glycan ligand; rather, the presentation of the ligand on that platform may not be conducive to favourable binding with the GBP partner. In assessing the specificities of binding by different GBPs, it is advantageous to record data from both covalent and noncovalent array platforms. The FAA glycan probe strategy goes a considerable way to increase confidence in the readouts. A special feature of lipid-linked glycan probes is their ability to be incorporated into cells to evaluate the cell biological relevance of binding by adhesins of infective agents, as was shown previously using conventional NGLs in SV40 studies^70^. Similar studies can be envisaged for the FAA-NGLs as a follow on of binding signals with viral adhesins and bacterial toxins.

Overall, the FAA strategy shows promise for future designs of comprehensive screening arrays to encompass both noncovalent and covalent arrays of glycans, as well as probes for cell binding assays. Moreover, the detection limit of the FAA-linked glycans at low pmol, renders the approach suitable for labelling natural glycans that are often available in limited in amounts.

The ease of conversion of FAA-tagged glycans into fluorescent probes renders them promising candidates for binding assays with cells in suspension. Our proof-of-concept data have shown distinct binding signals between Siglec-9 expressing cells and the fluorescently labelled high affinity ligand SLe^x^-FAA probe. Once the conditions are optimised, for example by rendering the glycans multimeric, the way would be open to use fluorescent FAA-based glycan probes to survey cell surfaces for binding of glycans with a range of avidities for GBPs, by flow cytometry or other fluorescence-based techniques such as microscopy or plate-based screening assays.

## Acknowledgements

This work was supported by the Wellcome Trust Biomedical Resource Grants (WT099197/Z/12/Z, 108430/Z/15/Z and 218304/Z/19/Z) and the MRC grant (MR/R010757/1). The March of Dimes Prematurity Research Center grant 22-FY18-82 provided partial financial support to the Imperial College Glycosciences Laboratory.

## References

(1) Varki, A. Biological roles of glycans. Glycobiology 2017, 27 (1), 3–49. DOI: 10.1093/glycob/cww086

(2) Taylor, M. E.; Drickamer, K. Mammalian sugar-binding receptors: known functions and unexplored roles. The FEBS Journal 2019, 286 (10), 1800–1814. DOI: https://doi.org/10.1111/febs.14759

(3) Thompson, A. J.; de Vries, R. P.; Paulson, J. C. Virus recognition of glycan receptors. Curr Opin Virol 2019, 34, 117–129. DOI: 10.1016/j.coviro.2019.01.004

(4) Ballal, S.; Inamdar, S. R. An overview of lectin-glycan interactions: a key event in initiating fungal infection and pathogenesis. Arch Microbiol 2018, 200 (3), 371–382. DOI: 10.1007/s00203-018-1487-1

(5) Poole, J.; Day, C. J.; von Itzstein, M.; Paton, J. C.; Jennings, M. P. Glycointeractions in bacterial pathogenesis. Nat Rev Microbiol 2018, 16 (7), 440–452. DOI: 10.1038/s41579-018-0007-2

(6) Rodrigues, J. A.; Acosta-Serrano, A.; Aebi, M.; Ferguson, M. A.; Routier, F. H.; Schiller, I.; Soares, S.; Spencer, D.; Titz, A.; Wilson, I. B.; et al. Parasite Glycobiology: A Bittersweet Symphony. PLoS Pathog 2015, 11 (11), e1005169. DOI: 10.1371/journal.ppat.1005169

(7) Wesener, D. A.; Dugan, A.; Kiessling, L. L. Recognition of microbial glycans by soluble human lectins. Curr Opin Struct Biol 2017, 44, 168–178. DOI: 10.1016/j.sbi.2017.04.002

(8) Antolín-Llovera, M.; Petutsching, E. K.; Ried, M. K.; Lipka, V.; Nürnberger, T.; Robatzek, S.; Parniske, M. Knowing your friends and foes – plant receptor-like kinases as initiators of symbiosis or defence. New Phytologist 2014, 204 (4), 791–802. DOI: https://doi.org/10.1111/nph.13117

(9) Fukui, S.; Feizi, T.; Galustian, C.; Lawson, A. M.; Chai, W. Oligosaccharide microarrays for high-throughput detection and specificity assignments of carbohydrate-protein interactions. Nat Biotechnol 2002, 20 (10), 1011–1017. DOI: 10.1038/nbt735

(10) Palma, A. S.; Feizi, T.; Childs, R. A.; Chai, W.; Liu, Y. The neoglycolipid (NGL)-based oligosaccharide microarray system poised to decipher the meta-glycome. Curr Opin Chem Biol 2014, 18, 87–94. DOI: 10.1016/j.cbpa.2014.01.007

(11) Rillahan, C. D.; Paulson, J. C. Glycan microarrays for decoding the glycome. Annu Rev Biochem 2011, 80, 797–823. DOI: 10.1146/annurev-biochem-061809-152236

(12) Pardo-Vargas, A.; Delbianco, M.; Seeberger, P. H. Automated glycan assembly as an enabling technology. Curr Opin Chem Biol 2018, 46, 48–55. DOI: 10.1016/j.cbpa.2018.04.007

(13) Na, L.; Li, R.; Chen, X. Recent progress in synthesis of carbohydrates with sugar nucleotide-dependent glycosyltransferases. Curr Opin Chem Biol 2021, 61, 81–95. DOI: 10.1016/j.cbpa.2020.10.007

(14) Li, T.; Liu, L.; Wei, N.; Yang, J. Y.; Chapla, D. G.; Moremen, K. W.; Boons, G. J. An automated platform for the enzyme-mediated assembly of complex oligosaccharides. Nat Chem 2019, 11 (3), 229–236. DOI: 10.1038/s41557-019-0219-8

(15) Ruprecht, C.; Blaukopf, M.; Pfrengle, F. Synthetic fragments of plant polysaccharides as tools for cell wall biology. Curr Opin Chem Biol 2022, 71, 102208. DOI: 10.1016/j.cbpa.2022.102208

(16) Song, X.; Ju, H.; Lasanajak, Y.; Kudelka, M. R.; Smith, D. F.; Cummings, R. D. Oxidative release of natural glycans for functional glycomics. Nat Methods 2016, 13 (6), 528–534. DOI: 10.1038/nmeth.3861

(17) Li, Z.; Zhang, Q.; Ashline, D.; Zhu, Y.; Lasanajak, Y.; Chernova, T.; Reinhold, V.; Cummings, R. D.; Wang, P. G.; Ju, T.; et al. Amplification and Preparation of Cellular O-Glycomes for Functional Glycomics. Anal Chem 2020, 92 (15), 10390–10401. DOI: 10.1021/acs.analchem.0c00632

(18) Schallus, T.; Jaeckh, C.; Fehér, K.; Palma, A. S.; Liu, Y.; Simpson, J. C.; Mackeen, M.; Stier, G.; Gibson, T. J.; Feizi, T.; et al. Malectin: A Novel Carbohydrate-binding Protein of the Endoplasmic Reticulum and a Candidate Player in the Early Steps of Protein N-Glycosylation. Molecular Biology of the Cell 2008, 19 (8), 3404–3414. DOI: 10.1091/mbc.e08-04-0354

(19) Liu, Y.; Palma, A. S.; Feizi, T. Carbohydrate microarrays: key developments in glycobiology. Biol Chem 2009, 390 (7), 647–656. DOI: 10.1515/bc.2009.071

(20) Prudden, A. R.; Chinoy, Z. S.; Wolfert, M. A.; Boons, G. J. A multifunctional anomeric linker for the chemoenzymatic synthesis of complex oligosaccharides. Chem Commun (Camb) 2014, 50 (54), 7132–7135. DOI: 10.1039/c4cc02222j

(21) Song, X.; Heimburg-Molinaro, J.; Smith, D. F.; Cummings, R. D. Glycan microarrays of fluorescently-tagged natural glycans. Glycoconj J 2015, 32 (7), 465–473. DOI: 10.1007/s10719-015-9584-8

(22) Wei, M.; McKitrick, T. R.; Mehta, A. Y.; Gao, C.; Jia, N.; McQuillan, A. M.; Heimburg-Molinaro, J.; Sun, L.; Cummings, R. D. Novel Reversible Fluorescent Glycan Linker for Functional Glycomics. Bioconjug Chem 2019, 30 (11), 2897–2908. DOI: 10.1021/acs.bioconjchem.9b00613

(23) Liu, Y.; Childs, R. A.; Palma, A. S.; Campanero-Rhodes, M. A.; Stoll, M. S.; Chai, W.; Feizi, T. Neoglycolipid-Based Oligosaccharide Microarray System: Preparation of NGLs and Their Noncovalent Immobilization on Nitrocellulose-Coated Glass Slides for Microarray Analyses. In Carbohydrate Microarrays: Methods and Protocols, Chevolot, Y. Ed.; Humana Press, 2012; pp 117–136.

(24) Gao, C.; Wei, M.; McKitrick, T. R.; McQuillan, A. M.; Heimburg-Molinaro, J.; Cummings, R. D. Glycan Microarrays as Chemical Tools for Identifying Glycan Recognition by Immune Proteins. Front Chem 2019, 7, 833. DOI: 10.3389/fchem.2019.00833

(25) Blixt, O.; Head, S.; Mondala, T.; Scanlan, C.; Huflejt, M. E.; Alvarez, R.; Bryan, M. C.; Fazio, F.; Calarese, D.; Stevens, J.; et al. Printed covalent glycan array for ligand profiling of diverse glycan binding proteins. Proc Natl Acad Sci U S A 2004, 101 (49), 17033–17038. DOI: 10.1073/pnas.0407902101

(26) Grant, O. C.; Smith, H. M.; Firsova, D.; Fadda, E.; Woods, R. J. Presentation, presentation, presentation! Molecular-level insight into linker effects on glycan array screening data. Glycobiology 2014, 24 (1), 17–25. DOI: 10.1093/glycob/cwt083

(27) Temme, J. S.; Campbell, C. T.; Gildersleeve, J. C. Factors contributing to variability of glycan microarray binding profiles. Faraday Discuss 2019, 219 (0), 90–111. DOI: 10.1039/c9fd00021f

(28) Padler-Karavani, V.; Song, X.; Yu, H.; Hurtado-Ziola, N.; Huang, S.; Muthana, S.; Chokhawala, H. A.; Cheng, J.; Verhagen, A.; Langereis, M. A.; et al. Cross-comparison of protein recognition of sialic acid diversity on two novel sialoglycan microarrays. J Biol Chem 2012, 287 (27), 22593–22608. DOI: 10.1074/jbc.M112.359323

(29) Wang, L.; Cummings, R. D.; Smith, D. F.; Huflejt, M.; Campbell, C. T.; Gildersleeve, J. C.; Gerlach, J. Q.; Kilcoyne, M.; Joshi, L.; Serna, S.; et al. Cross-platform comparison of glycan microarray formats. Glycobiology 2014, 24 (6), 507–517. DOI: 10.1093/glycob/cwu019

(30) Li, C.; Palma, A. S.; Zhang, P.; Zhang, Y.; Gao, C.; Silva, L. M.; Li, Z.; Trovao, F.; Weishaupt, M.; Seeberger, P. H.; et al. Noncovalent microarrays from synthetic amino-terminating glycans: Implications in expanding glycan microarray diversity and platform comparison. Glycobiology 2021, 31 (8), 931–946. DOI: 10.1093/glycob/cwab037

(31) Klamer, Z. L.; Harris, C. M.; Beirne, J. M.; Kelly, J. E.; Zhang, J.; Haab, B. B. CarboGrove: a resource of glycan-binding specificities through analyzed glycan-array datasets from all platforms. Glycobiology 2022, 32 (8), 679–690. DOI: 10.1093/glycob/cwac022

(32) Bigge, J. C.; Patel, T. P.; Bruce, J. A.; Goulding, P. N.; Charles, S. M.; Parekh, R. B. Nonselective and Efficient Fluorescent Labeling of Glycans Using 2-Amino Benzamide and Anthranilic Acid. Analytical Biochemistry 1995, 230 (2), 229–238. DOI: https://doi.org/10.1006/abio.1995.1468

(33) Song, X.; Xia, B.; Stowell, S. R.; Lasanajak, Y.; Smith, D. F.; Cummings, R. D. Novel fluorescent glycan microarray strategy reveals ligands for galectins. Chemistry & biology 2009, 16 1, 36–47.

(34) Chai, W.; Stoll, M. S.; Galustian, C.; Lawson, A. M.; Feizi, T. Neoglycolipid Technology: Deciphering Information Content of Glycome. In Methods in Enzymology, Vol. 362; Academic Press, 2003; pp 160–195.

(35) Gooi, H. C.; Feizi, T.; Kapadia, A.; Knowles, B. B.; Solter, D.; Evans, M. J. Stage-specific embryonic antigen involves αl→ 3 fucosylated type 2 blood group chains. Nature 1981, 292 (5819), 156–158. DOI: 10.1038/292156a0

(36) Kapadia, A.; Feizi, T.; Evans, M. J. Changes in the expression and polarization of blood group I and I antigens in post-implantation embryos and teratocarcinomas of mouse associated with cell differentiation. Experimental Cell Research 1981, 131 (1), 185–195. DOI: https://doi.org/10.1016/0014-4827(81)90418-3

(37) Streit, A.; Yuen, C.-T.; Loveless, R. W.; Lawson, A. M.; Finne, J.; Schmitz, B.; Feizi, T.; Stern, C. D. The Lex Carbohydrate Sequence Is Recognized by Antibody to L5, a Functional Antigen in Early Neural Development. Journal of Neurochemistry 1996, 66 (2), 834–844. DOI: https://doi.org/10.1046/j.1471-4159.1996.66020834.x

(38) Young, W. W.; Portoukalian, J.; Hakomori, S. Two monoclonal anticarbohydrate antibodies directed to glycosphingolipids with a lacto-N-glycosyl type II chain. Journal of Biological Chemistry 1981, 256 (21), 10967–10972. DOI: https://doi.org/10.1016/S0021-9258(19)68541-8

(39) Kong, L.; Lee, J. H.; Doores, K. J.; Murin, C. D.; Julien, J.-P.; McBride, R.; Liu, Y.; Marozsan, A.; Cupo, A.; Klasse, P.-J.; et al. Supersite of immune vulnerability on the glycosylated face of HIV-1 envelope glycoprotein gp120. Nature Structural & Molecular Biology 2013, 20 (7), 796–803. DOI: 10.1038/nsmb.2594

(40) Bojar, D.; Meche, L.; Meng, G.; Eng, W.; Smith, D. F.; Cummings, R. D.; Mahal, L. K. A Useful Guide to Lectin Binding: Machine-Learning Directed Annotation of 57 Unique Lectin Specificities. ACS Chemical Biology 2022, 17 (11), 2993–3012. DOI: 10.1021/acschembio.1c00689

(41) Macauley, M. S.; Crocker, P. R.; Paulson, J. C. Siglec-mediated regulation of immune cell function in disease. Nature Reviews Immunology 2014, 14 (10), 653–666. DOI: 10.1038/nri3737

(42) Barthel, S. R.; Gavino, J. D.; Descheny, L.; Dimitroff, C. J. Targeting selectins and selectin ligands in inflammation and cancer. Expert Opinion on Therapeutic Targets 2007, 11 (11), 1473–1491. DOI: 10.1517/14728222.11.11.1473

(43) Guo, Y.; Feinberg, H.; Conroy, E.; Mitchell, D. A.; Alvarez, R.; Blixt, O.; Taylor, M. E.; Weis, W. I.; Drickamer, K. Structural basis for distinct ligand-binding and targeting properties of the receptors DC-SIGN and DC-SIGNR. Nature Structural & Molecular Biology 2004, 11 (7), 591–598. DOI: 10.1038/nsmb784

(44) Gao, C.; Stavenhagen, K.; Eckmair, B.; McKitrick, T. R.; Mehta, A. Y.; Matsumoto, Y.; McQuillan, A. M.; Hanes, M. S.; Eris, D.; Baker, K. J.; et al. Differential recognition of oligomannose isomers by glycan-binding proteins involved in innate and adaptive immunity. Science Advances 2021, 7 (24), eabf6834. DOI: doi:10.1126/sciadv.abf6834

(45) van Liempt, E.; Bank, C. M. C.; Mehta, P.; García-Vallejo, J. J.; Kawar, Z. S.; Geyer, R.; Alvarez, R. A.; Cummings, R. D.; Kooyk, Y. v.; van Die, I. Specificity of DC-SIGN for mannose- and fucose-containing glycans. FEBS Letters 2006, 580 (26), 6123–6131. DOI: https://doi.org/10.1016/j.febslet.2006.10.009

(46) Yamaji, T.; Teranishi, T.; Alphey, M. S.; Crocker, P. R.; Hashimoto, Y. A Small Region of the Natural Killer Cell Receptor, Siglec-7, Is Responsible for Its Preferred Binding to α2,8-Disialyl and Branched α2,6-Sialyl Residues: A COMPARISON WITH Siglec-9 *. Journal of Biological Chemistry 2002, 277 (8), 6324–6332. DOI: 10.1074/jbc.M110146200 (acccessed 2023/05/29).

(47) Campanero-Rhodes, M. A.; Childs, R. A.; Kiso, M.; Komba, S.; Le Narvor, C.; Warren, J.; Otto, D.; Crocker, P. R.; Feizi, T. Carbohydrate microarrays reveal sulphation as a modulator of siglec binding. Biochemical and Biophysical Research Communications 2006, 344 (4), 1141–1146. DOI: https://doi.org/10.1016/j.bbrc.2006.03.223

(48) Larkin, M.; Ahern, T. J.; Stoll, M. S.; Shaffer, M.; Sako, D.; O’Brien, J.; Yuen, C. T.; Lawson, A. M.; Childs, R. A.; Barone, K. M. Spectrum of sialylated and nonsialylated fuco-oligosaccharides bound by the endothelial-leukocyte adhesion molecule E-selectin. Dependence of the carbohydrate binding activity on E-selectin density. Journal of Biological Chemistry 1992, 267 (19), 13661–13668. DOI: 10.1016/S0021-9258(18)42264-8 (acccessed 2023/06/14).

(49) Li, L.; Guan, W.; Zhang, G.; Wu, Z.; Yu, H.; Chen, X.; Wang, P. G. Microarray analyses of closely related glycoforms reveal different accessibilities of glycan determinants on N-glycan branches. Glycobiology 2019, 30 (5), 334–345. DOI: 10.1093/glycob/cwz100 (acccessed 6/14/2023).

(50) O’Hara, S. D.; Stehle, T.; Garcea, R. Glycan receptors of the Polyomaviridae: structure, function, and pathogenesis. Current Opinion in Virology 2014, 7, 73–78. DOI: https://doi.org/10.1016/j.coviro.2014.05.004

(51) Campanero-Rhodes, M. A.; Smith, A.; Chai, W.; Sonnino, S.; Mauri, L.; Childs, R. A.; Zhang, Y.; Ewers, H.; Helenius, A.; Imberty, A.; et al. <i>N</i>-Glycolyl GM1 Ganglioside as a Receptor for Simian Virus 40. Journal of Virology 2007, 81 (23), 12846–12858. DOI: doi:10.1128/JVI.01311-07

(52) Neu, U.; Maginnis, M. S.; Palma, A. S.; Ströh, L. J.; Nelson, C. D. S.; Feizi, T.; Atwood, W. J.; Stehle, T. Structure-Function Analysis of the Human JC Polyomavirus Establishes the LSTc Pentasaccharide as a Functional Receptor Motif. Cell Host & Microbe 2010, 8 (4), 309–319. DOI: 10.1016/j.chom.2010.09.004 (acccessed 2023/05/29).

(53) Neu, U.; Allen, S.-a. A.; Blaum, B. S.; Liu, Y.; Frank, M.; Palma, A. S.; Ströh, L. J.; Feizi, T.; Peters, T.; Atwood, W. J.; et al. A Structure-Guided Mutation in the Major Capsid Protein Retargets BK Polyomavirus. PLOS Pathogens 2013, 9 (10), e1003688. DOI: 10.1371/journal.ppat.1003688

(54) Casellato, R.; Brocca, P.; Li, S.-C.; Li, Y.-T.; Sonnino, S. Isolation and Structural Characterization of N-Acetyl- and N-Glycolylneuraminic-Acid-Containing GalNAc-GD1a Isomers, IV4GalNAcIV3Neu5AcII3Neu5GcGgOse4Cer and IV4GalNAcIV3Neu5AcII3Neu5AcGgOse4Cer, from Bovine Brain. European Journal of Biochemistry 1995, 234 (3), 786–793. DOI: https://doi.org/10.1111/j.1432-1033.1995.786_a.x

(55) Sorin, M. N.; Di Maio, A.; Silva, L. M.; Ebert, D.; Delannoy, C. P.; Nguyen, N.-K.; Guerardel, Y.; Chai, W.; Halary, F.; Renaudin-Autain, K.; et al. Structural and functional analysis of natural capsid variants suggests sialic acid-independent entry of BK polyomavirus. Cell Reports 2023, 42 (2). DOI: 10.1016/j.celrep.2023.112114 (acccessed 2023/05/29).

(56) Ströh, L. J.; Maginnis, M. S.; Blaum, B. S.; Nelson, C. D. S.; Neu, U.; Gee, G. V.; O’Hara, B. A.; Motamedi, N.; DiMaio, D.; Atwood, W. J.; et al. The Greater Affinity of JC Polyomavirus Capsid for α2,6-Linked Lactoseries Tetrasaccharide c than for Other Sialylated Glycans Is a Major Determinant of Infectivity. Journal of Virology 2015, 89 (12), 6364–6375. DOI: doi:10.1128/jvi.00489-15

(57) Mandal, P. K.; Branson, T. R.; Hayes, E. D.; Ross, J. F.; Gavín, J. A.; Daranas, A. H.; Turnbull, W. B. Towards a Structural Basis for the Relationship Between Blood Group and the Severity of El Tor Cholera. Angewandte Chemie International Edition 2012, 51 (21), 5143–5146. DOI: https://doi.org/10.1002/anie.201109068

(58) Holmgren, J.; Lönnroth, I.; Månsson, J.; Svennerholm, L. Interaction of cholera toxin and membrane GM1 ganglioside of small intestine. Proceedings of the National Academy of Sciences 1975, 72 (7), 2520–2524. DOI: doi:10.1073/pnas.72.7.2520

(59) Angström, J.; Teneberg, S.; Karlsson, K. A. Delineation and comparison of ganglioside-binding epitopes for the toxins of Vibrio cholerae, Escherichia coli, and Clostridium tetani: evidence for overlapping epitopes. Proceedings of the National Academy of Sciences 1994, 91 (25), 11859–11863. DOI: doi:10.1073/pnas.91.25.11859

(60) Kuziemko, G. M.; Stroh, M.; Stevens, R. C. Cholera Toxin Binding Affinity and Specificity for Gangliosides Determined by Surface Plasmon Resonance. Biochemistry 1996, 35 (20), 6375–6384. DOI: 10.1021/bi952314i

(61) Holmner, A.; Askarieh, G.; Okvist, M.; Krengel, U. Blood group antigen recognition by Escherichia coli heat-labile enterotoxin. J Mol Biol 2007, 371 (3), 754–764. DOI: 10.1016/j.jmb.2007.05.064

(62) Mandal, P. K.; Branson, T. R.; Hayes, E. D.; Ross, J. F.; Gavin, J. A.; Daranas, A. H.; Turnbull, W. B. Towards a structural basis for the relationship between blood group and the severity of El Tor cholera. Angew Chem Int Ed Engl 2012, 51 (21), 5143–5146. DOI: 10.1002/anie.201109068

(63) Heggelund, J. E.; Burschowsky, D.; Bjornestad, V. A.; Hodnik, V.; Anderluh, G.; Krengel, U. High-Resolution Crystal Structures Elucidate the Molecular Basis of Cholera Blood Group Dependence. PLoS Pathog 2016, 12 (4), e1005567. DOI: 10.1371/journal.ppat.1005567

(64) Heim, J. B.; Hodnik, V.; Heggelund, J. E.; Anderluh, G.; Krengel, U. Crystal structures of cholera toxin in complex with fucosylated receptors point to importance of secondary binding site. Sci Rep 2019, 9 (1), 12243. DOI: 10.1038/s41598-019-48579-2

(65) Wands, A. M.; Cervin, J.; Huang, H.; Zhang, Y.; Youn, G.; Brautigam, C. A.; Matson Dzebo, M.; Björklund, P.; Wallenius, V.; Bright, D. K.; et al. Fucosylated Molecules Competitively Interfere with Cholera Toxin Binding to Host Cells. ACS Infectious Diseases 2018, 4 (5), 758–770. DOI: 10.1021/acsinfecdis.7b00085

(66) Harris, J. B.; Khan, A. I.; LaRocque, R. C.; Dorer, D. J.; Chowdhury, F.; Faruque, A. S. G.; Sack, D. A.; Ryan, E. T.; Qadri, F.; Calderwood, S. B. Blood Group, Immunity, and Risk of Infection with <i>Vibrio cholerae</i> in an Area of Endemicity. Infection and Immunity 2005, 73 (11), 7422–7427. DOI: doi:10.1128/iai.73.11.7422-7427.2005

(67) Barra, J. L.; Monferran, C. G.; Balanzino, L. E.; Cumar, F. A. Escherichia coli heat-labile enterotoxin preferentially interacts with blood group A-active glycolipids from pig intestinal mucosa and A- and B-active glycolipids from human red cells compared to H-active glycolipids. Molecular and Cellular Biochemistry 1992, 115 (1), 63–70. DOI: 10.1007/BF00229097

(68) Temme, J. S.; Campbell, C. T.; Gildersleeve, J. C. Factors contributing to variability of glycan microarray binding profiles. Faraday Discussions 2019, 219 (0), 90–111, 10.1039/C9FD00021F. DOI: 10.1039/C9FD00021F

(69) Liu, Y.; McBride, R.; Stoll, M.; Palma, A. S.; Silva, L.; Agravat, S.; Aoki-Kinoshita, K. F.; Campbell, M. P.; Costello, C. E.; Dell, A.; et al. The minimum information required for a glycomics experiment (MIRAGE) project: improving the standards for reporting glycan microarray-based data. Glycobiology 2017, 27 (4), 280–284. DOI: 10.1093/glycob/cww118

(70) Ewers, H.; Romer, W.; Smith, A. E.; Bacia, K.; Dmitrieff, S.; Chai, W.; Mancini, R.; Kartenbeck, J.; Chambon, V.; Berland, L.; et al. GM1 structure determines SV40-induced membrane invagination and infection. Nat Cell Biol 2010, 12 (1), 11–18; sup pp 11-12. DOI: 10.1038/ncb1999

