## Supporting Info for "Novel multifunctional glycan probes to elucidate specificities of diverse glycan recognition systems"

- Supporting Information

MATERIALS AND METHODS

General methods

Commercial chemicals and solvents were of the highest grade available. All chemicals were from Merck (Dorset, UK) unless otherwise stated. Thin layer chromatography (TLC) was performed on Silica Gel F254 (Merck, Dorset, UK) with detection by UV-light. High performance TLC (HPTLC) for glycolipids (GLs) and neoglycolipids (NGLs) was performed on Silica Gel plates (Merck, Dorset, UK). For HPTLC analyses, glycans, GLs and NGLs were applied onto the silica gel plates with a TLC applicator (Camag Linomat 5, Muttenz, Switzerland) and the TLC tank was equilibrated with the developing solvents (as indicated) for 30 min prior to chromatography. Lipids were detected by UV-light after staining with primulin solution^1^ and glycans were visualized by staining with orcinol solution^1^. The quantitation based on readouts from the chromatogram plates was performed using a Camag TLC Scanner 3. Silica and C18 cartridges used for the purification of glycans were purchased from WATERS (Manchester, UK). Cellulose powder (10-100 μm particle size) used for column chromatography was from Sigma. Flash column chromatography was performed on silica gel 60 Å (35-70 mm, Acros, Geel, Belgium). ^1^H-NMR, as well as 2D HSQC and HMBC NMR spectra were recorded at 303 K with a Bruker Avance III HD 500 MHz NMR spectrometer using CDCl_3_ (internal CDCl_3_, δH 7.26; δC 77.16 ppm at 303 K) or Methanol-d4 (internal MeOD-D_4_, δH 4.87; δC 49.13 ppm at 303 K). HPLC purification of derivatized glycans was performed on a WATERS E2695 system with 2489 UV and 2475 fluorescence detectors. For normal phase (NP) HPLC, an Amide HPLC column (3.5 μm, 4.6 mm x 250 mm, Xbridge, WATERS) was used. High resolution MS spectra (HR-ESI-MS) were recorded with a Waters Synapt G2-S mass spectrometer. HPLC fractions were analysed on a Shimadzu Axima Resonance MALDI QIT-TOF mass spectrometer (Milton Keynes, Buckinghamshire, UK).

Abbreviations of chemicals: Hexafluorophosphate azabenzotriazole tetramethyl uronium (HATU), 1-Hydroxy-7-azabenzotriazole (HOAt), Dimethylformamide (DMF), N,N-Diisopropylethylamine (DIPEA), Dibenzocyclooctyne-N-hydroxysuccinimidyl ester (DBCO-NHS), 1,2-Dihexadecyl-*sn*-glycero-3-phosphoethanolamine (DHPE) and Dibenzocyclooctyne-PEG4-Fluor 545 (DBCO-PEG4-Flour545).

Glycan materials

*Sequence defined glycans and glycosylceramides*

Forty-four sequence-defined glycans were purchased from Elicityl (Crolles, France) and Dextra (Reading, UK), as indicated in Table S1. The glycan sequences are given in Excel Table 1. In addition, eleven glycosylceramides from the collection from Glycosciences Laboratory are given in Excel Table 2 in the Excel data sheet included

*Glycans released from proteins*

Ribonuclease B (RNaseB) was purchased from TCI chemicals (Tokyo, Japan). Fetuin was from Merck. Endoglycosidase H (EndoH) and peptide *N*-glycanse F (PNGF) were from New England Biolabs (Manchester, UK). Oligomannose *N*-glycans were released from RNaseB (40 mg) using EndoH according to the recommended procedure from the manufacturer (<https://international.neb.com/protocols/2018/04/05/protocol-for-endo-h-p0702-and-p0703>). Complex type *N*-glycans were released from fetuin (40 mg) using PNGF according to the recommended procedure from the manufacturer ([https://international.neb.com/protocols
/2014/07/31/pngase-f-protocol](https://international.neb.com/protocols/2014/07/31/pngase-f-protocol)).

2-6Sialylated biantennary *N*-glycan was released from sialylglycopeptides (TCI chemicals, Tokyo, Japan) using PNGF as described^2^.

Synthesis of the FAA linker

**(9*H*-fluoren-9-yl)methyl (*S*)-(5-azido-1-((2-((*tert*-butoxycarbonyl) amino)ethyl)amino)-1-oxopentan-2-yl)carbamate** (Compound **1**)

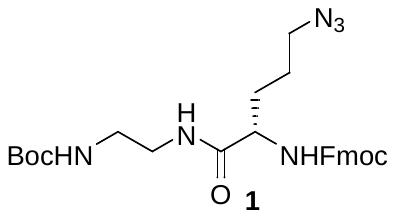

To a solution of Fmoc-L-azidoornithine (3.0 g, 7.9 μmol, 1 eq) with HATU (3.0 g, 7.9 mmol, 1 eq) and HOAt (1.07 g, 7.9 mmol, 1 eq) in DMF (20 mL), DIPEA (2.47 mL, 15.8 mmol, 2 eq) was added. The activated amino acid mixture was then slowly added to a solution of N-Boc-ethylenediamine (1.87 mL, 11.8 mmol, 1.5 eq) in DMF (2 mL). After incubating at room temperature (R.T.) for 2 h, TLC indicated a full conversion of the starting material Fmoc-L-azidoornithine into a single product. The reaction mixture was then diluted with 50 mL CHCl_3_ and extracted three times with 50 mL H_2_O. The residue was concentrated under reduced pressure and purified by flash column chromatography (CHCl_3_:MeOH 99:1 to 80:20) to give compound **1** in 81% yield (3.34 g, 6.4 umol). R_f_ = 0.4 (cyclohexane:ethyl acetate:acetic acid 3:2:0.01). The MS and NMR spectra are in Figure S1 and Figure S3. ***HR-ESI-MS***, *m/z*: 545.2593 ([M+Na]^+^, calc. 545.2483). ***^1^H-NMR*** (500 MHz, Chloroform-*d*) δ = 7.75 (d, *J*=7.6, 2H, H-4-; H-5-Fmoc), 7.57 (dd, *J*=7.5, 2.7, 2H, H-1-; H-8-Fmoc), 7.38 (td, *J*=7.5, 3.2, 2H, H-2-; H-7-Fmoc), 7.32 – 7.26 (m, 2H, H-3-; H-6-Fmoc), 7.05 (s, 1H, (CH_2_)_2_N*H*CO), 5.77 (d, *J*=8.1, 1H, Fmoc-N*H*), 5.11 (t, *J*=6.3, 1H, Boc-N*H*), 4.43 (dd, *J*=10.6, 7.1, 1H, Fmoc C*H*_2_), 4.33 (dd, *J*=10.6, 7.0, 1H, Fmoc C*H*_2_), 4.19 (m, 2H, C*H*α, H-9-Fmoc), 3.38 – 3.05 (m, 6H, BocNHCH_2_C*H*_2_, C*H*_2_δ, BocNHC*H*_2_CH_2_), 1.90 (p, *J*=6.2, 1H, C*H*_2_β), 1.70 (dt, *J*=14.3, 7.4, 1H, C*H*_2_β), 1.61 (p, *J*=7.2, 2H, C*H*_2_γ), 1.41 (s, 9H,C(C*H*_3_)_3_). ***^13^C-NMR*** (126 MHz, CDCl_3_) δ = 172.02 (*C*=OCHα), 156.97 (Boc *C*=O), 156.31(Fmoc *C*=O), 143.84, 143.78 (C-1_a_-; C-8_a_-Fmoc), 141.39 (C-4_a_-; C-5_a_-Fmoc), 127.84 (C-2-; C-7-Fmoc), 127.16 (C-3-; C-6-Fmoc), 125.18, 125.11 (C-1-; C-8-Fmoc), 120.10, 120.08 (C-4-; C-5-Fmoc), 79.85 (*C*(CH_3_)_3_), 67.10 (Fmoc *C*H_2_), 54.49 (*C*Hα), 51.03 (*C*H_2_δ), 47.21 (C-9-Fmoc), 40.78 (BocNHCH_2_*C*H_2_), 40.16 (BocNH*C*H_2_CH_2_), 30.20 (*C*H_2_β), 28.43(C(*C*H_3_)_3_), 24.95 (*C*H_2_γ).

**(9*H*-fluoren-9-yl)methyl (*S*)-(1-((2-aminoethyl)amino)-5-azido-1-oxopentan-2-yl)carbamate (**Compound **2, FAA linker)**

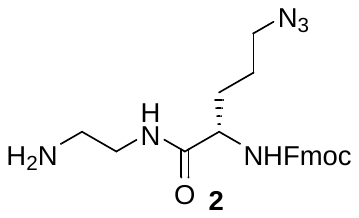

To an Eppendorf tube containing compound **1** (20 mg, 38.3 μmol), 200 μL 2 M HCl in MeOH was added and the vial was vortexed vigorously at R.T. for 2 h until full solubilisation of the starting material. The reaction was monitored to completion by TLC; and the solvent was removed to semi-dryness under reduced pressure, diluted with H_2_O and freeze-dried twice to give compound **2** (FAA linker) as a white powder in quantitative yield. R_f_ = 0.25 (ethyl acetate:acetic acid:MeOH:H_2_O 60:3:3:2). ***HR-ESI-MS***, *m/z*: 423.2335 ([M+H]^+^, calc. 423.2139). The MS and NMR spectra are in Figure S2 and Figure S4. ***^1^H-NMR*** (500 MHz, Methanol-*d*_4_) δ = 7.70 (d, *J*=7.5, 2H, H-4-; H-5-Fmoc), 7.58 (d, *J*=7.4, 2H, H-1-; H-8-Fmoc), 7.30 (t, *J*=7.4, 2H, H-2-; H-7-Fmoc), 7.22 (td, *J*=7.5, 1.1, 2H, H-3-; H-6-Fmoc), 4.39 (dd, *J*=10.6, 6.7, 1H, Fmoc C*H*_2_), 4.27 (dd, *J*=10.7, 6.6, 1H, Fmoc C*H*_2_), 4.13 (t, *J*=6.5, 1H, H-9-Fmoc), 3.95 (dd, *J*=8.9, 5.0, 1H, C*H*α), 3.47 (dt, *J*=14.3, 5.9, 1H, NH_2_CH_2_C*H*_2_), 3.33 (dt, *J*=14.2, 5.8, 1H, BocNHCH_2_C*H*_2_), 3.28 – 3.18 (m, 2H, C*H*_2_δ), 2.97 (m, 2H, NH_2_C*H*_2_CH_2_), 1.84 – 1.73 (m, 1H, C*H*_2_β), 1.63 – 1.45 (m, 3H, C*H*_2_β, C*H*_2_γ). ***^13^C-NMR*** (126 MHz, MeOD) δ = 175.80 (*C*=OCHα), 158.77 (Fmoc *C*=O), 145.22, 145.07 (C-1_a_-; C-8_a_-Fmoc), 142.60 (C-4_a_-; C-5_a_-Fmoc), 128.80 (C-2-; C-7-Fmoc), 128.14 (C-3-; C-6-Fmoc), 126.20, 126.10 (C-1-; C-8-Fmoc), 120.95, 120.93 (C-4-; C-5-Fmoc), 67.92 (Fmoc *C*H_2_), 56.32 (*C*Hα), 51.96 (*C*H_2_δ), 48.38 (C-9-Fmoc), 40.81 (NH_2_*C*H_2_CH_2_), 38.13 (NH_2_CH_2_*C*H_2_), 29.96 (*C*H_2_β), 26.41 (*C*H_2_γ).

Generation of multifunctional glycan probes using FAA linker

A solution containing 0.18 M FAA linker (compound **2**) with 0.5 M NaCNBH_3_ in anhydrous MeOH was added to the lyophilized glycans. Typically, 20 eq FAA linker was used relative to the glycans (1 nmol to 10 µmol). Where the glycans were in trace amounts, a minimum of 10 μL of solution was required. The pH of the solution was adjusted to 6 using a solution of 0.1% w/v NaOMe in MeOH and the reaction mixture was vortexed at 50 ^o^C on a ThermoMixer (Eppendorf ^TM^) for 4 h. Thereafter, the excess reagents were removed using a cellulose cartridge. For a typical glycan (200 nmol) labelling using 2 mg of FAA linker, a cellulose cartridge with a 50 μL bed volume was used. The cartridge was first activated by washing sequentially with 0.3 ml of MeOH, followed by 0.3 mL of MeCN, followed by 0.3 mL of 0.1% TFA in H_2_O, and then equilibrated with 90% MeCN in H_2_O. The reaction mixture was diluted with 10 volumes of MeCN and loaded onto the cartridge. The cartridge was then washed with 1 ml of 90% MeCN in H_2_O to remove excess reagents and the FAA-glycan probes were collected by eluting with 0.5 mL 0.1% TFA in H_2_O, concentrated and subjected to NP HPLC (amide column) for further purification.

Generation of glycan probes for covalent arrays on NHS coated glass slides

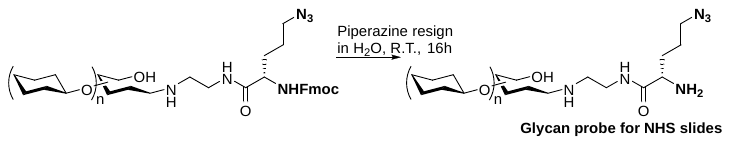

Scheme S1. Generating of the glycan probes for covalent array.

For covalent arrays the Fmoc group was removed from the quantified FAA-tagged glycans. One nmol of each in H_2_O, was added to a plastic Eppendorf vial containing 15 mg piperazine resin (Merck, Dorset, UK) and was topped up with H_2_O to 100 µL. The reaction was kept at R.T. for 16 h and MS analysis indicated full deprotection of the Fmoc group. The resin was separated by filtration and the freshly prepared probe in the filtrate was lyophilized to dryness. Ten µL of 0.1 M phosphate buffer (pH 8.5) was added to the glycan probe, to have the probe at 100 µM.

Generation of NGL probes for printing on nitrocellulose-coated glass slides

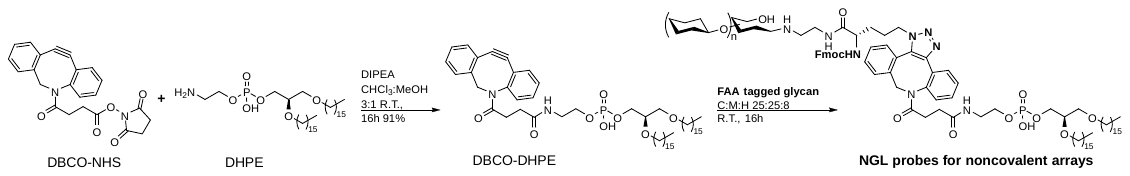

**Scheme S2. Conversion of the FAA tagged glycan into NGLs.**

To a solution of DHPE (3.0 mg, 4.5 μmol, 1 eq) in CHCl_3_:MeOH 2:1 (1.5 mL), DBCO-NHS (3.6 mg, 9 μmol, 2 eq) with DIPEA (3.2 μL, 20 μmol, 4 eq) in CHCl_3_ (0.5 mL) was added dropwise. The reaction mixture was stirred at R.T. for 16 h and followed by HPTLC. The reaction mixture was first extracted three times with 2 mL H_2_O and purified by flash column chromatography (solvent CHCl_3_:MeOH 10:0 to 8.5:1.5) to give DBCO-DHPE in 82% yield (3.5 mg, 3.7 μmol). R_f_ = 0.4 (CHCl_3_:MeOH 9:1). ***HR-ESI-MS***, *m/z*: 949.6464 ([M-H]^-^, calc. 949.6435).

For conversion to NGLs, 10 nmol of dry FAA-tagged glycans were each dissolved in 40 μL CHCl_3_:MeOH:100mM ammonium acetate in H_2_O 25:25:8 containing 40 nmol of DBCO-DHPE. The reaction was kept at R.T. for 16 h and the NGLs purified by NP HPLC with an amide column using CHCl_3_:MeOH:H_2_O (Solvent A 130:70:9 to Solvent B 10:20:8). A short gradient was used here to effectively remove excess DBCO-DHPE. The column was then kept in 95% solvent A for 5 min to elute DBCO-DHPE followed by switching to 90% B for 4 min isocratically to elute the NGL product.

Generation of florescent glycan probes for cell binding studies

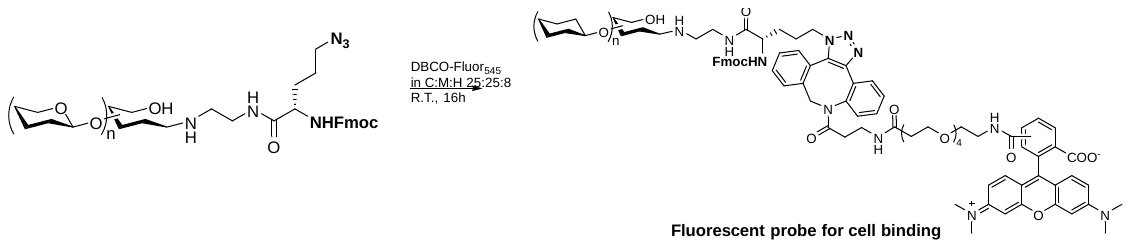

Scheme S3. Florescence tagging of glycan probes.

For florescence labelling, seven FAA tagged glycans were used, which included LSTb, LSTc, LSTd, LNnT, Mal3, Mal5 and SLe^x^. The FAA tagged glycans, 5 nmol, were dissolved in 40 μL CHCl_3_:MeOH:100mM ammonium acetate in H_2_O 25:25:8 containing 20 nmol of DBCO-PEG4-Flour545. The reactions were kept at R.T. for 16 h and MALDI-MS indicated full conversion of the starting material. The solvent was removed under reduced pressure and the reaction residue was used directly in cell incubation experiments.

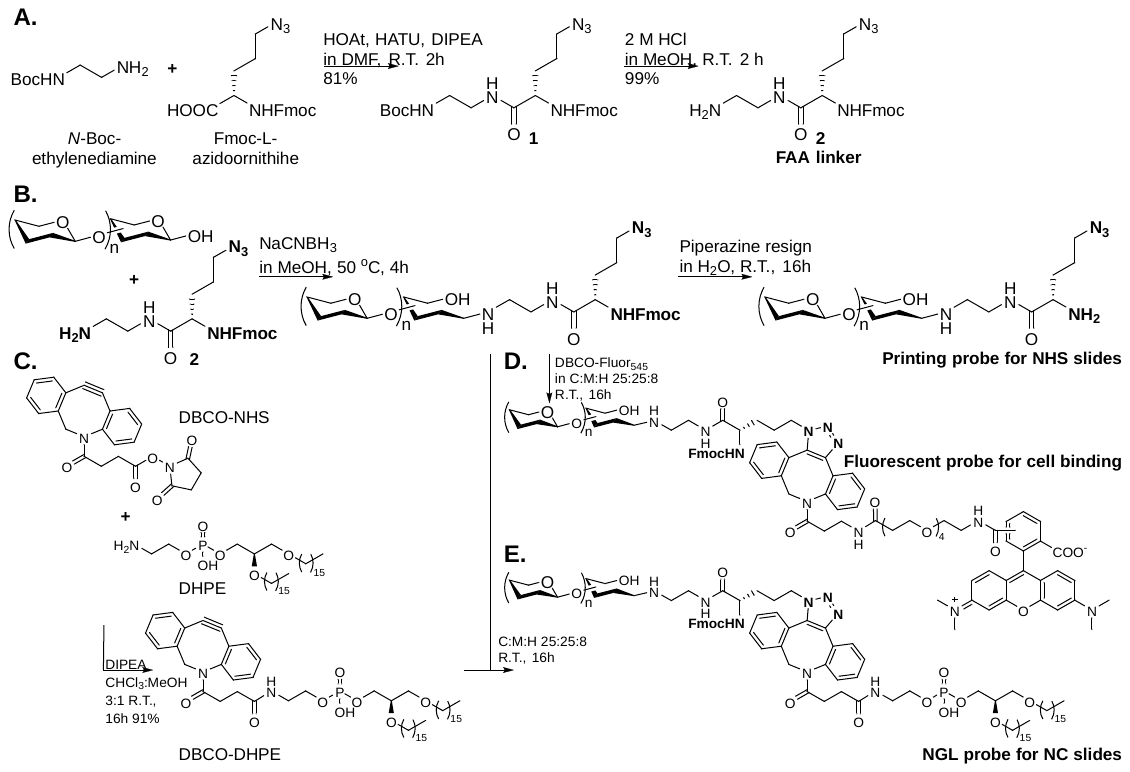

Scheme S4. Overall synthesis scheme of glycan probes for covalent and noncovalent arrays.

**A)** Synthesis of FAA linker from commercial Fmoc amino acid. **B)** Glycan labeling by reductive method and Fmoc remove on solid support. **C)** Synthesis of DBCO-DHPE reagent. **D)** Florescent labelling of the glycan probe. **E)** Conversion of the labelled glycan into NGL form.

Preparation of 2AB- and AEAB- tagged glycans for comparisons of chromatographic profiles with those tagged with FAA

For comparing coupling efficiency, 3 nmol each of the following six sequence-defined glycans were mixed together and tagged with FAA: cellobiose (Glc2), maltotriose (Mal3), α2,3-sialyllactose (3SL), lacto-N-tetraose (LNT), maltopentaose (Mal5) and maltoheptaose (Mal7), in a 10 μL solution containing 0.27 μmol FAA linker (15 eq). The HPLC profile of the tagged glycan mixture is in Figure S5A.

Aliquots, 0.5%, of the oligomannose *N*-glycan mixture from 40 mg RNAse B and of the complex type *N*-glycan mixture from fetuin (~ 10 ug scale) were tagged with 2AB^3^ and AEAB^4^ using reported procedures. The HPLC of the tagged glycans are in Figure S5B-D and Figure S5E-G, respectively.

Preparation of sequence defined glycans tagged with FAA

For generation of glycan microarrays, the 39 sequence-defined glycans (Table S1) were tagged with FAA. To quantify the amounts of tagged glycans based on their florescence intensities, calibration curves were generated using 2AB- and FAA-tagged maltotriose as standards (Figure S7). The HPLC profiles, MS spectra are in the in Figure S6. Selected MS spectra of probes after Fmoc removal are in Figure S8. The NGLs were prepared for noncovalent array and the MS spectra are shown in Figure S6. The products were quantified^1^. MS spectra of DBCO-PEG_4_-Flour545 modified glycan probes are in Figure S9.

Enzymatic modifications on *N*-glycan probe 36

The synthesis of biantennary *N*-glycan probes 37-41 (Figure S10) was performed by enzymatic modifications using published procedures^5^. In brief, for 2-3 sialyation, to a solution of probe 36 (100 nmol, 1.0 eq) and cytidine-5'-monophosphate-5-N-acetylneuraminic acid (CMP-Sia) (2.5 µmol, 25 eq) in 10 uL 0.1M Tris·HCl buffer containing 10 mM MgCl_2_, 2→3 sialyltransferase from *Pasteurella multocida* (Merck, Dorset, UK) (15 mU) and alkaline phosphatase from calf intestine (CIAP) (Merck, Dorset, UK) (10 U) were added sequentially. The reaction was kept at 37 ^o^C for 2 h and the product was purified on NP HPLC. For 1-2 fucosylation, to a microtube containing dry probe 36 (25 nmol, 1.0 eq) and 6-deoxy-β-L-galactopyranosylguanosine 5′-diphosphate (GDP-Fuc) (0.4 µmol, 16 eq), 25 uL 0.1M Tris·HCl buffer containing 10 mM MgCl_2_, 1→2 fucosyltransferase from *Helicobacter hepaticus* (Chemily, Georgia, US) (40 mU) and CIAP (10 U) were added. The reaction was kept at 37 ^o^C for 16 h and the product was purified on NP HPLC. For 1-3 fucosylation, to a solution of probe 36, 37 and 40 (10 nmol, 1.0 eq respectively) and GDP-Fuc (0.1 µmol, 10 eq) in 10 uL 0.1M Tris·HCl buffer containing 10 mM MgCl_2_ and 10 mM MnCl_2_, 1→3/4 fucosyltransferase from *Helicobacter mustelae* (Chemily, Georgia, US) (10 mU) and CIAP (10 U) were added. The reaction was kept at 37 ^o^C for 16 h and the product was purified on NP HPLC. The characterizations of the products are in Figure S6.

Generation of glycan microarrays

The printing of the microarrays was performed with a noncontact arrayer, Nano-Plotter from Gesim, Germany.

For printing, NHS coated TRIDIA (Codelink) and TRIDIA-High Density (Codelink-HD) glass microarray slides from Surmodics, Minnesota, USA, were used. Triplicate droplets (approximate 0.33 nL x3) were dispensed for each spot. Four replicate spots were made per glycan. Each subarray contained 44 glycans spotted as quadruplicates in a row; in total 11 rows were printed per subarray. Each arrayed slide contained 16 identical subarrays (pads).

For noncovalent array, NGLs and GLs were printed in a liposomal formulation^1^, which has been adjusted recently. In brief, each glycan probe was printed in duplicate at two levels (2 and 5 fmol per spot). To prepare the probes at the two levels, 50 and 150 pmol of lipid-linked probes (GLs and NGLs) were each dried with 1 nmol phosphatidylcholine (DHPC), 1 nmol cholesterol for 16 h. To generate the liposomes 10 µL of 20 ng/mL Cy3 NHS ester (GE Healthcare, Buckinghamshire, UK) in H_2_O was added followed by sonication at 35 ^o^C for 15 min. NC glass microarray slides (16-pad Slides) from Sartorius Stedim, Goettingen, Germany were used. One drop (approximately 0.33 nL) of the liposomes was dispensed for each spot. Each array slide contained 16 identical subarrays (pads). Each subarray contained up to 55 NGL or glycosylceramides which were printed at two levels in duplicate (four spots per saccharide in a row). Fourteen rows were printed per subarray.

Glycan binding proteins (GBPs)

The glycan binding proteins used are indicated in Table S2-6.

Microarray analysis of glycan binding

Microarray binding analyses were performed at R.T. essentially as described^1, 6, 7^. In brief, the covalent slides were covered with 1 mM ethanolamine at pH 9 for 1 h and the NC slides were treated with blocking buffer for 1 h prior to incubations. All overlay steps of the VP1 virus proteins were performed in blocking buffer A containing 0.33%BSA, 0.33% Casein in HEPES-Buffered Saline (HBS) (5 mM HEPES buffer pH 7.4, 150 mM NaCl) whereas the incubations of lectins and antibodies were performed in blocking buffer B containing 1%BSA, 10 mM CaCl_2_ in HBS. The concentration of the antibodies and lectins are indicated in Table S2-6. The stepwise incubations were performed 1 h per overlay step, except for the incubation of VP1 virus proteins were performed with one step pre-complexed condition with VP1 protein:anti His:anti-M IgG 150:75:37.5 μg/mL for 1.5 h. For on-slide visualization of printed covalent probes, the slide was incubated with a solution of DBCO-PEG_3_-Biotin in 1:1 DMF:H_2_O (1 nM to 500 µM) at R.T. overnight. The final detection step was carried out by 30 min overlay with 1 μg/mL Alexa Fluor-647-labeled streptavidin (Molecular Probes, Eugene).

The incubated air-dried slides were scanned using a GenePix 4300A instrument from Molecular Devices, Sunnyvale, USA. The gpr file containing the quantified binding data was entered into Excel to generate the histogram and data matrix. The scanning of the incubated slides was carried out at a resolution of 5 µm/pixel. For NC slides a PMT Voltages at 350V and the scan power: ranging from 30 to 100% was used to avoid saturation of binding signals, whereas for covalent slides, PMT Voltages at 400V and the scan power at 100% was used.

Mammalian cell culture and generation of HEK Siglec 9 cells.

HEK Null1 cells were obtained from InvivoGen and were cultured and maintained under standard conditions in DMEM medium (Sigma Aldrich) containing 10% decomplemented fetal bovine serum (Sigma Aldrich) and 1% penicillin– streptomycin (Sigma Aldrich). The cells were cultured at 37 °C and 5% CO_2_ in a humidified incubator, were free of mycoplasma and routinely tested using the MycoStrip Detection Kit (InvivoGen). HEK Null1 cells were seeded at a density of 2 × 10^6^ cells per well in a 6 well plate (Corning) and incubated 4h, after which cells were transfected with a mixture of pCMV SPORT6 Siglec 9 (2.5 μg; Horizon Discovery) and Lipofectamine 2000 (10 μl; ThermoFisher) in OptiMEM (250ul; Gibco). 24 h after transfection, the medium was removed and the cells were gently detached in PBS by flushing.

Glycan binding to HEK Siglec 9 cells.

The F545-FAA-glycan probes were added to 5 × 10^5^ HEK Null1 cells or HEK Siglec 9 cells in 500ul PBS at a final concentration of 50pM and the cells were stirred on a wheel at 4°C for 1h. Glycan labelling of the cells was determined by flow cytometry analysis on a FACSCalibur (BD Biosciences) and analyzed using the CellQuest software (BD Biosciences). Viability of the cells before and after the glycan binding were approximately 85% for all the conditions tested as determined by trypan blue cell counting. Additionally, 10^5^ cells were transferred in a black 96 well plate and total fluorescence was measured on a BMG PHERAStar Plus plate reader (BMG Labtech).

SUPPLEMENTARY RESULTS AND DISCUSSION

Details of binding signals given by two anti-ganglioside antibodies

**Anti-GD1a** gave binding to GD1a glycan probes (#31) in covalent and noncovalent array, and GD1a ceramide (#46) in noncovalent array (Figure S12A).

**Anti-Gb3** gave binding to Gb3 glycan probes (#15) in all three arrays specifically (Figure S12B).

Details of binding signals given by Sia, Man, Gal, GalNAc terminating glycans using lectins

**SNA** bound to the two terminal α2-6 sialylated probes (#25, #42) as predicted. Weak binding to terminal α2-3 sialylated biantennary *N*-glycan (#40) was also observed (Figure S13A).

**MAA-I** gave binding signals with terminal α2-3 Sia on type 2 LacNAc probes (#24, #40 **Fig S13B**) as predicted. On the covalent arrays weak bindings to the type2 LacNAc probes in the absence of sialylation (#6, #7, #36 **Fig S13B**) was observed, which is consistent with its specificity to type 2 backbones.

**ConA** Binding signals to high-mannose and complex type *N*-glycans were in accord with previous knowledge (Figure S14A).

**PNA** gave binding with all of the Gal-GalNAc terminating globoside and ganglioside glycan probes (#17, #29, #32, #35) as predicted (Figure S14B).

**RCA 120** binding to terminal type2 LacNAc (#6, #7, #36) 2-6 sialyl LacNAc probes (#41) are in accordance with previous knowledge. The relatively weak to biantennary Le^x^ terminating *N*-glycan is noted (Figure S14C).

**WGA** binding to NH-acetyl terminating glycan probes, GalNAc terminating blood group A related (#1, #2, #20), Forssmann antigen (#14) and Sia terminating (#23, #24, #27, #28) are in accord with previous knowledge (Figure S14D). The relatively strong binding on the covalent arrays to LacNAc or poly LacNAc based sequences (#4, #7, #9, #19) likely to reflect the relatively high concentration of the lectin used in this study^8^.

Table S1. Designations of sequence defined glycans. Their sequences are in Excel Table S1 and S2. The definition of the color pattern was given in Fig 1 in the main text.

| **No*** | **Short name** | **Full name** | **Vendor** | **Code** |
| --- | --- | --- | --- | --- |
| **1** | A-penta-T1 | Blood group A antigen pentaose type 1 | Elicityl | GLY036-1 |
| **2** | A-hexa-T2 | Blood group A antigen hexaose type 2 | Elicityl | GLY037-2 |
| **3** | B-hexa-T1 | Blood group B antigen hexaose type 1 | Elicityl | GLY040-1 |
| **4** | B-hexa-T2 | Blood group B antigen hexaose type 2 | Elicityl | GLY040-2 |
| **5** | LNT | Lacto-N-tetraose | Dextra | L403 |
| **6** | LNnT | Lacto-N-neotetraose (LNnT / neo-LNT) | Dextra | GLY021 |
| **7** | LN3 | Tri-N-Acetyl-D-Lactosamine | Elicityl | GLY007-2 |
| **8** | LNFPI | Lacto-N-fucopentaose I | Dextra | L502 |
| **9** | LNnFPI | Lacto-N-neofucopentaose I (LNnFP I) | Elicityl | GLY033-2 |
| **10** | LNFPII | Lacto-N-fucopentaose II | Dextra | L503 |
| **11** | Le^b^ | Lewis b (Le^b^) pentaose | Elicityl | GLY056 |
| **12** | Le^x^ | Lewis x (Le^x^) tetraose | Elicityl | GLY050 |
| **13** | Le^y^ | Lewis y (Le^y^) pentaose | Elicityl | GLY052 |
| **14** | FA-penta | Forssman antigen pentaose | Elicityl | GLY132 |
| **15** | Gb3 | Globotriaose (Gb3) / Pk antigen | Elicityl | GLY120 |
| **16** | Gb4 | Globotetraose (Gb4) / P antigen | Elicityl | GLY121 |
| **17** | Gb5 | Globopentaose/Stage Specific Embryonic Antigen 3a (SSEA-3a) | Elicityl | GLY122 |
| **18** | Globo-H | Globo-H hexaose /Stage Specific Embryonic Antigen 3b (SSEA-3b) | Elicityl | GLY123 |
| **19** | Globo-H-T2 | Globo-H analogue type 2 | Elicityl | GLY194-2 |
| **20** | Globo-A | Globo-A heptaose | Elicityl | GLY124 |
| **21** | Globo-B | Globo-B heptaose | Elicityl | GLY125 |
| **22** | 3'-SL | 3'-Sialyllactose | Dextra | SL302 |
| **23** | LST a | LS-Tetrasacchride a | Dextra | SLN503 |
| **24** | LSTd | LS-Tetrasaccharide d/ Sialyl-lacto-N-neotetraose d (Sodium salt) | Elicityl | GLY083 |
| **25** | LST c | LS-Tetrasacchride c | Dextra | SLN506 |
| **26** | LSTb | LS-Tetrasaccharide b | Dextra | SLN516 |
| **27** | SSEA4-hexa | SSEA-4 hexaose/ Stage-Specific Embryonic Antigen-4 hexaose | Elicityl | GLY131 |
| **28** | SLe^x^ Penta | Sialyl Lewis x (sLe^x^) pentaose - Sodium salt | Elicityl | GLY053 |
| **29** | GM1a | GM1a Ganglioside oligosaccharide | Elicityl | GLY096 |
| **30** | GM1b | GM1b Ganglioside oligosaccharide | Elicityl | GLY097 |
| **31** | GD1a | GD1a Ganglioside oligosaccharide | Elicityl | GLY098 |
| **32** | GalGalNAc-aGM1 | GalGalNAc asialo GM1 Ganglioside oligosaccharide | Elicityl | GLY174 |
| **33** | GT1a | GT1a Ganglioside oligosaccharide | Elicityl | GLY100 |
| **34** | GD2 | GD2 Ganglioside oligosaccharide | Elicityl | GLY094 |
| **35** | GD1b | GD1b Ganglioside oligosaccharide | Elicityl | GLY099 |
| **36** | H5N4 | Asialo, galactosylated, biantennary (NA2) | Dextra | C0920 |
| **37** | F2H5N4(H2) | FAA tagged, enzymatically modified from probe 36 |  |  |
| **38** | F2H5N4(Le^x^) | FAA tagged, enzymatically modified from probe 36 |  |  |
| **39** | F2H5N4(Le^y^) | FAA tagged, enzymatically modified from probe 36 |  |  |
| **40** | A2H5N4(2-3Sia) | FAA tagged, enzymatically modified from probe 36 |  |  |
| **41** | A2F2H5N4(SLe^x^) | FAA tagged, enzymatically modified from probe 36 |  |  |
| **42** | A2H5N4(2-6Sia) | Disialo (2,6), biantennary (A2) | Dextra | SC1120 |
| **43** | M5N2 | Oligomannose-5 glycan | Dextra | MC0731 |
| **44** | M9N2 | Oligomannose-9 glycan | Dextra | MC1131 |
| ** | Glc2 | Cellobiose: Glcβ1→4Glc | Dextra | BC121 |
| ** | Mal3 | Maltotriose: Glc(α1→4Glc)_2_ | Dextra | G302 |
| ** | Mal5 | Maltopentaose: Glc(α1→4Glc)_4_ | Dextra | G502 |
| ** | Mal7 | Maltoheptaose: Glc(α1→4Glc)_6_ | Dextra | G702 |

* Glycan numbers refer to positions in sequence defined array.
** Additional glycans were used as reference FAA tagged compounds.

Table S2. Anti-carbohydrate antibodies.

| Antibody | Other  designation | Clone | Vendor or collaborator | Catalogue # | Isotype | Dilution used |
| --- | --- | --- | --- | --- | --- | --- |
| Anti-Le^a^ | BG5 | T174 | Biolegend | 922202 | Mouse IgG | 1:50 |
| Anti-Le^b^ | BG6 | T218 | Biolegend | 922302 | Mouse IgM | 1:50 |
| Anti-Le^x^ | L5* |  | Andrea Streit |  | Rat IgM | 1:50 |
| Anti-Le^x^ | BG7 | P12 | Biolegend | 912901 | Mouse IgM | 1:50 |
| Anti-Le^x^ | SSEA1 |  | DHSB | MC-480-c | Mouse IgM | 1:50 |
| Anti-Le^y^ | BG8 | F3 | Biolegend | 912501 | Mouse IgM | 1:50 |
| Anti-Le^y^ | AH6** |  | Henrik Clausen |  | Mouse IgM | 1:5 |
| Anti-A | BG2 | T36 | Biolegend | 921902 | Mouse IgG | 1:10 |
| Anti-B | HEB29 | - | Abcam | Ab2524 | Mouse IgM | 1:10 |
| Anti-H type 1 | H(O) type 1 | 17-206 | Invitrogen | 14-9810-82 | Mouse IgG | 1:50 |
| Anti-H type 2 | BE2** |  | Henrik Clausen |  | Mouse IgM | 1:2 |
| Anti-SLe^x^ | CD15S | CSLEX1 | BD Pharmingen | 551344 | Mouse IgM | 1:50 |
| Anti-GD1a |  |  | Millipore | MAB5606 | Mouse IgG | 1:250 |
| Anti-Gb3 |  |  | TCI | A2506 | Mouse IgG | 1:140 |

* Rat anti-L5 purified from culture supernatant was a gift from Dr Andrea Streit (King’s College London)^9^.
** Anti-H type 2 (BE2) and Anti-Le^y^ (AH6) was kindly provided by Henrik Clausen Lab, Copenhagen Center for Glycomics^10^.

Table S3. Detection antibodies.

The dilution was indicated for stepwise incubation.

|  | Vendor | Cat | Stock conc. | Used dilution |
| --- | --- | --- | --- | --- |
| Anti-Mouse IgG (biotinylated) | Sigma | B7264 | 0.4 mg/mL | 1:200 |
| Anti-Mouse IgM (biotinylated) | Vector | BA-2020 | 0.5 mg/mL | 1:200 |
| Anti-Rat IgM (biotinylated) | Rockland | 612-4607 | 1.0 mg/mL | 1:200 |
| Anti-Rabbit IgG (biotinylated) | Sigma | B7389 | 0.6 mg/mL | 1:200 |
| Anti-Human IgG (biotinylated) | Vector | BA-3000 | 1.5 mg/mL | 1:200 |
| Anti-Human IgM (biotinylated) | Vector | BA-3020 | 0.5 mg/mL | 1:200 |
| Anti-His (Mouse IgG isoform) | Sigma | SAB4200620 | 1.0 mg/mL | 1:100 |
| Anti-Cholera toxin (Rabbit IgG) | Sigma | C3062 | 1.0 mg/mL | 1:200 |

Table S4. Biotinylated lectins.

|  | Vendor | Cat | Stock conc. | Used conc. |
| --- | --- | --- | --- | --- |
| *Aleuria Aurantia* Lectin (AAL) | Vector Laboratories | B-1395 | 2 mg/mL | 20 μg/mL |
| *Ulex Europaeus* Agglutinin I (UEA-I) | Vector Laboratories | B-1065 | 2 mg/mL | 50 μg/mL |
| *Lotus Tetragonolobus* Lectin (LTL) | Vector Laboratories | B-1325 | 2 mg/mL | 100 μg/mL |
| Concanavalin A (ConA) | Vector Laboratories | B-1005 | 5 mg/mL | 5 μg/mL |
| *Sambucus Nigra* Lectin (SNA) | Vector Laboratories | B-1305 | 2 mg/mL | 50 μg/mL |
| *Maackia Amurensis* Lectin I (MAA-I) | Vector Laboratories | B-1315 | 2 mg/mL | 100 μg/mL |
| *Maackia Amurensis* Lectin II (MAA-II) | Vector Laboratories | B-1265 | 1 mg/mL | 100 μg/mL |
| Peanut Agglutinin (PNA) | Vector Laboratories | B-1075 | 5 mg/mL | 50 μg/mL |
| *Ricinus Communis* Agglutinin I (RCA 120) | Vector Laboratories | B-1085 | 5 mg/mL | 5 μg/mL |
| Wheat Germ Agglutinin (WGA) | Vector Laboratories | B-1025 | 5 mg/mL | 20 μg/mL |

Table S5. Glycan binding receptors of the immune system

|  | **Source** | **Tag** | **Incubation conditions** |
| --- | --- | --- | --- |
| Human DC-SIGN | Sino Biological, 10200-H01H | hFc tagged | 5 μg/mL; pre-complexed with biotinylated anti-human IgG (1:2 by weight) |
| Human DC-SIGNR | Sino Biological, 10559-H01H | hFc tagged | 20 μg/mL; pre-complexed with biotinylated anti-human IgG (1:2 by weight) |
| Rhesus Langerin | Sino Biological, 90159-C01H | hFc tagged | 20 μg/mL; pre-complexed with biotinylated anti-human IgG (1:2 by weight) |
| Human Siglec-7 | R&Dsystems, 1138-SL-050 | hFc tagged | 5 μg/mL; pre-complexed with biotinylated anti-human IgG (1:1 by weight) |
| Human Siglec-9 | R&Dsystems, 1139-SL-050 | hFc tagged | 3 μg/mL; pre-complexed with biotinylated anti-human IgG (1:1 by weight) |
| Human E-Selectin (CD62E) | SinoBiological, 10335-h08h | His tagged | 20 μg/mL; pre-complexed protein:anti-His: biotinylated anti-mouse IgG (8:6:3 by weight) |
| Murine E-Selectin IgM* | John B. Lowe, University of Michigan Medical School^11^ | Human IgM | 1:4 dilution |

* E-Selectin was kindly provided by John B. Lowe, University of Michigan Medical School.

Table S6. Microbial glycan binding proteins used in array analysis.

|  | **Source** | **Tag** | **Incubation conditions** |
| --- | --- | --- | --- |
| Wild type (WT) BKPyV VP1 | Thilo Stehle laboratory  (University of Tübingen) | His tagged | 150 μg/mL; pre-complexed protein:anti-His: biotinylated anti-mouse IgG (4:2:1 by weight) |
| SV40 VP1 |  |  |  |
| JCPyV VP1 (Mad-1) |  |  |  |
| Classical *Vibrio cholerae* Toxin Subunit B (cCTB) | Sigma-Aldrich | N/A | 5 μg/mL and 0.5 μg/mL |
| El Tor Cholera Toxin Subunit B (El Tor CTB)^12, 13^ | Bruce Turnbull laboratory  (University of Leeds) |  |  |
| *Escherichia Coli* heat-labile Toxin (LTBh)^14^ |  |  |  |

Table S7. Highlights of glycan binding readouts from the covalent and noncovalent arrays

|  | **Equivalent in noncovalent and covalent** | **Subtle but significant differences** | **Striking difference** | |
| --- | --- | --- | --- | --- |
|  |  |  | **Preference for covalent arrays** | **Preference for noncovalent array** |
| Plant lectins (9) | AAL, ConA | UEA-I, PNA, RCA120, SNA, MAA-I, WGA | LTL |  |
|  | - Lack of binding of AAL to blood group A and B glycans as recently reported^15^. - In addition to Le^x^ and Le^y^, LTL also bound to SLe^x^ and H type 2 (not reported in microarray studies). | | | |
| Anti-glycan antibodies (14) | Anti-A (BG2)  Anti-B (HEB29)  Anti-H1 (17-206)  Anti-H2 (BE2)  Anti-Le^a^ (BG5)  Anti-Le^b^ (BG6)  Anti-Le^x^ (anti-L5, BG-7, anti-SSEA-1)  Anti-Le^y^ (AH6)  Anti-SLe^x^  Anti-GD1a  Anti-Gb3 | Anti-Le^y^ (BG8) |  |  |
|  | - Anti-H type 1 (clone 17-206) bound weakly to blood group B-type 1 in addition to H-type 1 probes. - Lack of binding anti-H type 2 to globo H analog 2 (not reported). - Anti-Le^x^ (anti-L5) weakly bound to Le^a^ probe LNFPII as previously reported^16^. - Anti-Le^y^ BG8 (clone F3) bound to H type 2, and strong binding to Le^x^ probes observed only on the HD slide. - Lack of binding of anti-Le^x^ (BG-7 as with anti-SSEA-1) to short chain Le^x^ and Le^x^ on biantennary N-glycan. - Lack of binding of anti SLe^x^ to SLe^x^ presented on short outer arms of biantennary N-glycan (reminiscent of the “masking effect”)^17^ | | | |
| Immune lectins (7) | hE-selectin  mE-selectin | hDC-SIGN  hDC-SIGNR  Rhesus Langerin |  | hSiglec-7  hSiglec-9 |
|  | - Binding of DC-SIGNR to Le^x^-terminating complex type *N*-glycan (not reported). - Murine E-selectin (IgM construct) but not his-tagged human E-selectin gave weak binding to Le^a^, Le^b^ probes reminiscent of high density E-selectin expression on cells^18^. | | | |
| Viral adhesins (3) | SV40 VP1 |  |  | JCPyV VP1  BKPyV VP1 |
|  | - Binding of SV40 VP1 to disialyl glycolipid GalNAc-GD1a (not reported). - JCPyV VP1 binding to sialyl glycans beyond LSTc (new finding). | | | |
| Bacterial toxins (3) |  | cCTB  El Tor CTB  LTBh |  |  |
|  | - The three toxins all bound strongly to a range of gangliosides, including GM1a and GD1b. - Covalent and non-covalent arrays revealed differential binding to low affinity glycan ligands. - The toxins also have differing preference for blood-group probes:   cCTB: A type 1, Le^y^, SLe^x^;  El Tor CTB: H type 2, Le^y^, Le^b^;  LTBh: A and B, H type 2, Le^y^. | | | |

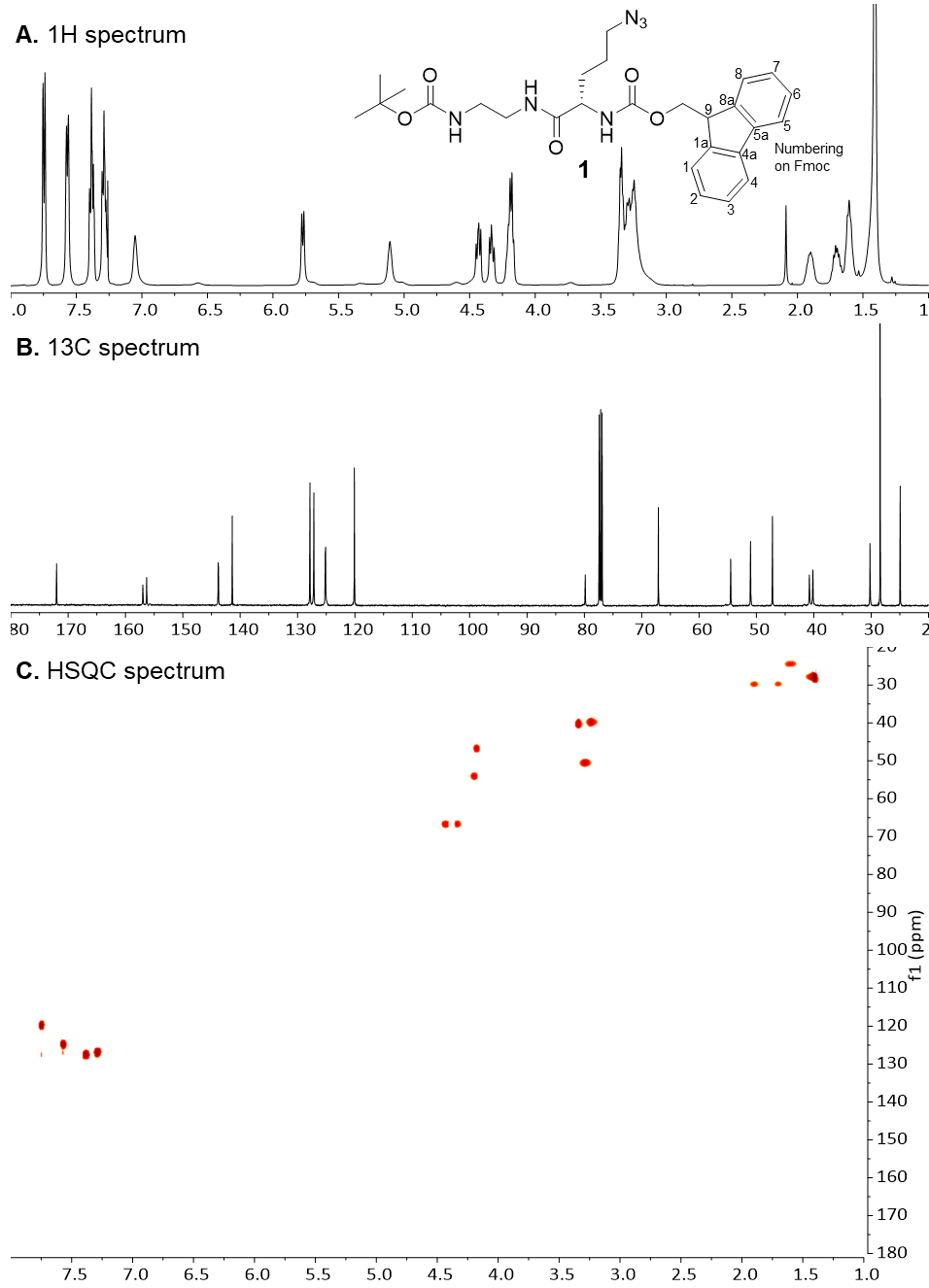

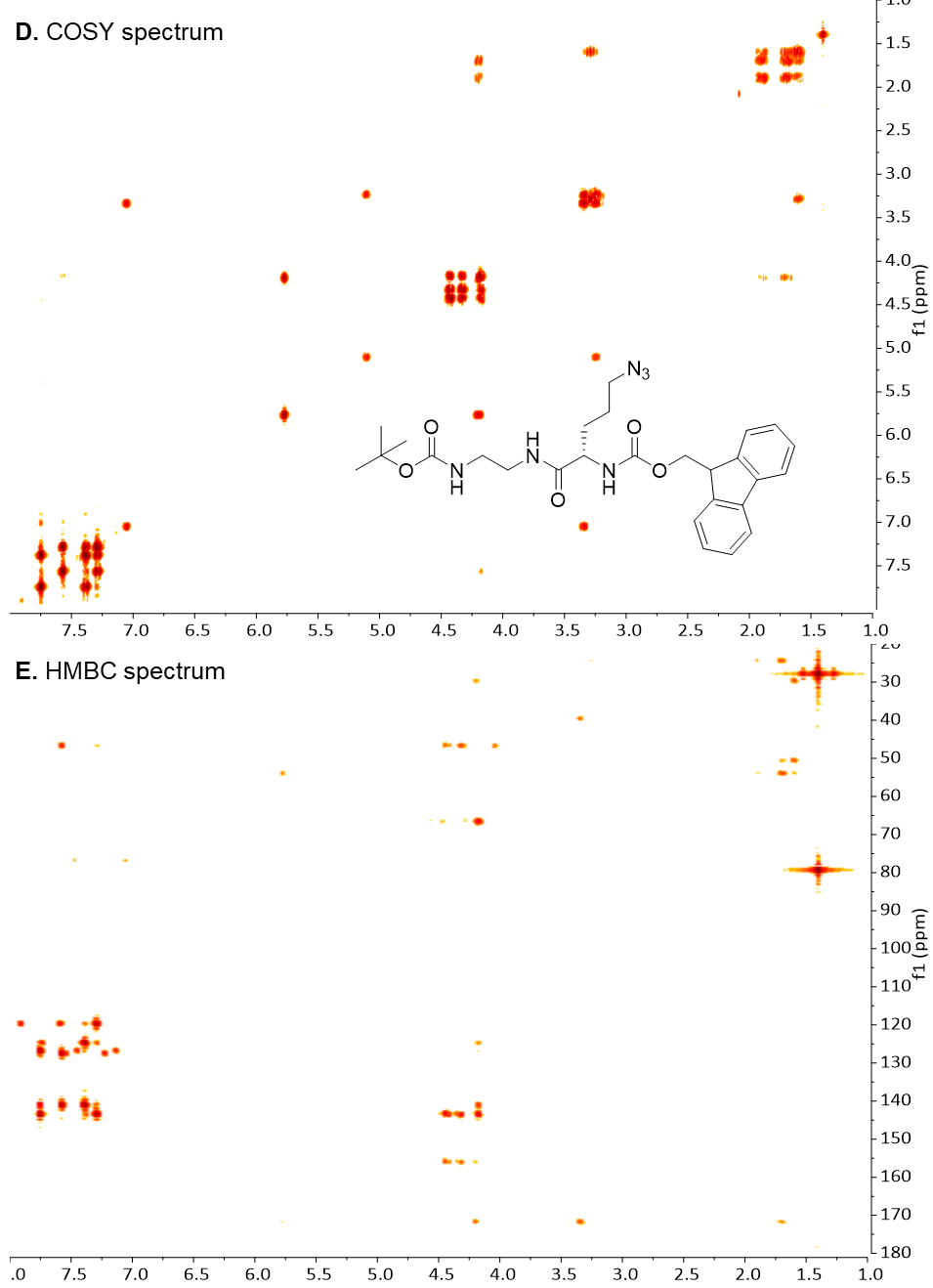

Figure S1. NMR spectra of compound 1.

**A)** 1H spectrum. **B)** 13C spectrum. **C)** HSQC spectrum. **D)** COSY spectrum. **E)** HMBC spectrum.

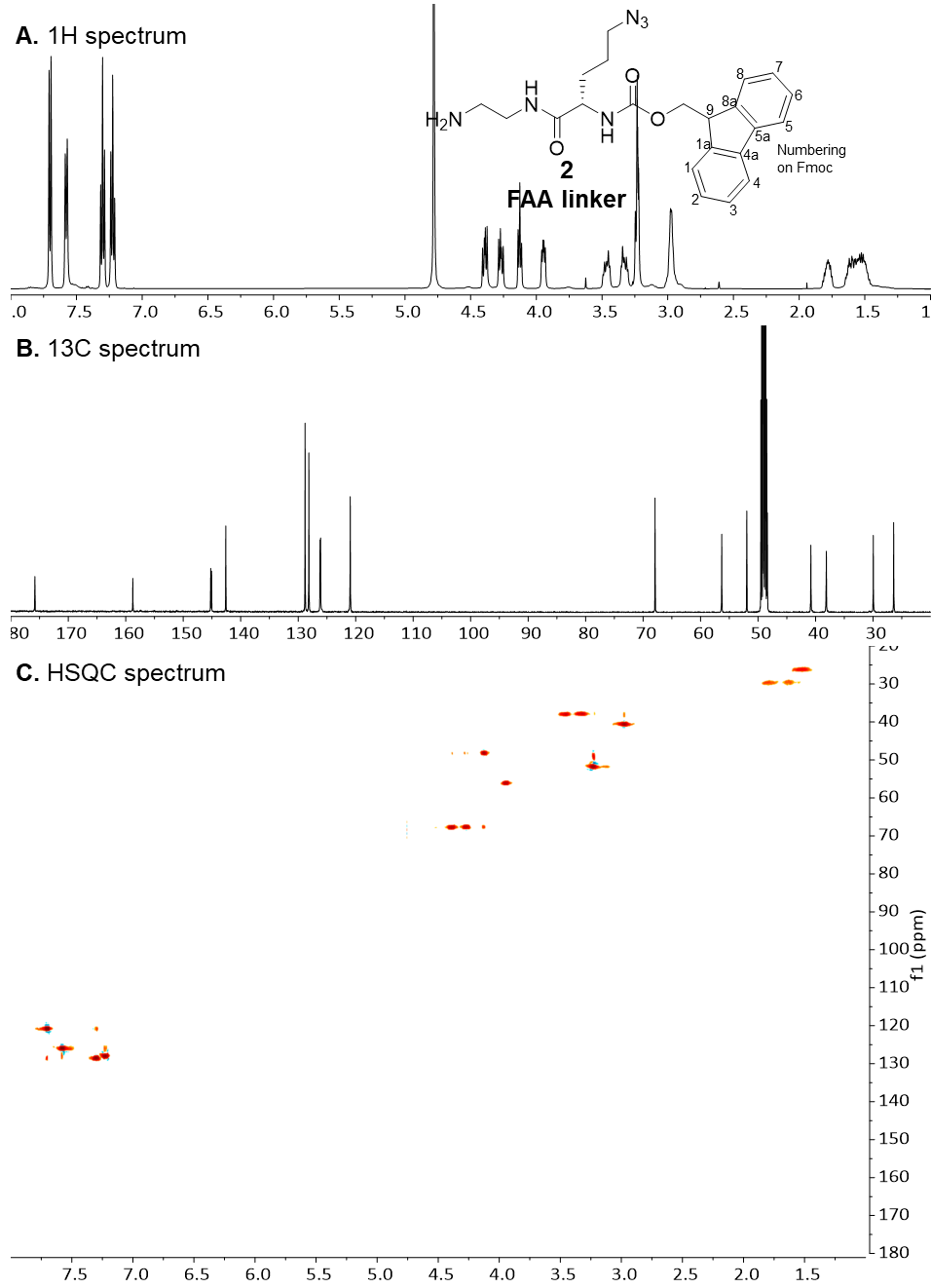

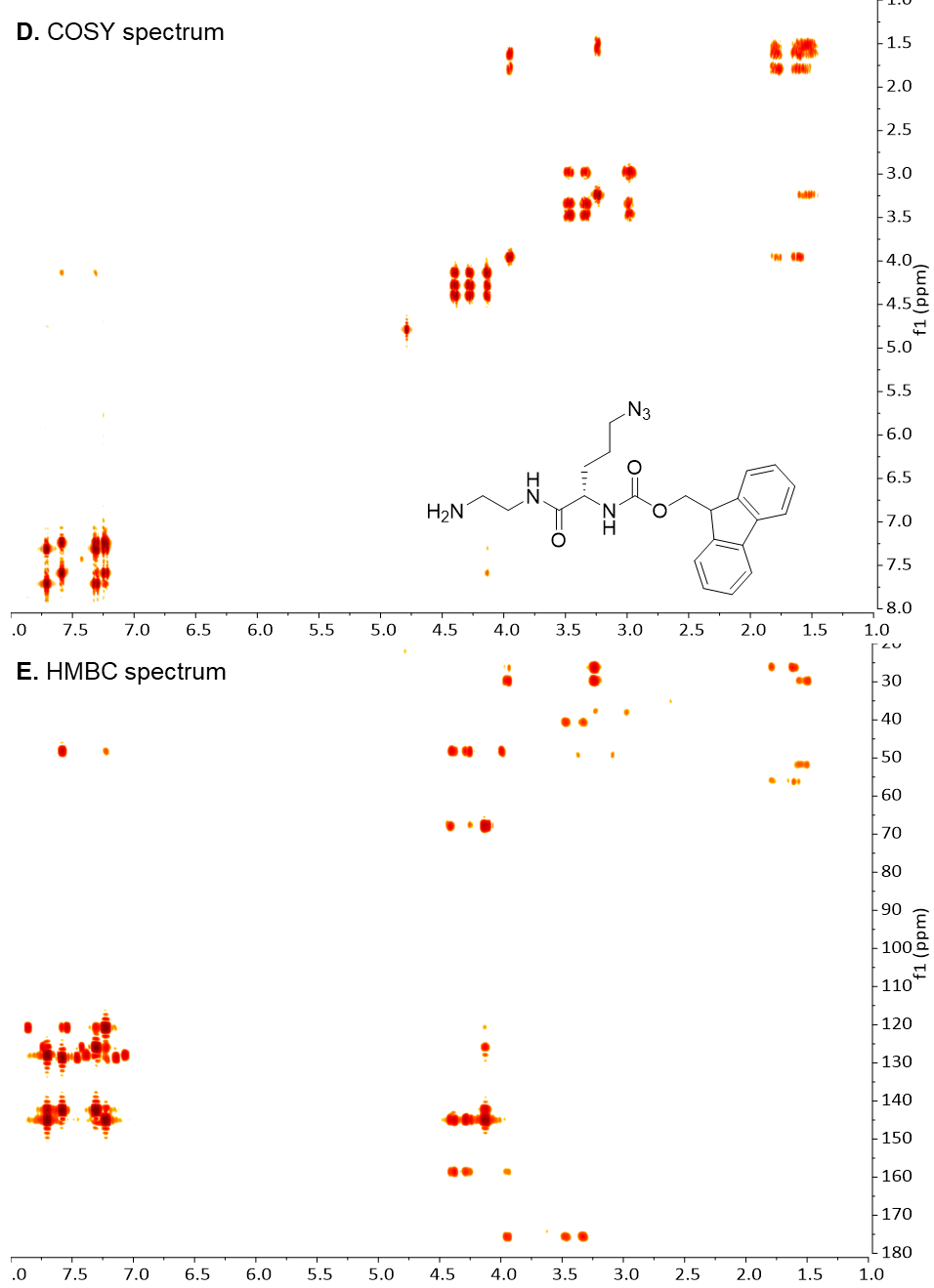

Figure S2. NMR spectra of compound 2.

**A)** 1H spectrum. **B)** 13C spectrum. **C)** HSQC spectrum. **D)** COSY spectrum. **E)** HMBC spectrum.

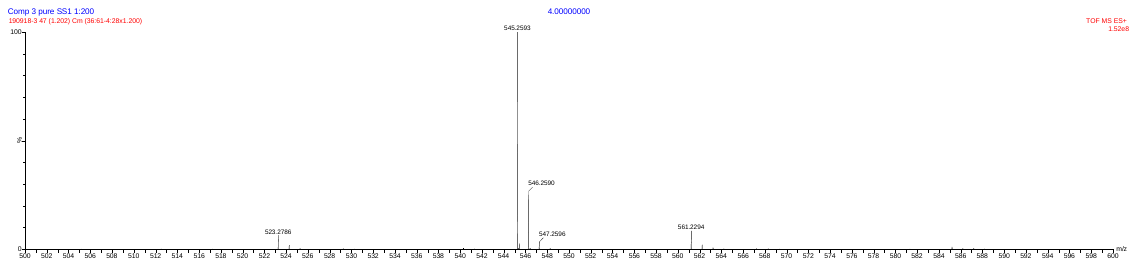

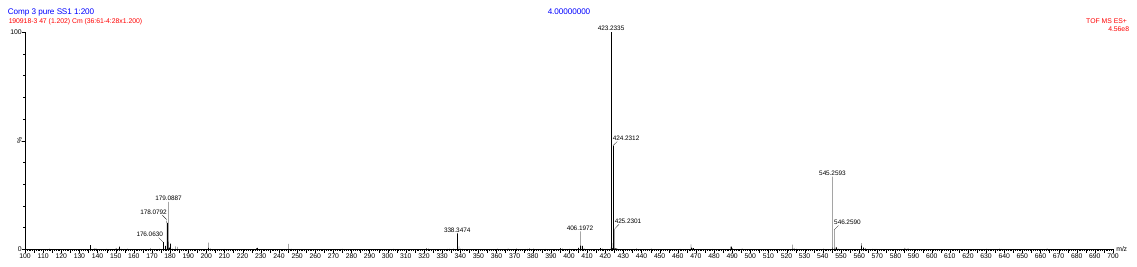

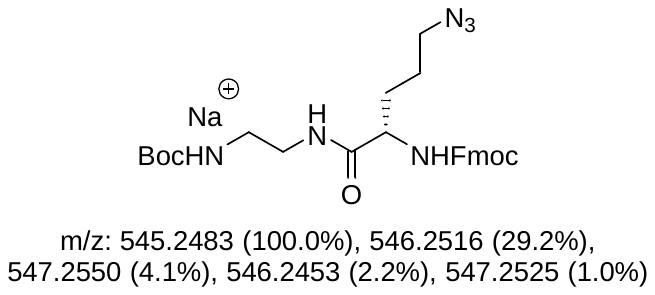

Figure S3. HI-ESI-MS spectra of compound 1

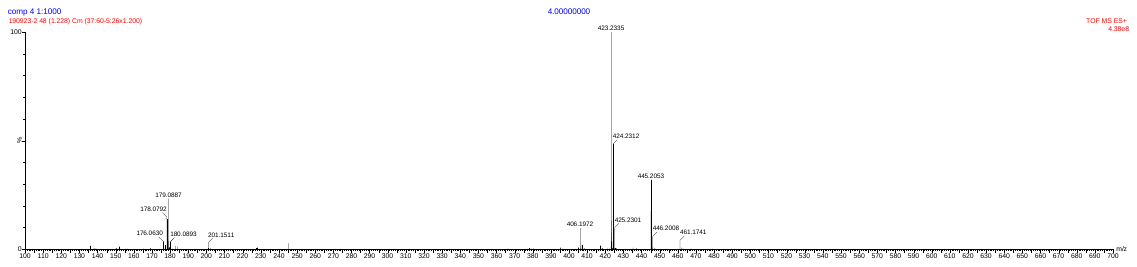

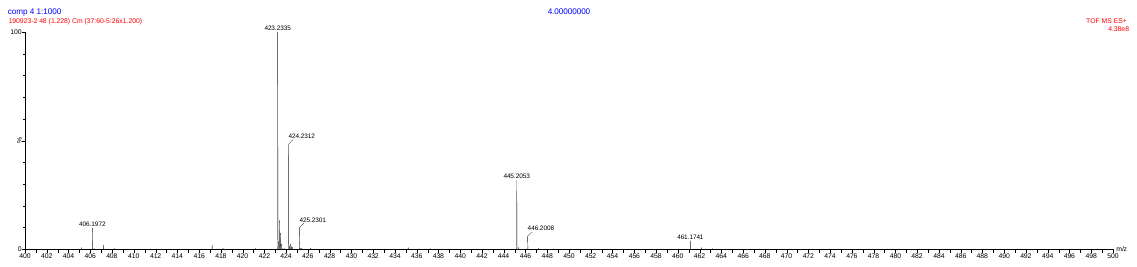

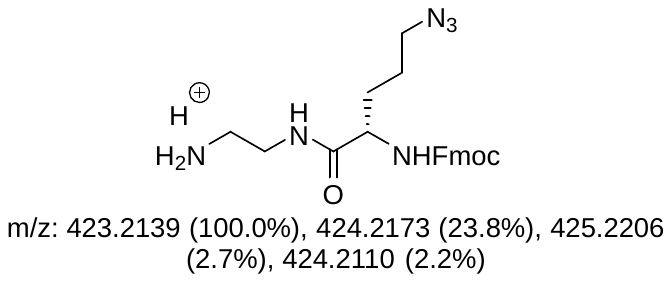

Figure S4. HI-ESI-MS spectra of compound 2

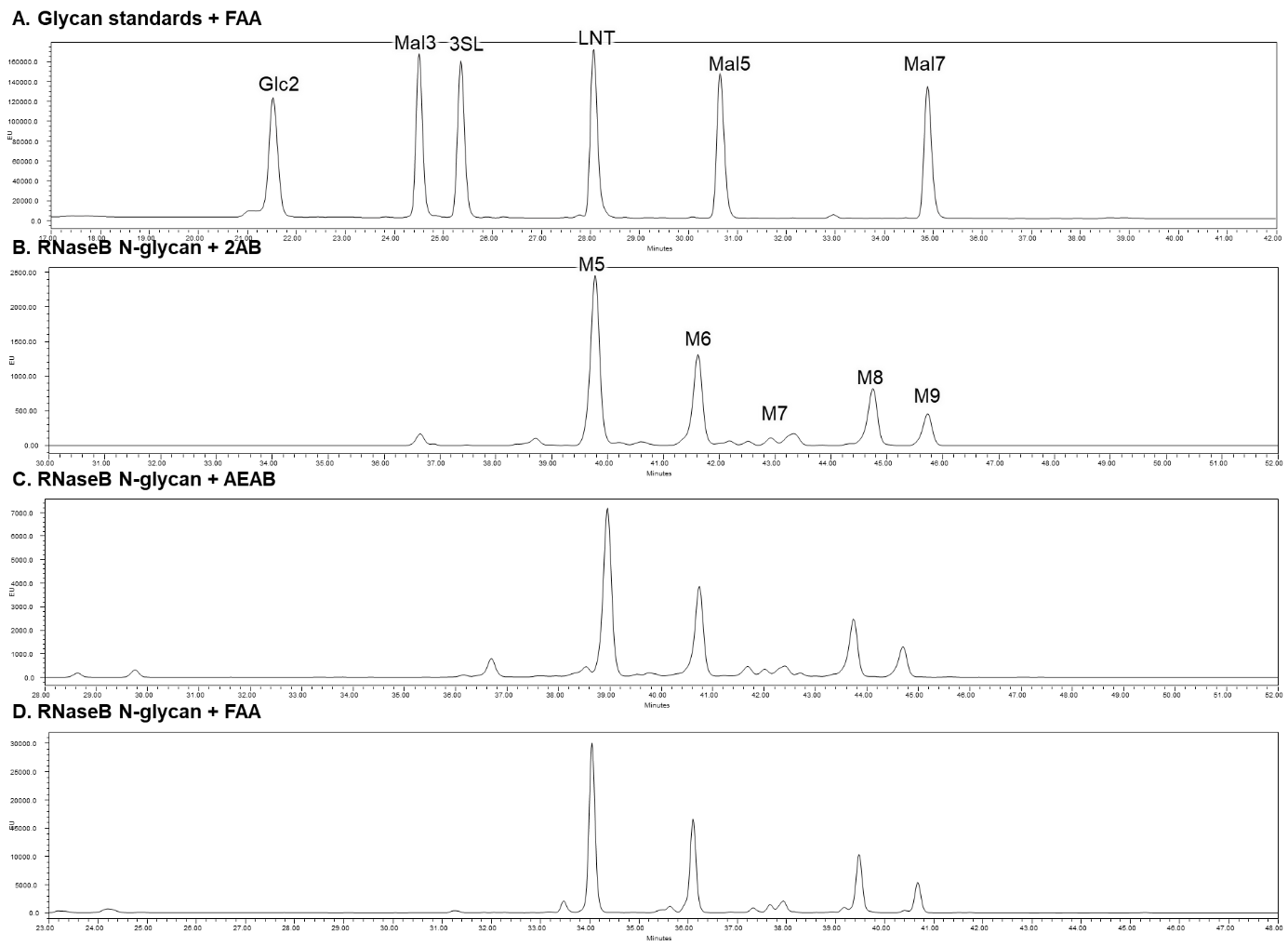

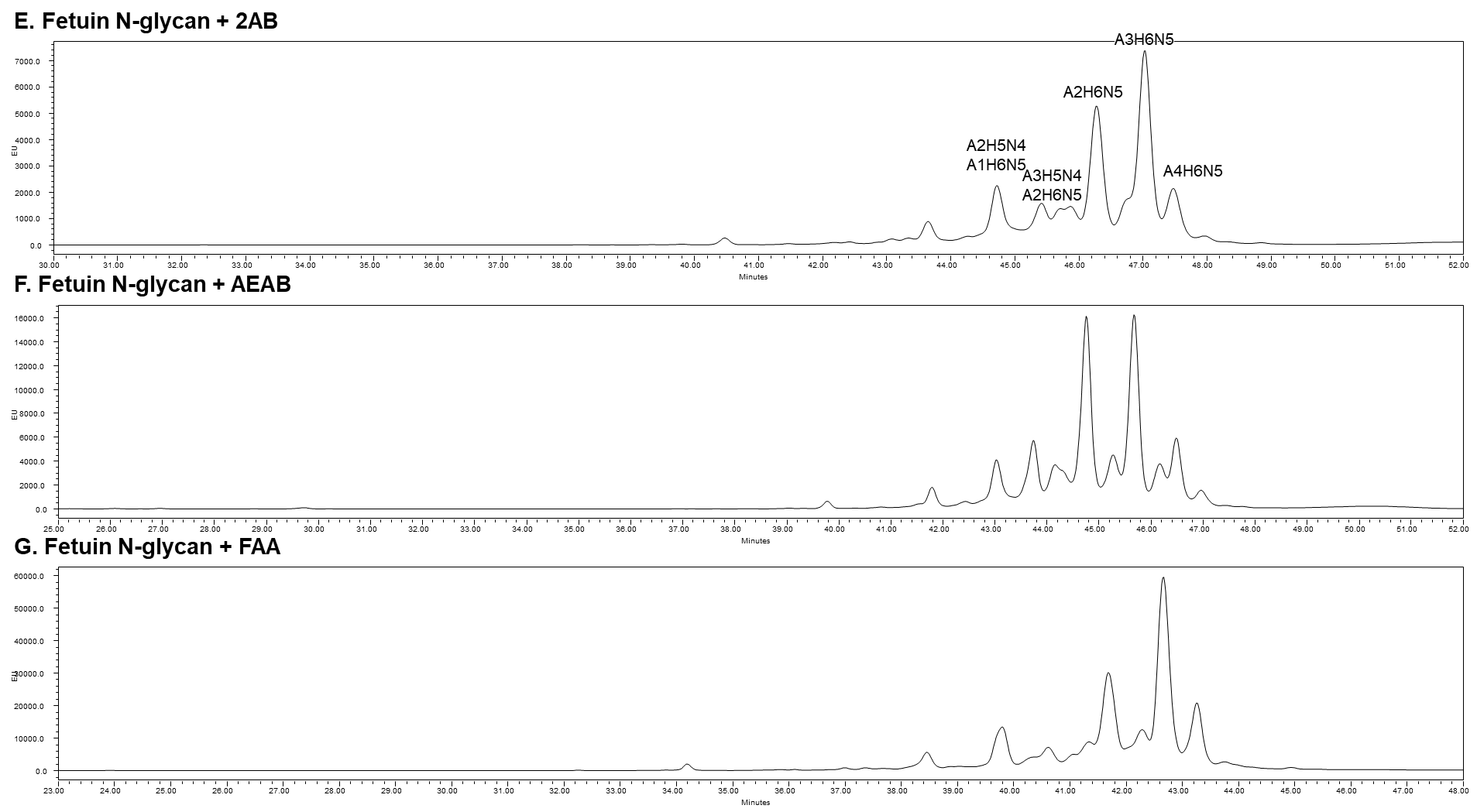

Figure S5. Comparative HPLC charts of glycans tagged withFAA, 2AB and AEAB .

**Panel A:** Mixture of sequence defined glycans, 3 nmol of each, tagged with FAA. Panels **B-D**: Aliquots, 0.5% of oligomannose mixture released with EndoH from 40 mg RNaseB tagged with 2AB (panel **B)**, with AEAB (panel **C)**, and with FAA (panel **D)**. Panels **E-G):** Aliquots, 0.5% of *N*-glycan mixture released with PNGF from 40 mg fetuin tagged with 2AB (panel **E)**, with AEAB (panel **F)** and with FAA (panel **G).**

Figure S6. HPLC profiles and MS spectra of 44 FAA-linked sequence defined glycans and corresponding NGLs

**Probe No 1**

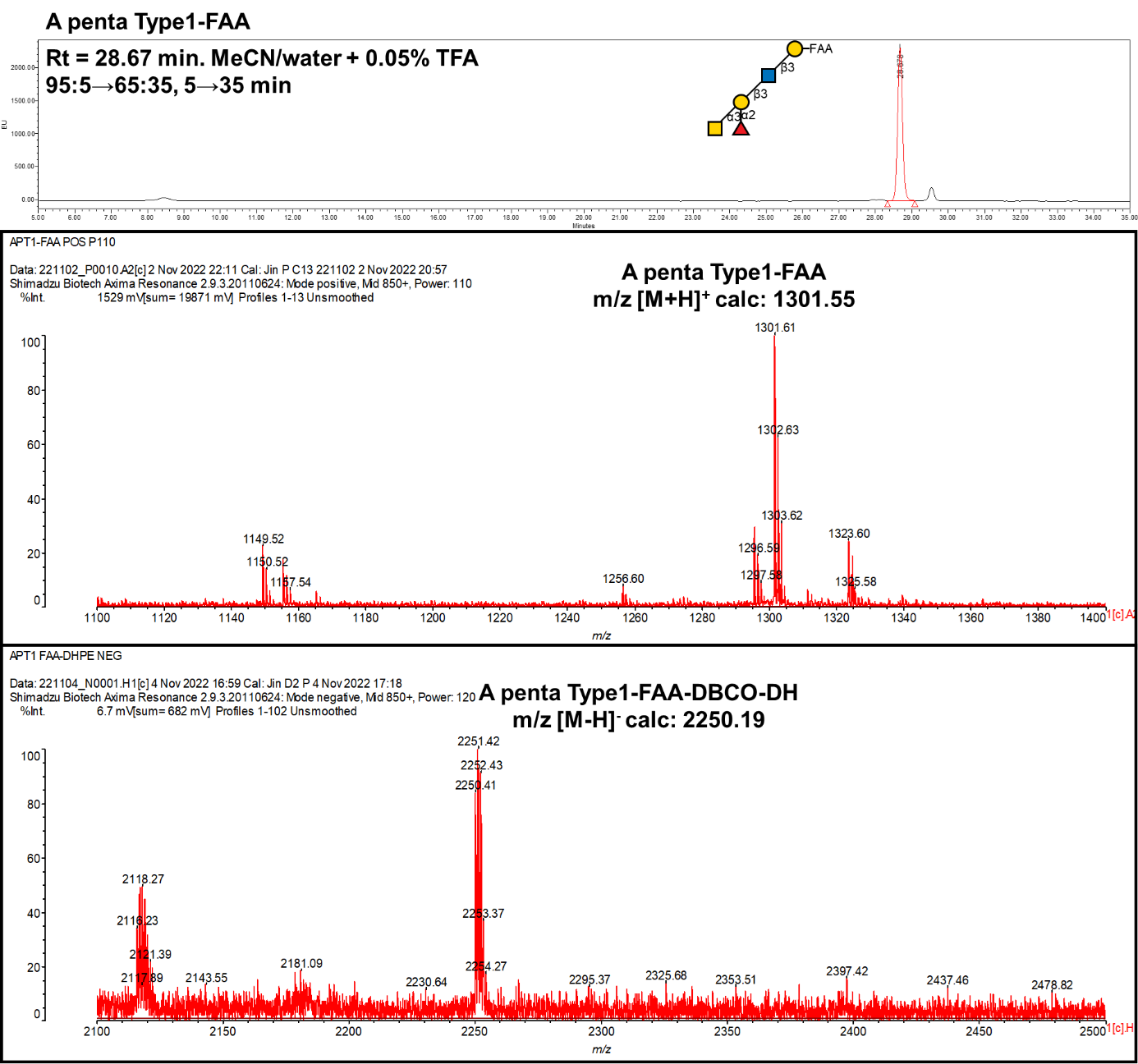

**Probe No 2**

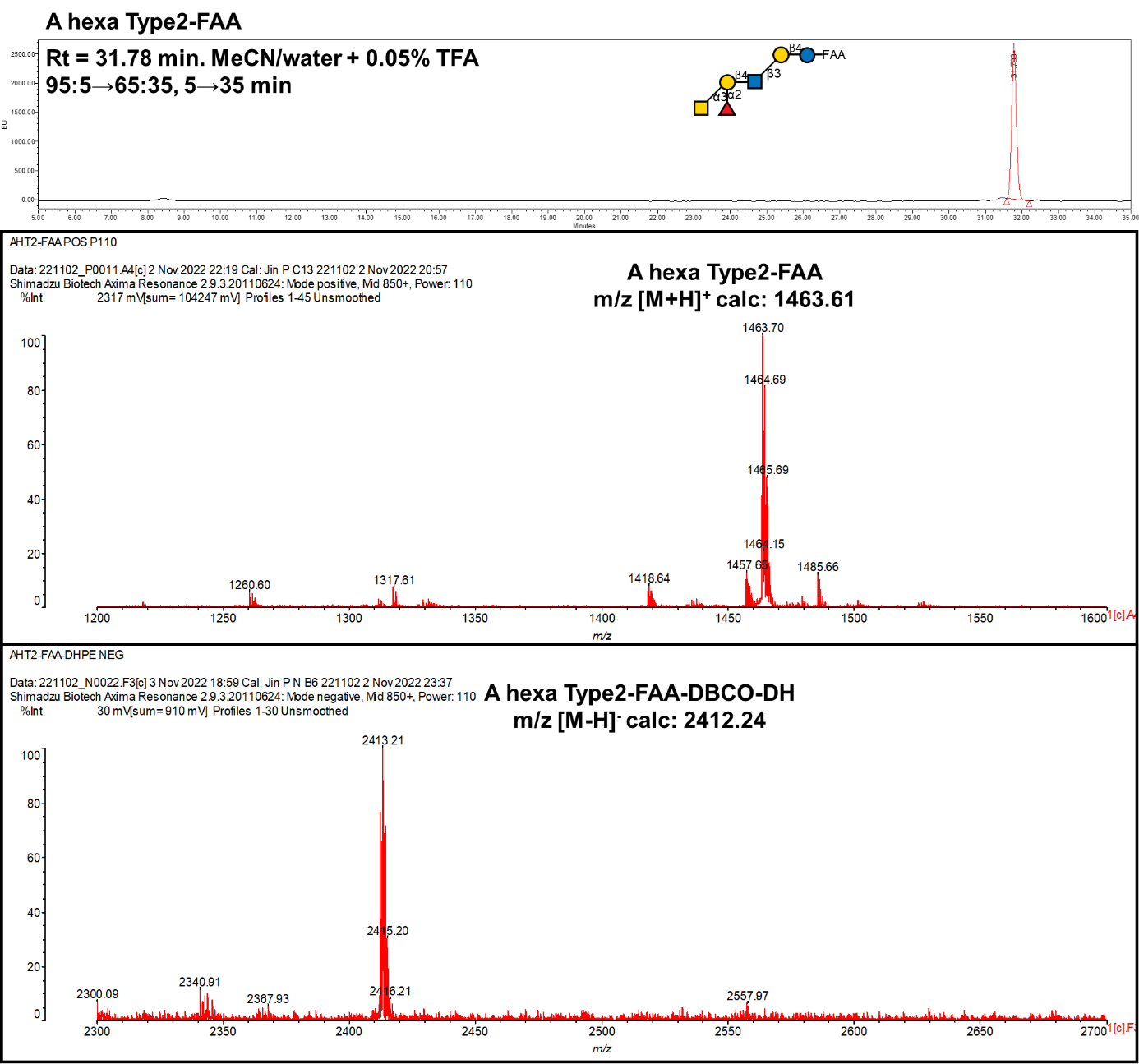

**Probe No 3**

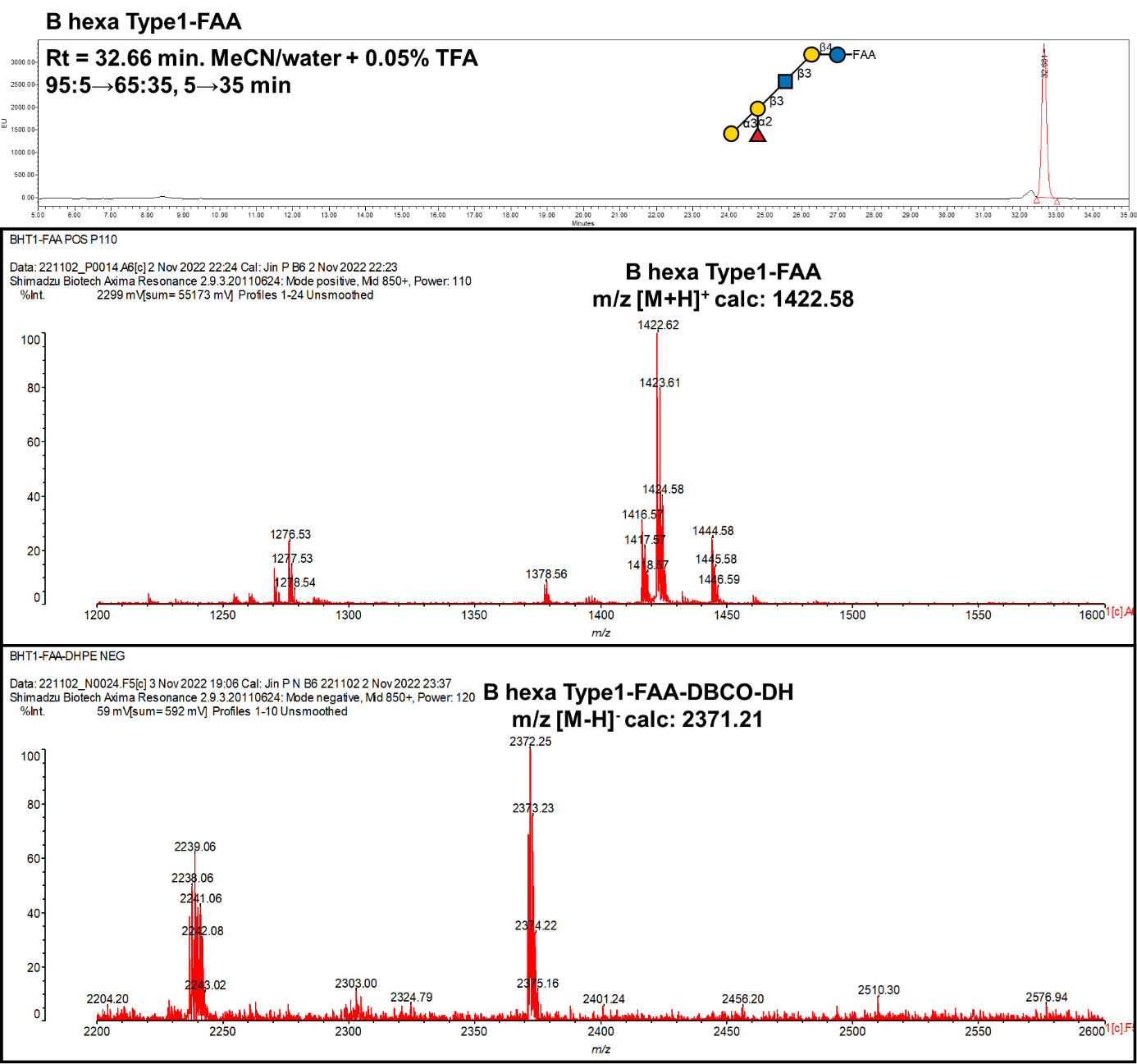

**Probe No 4**

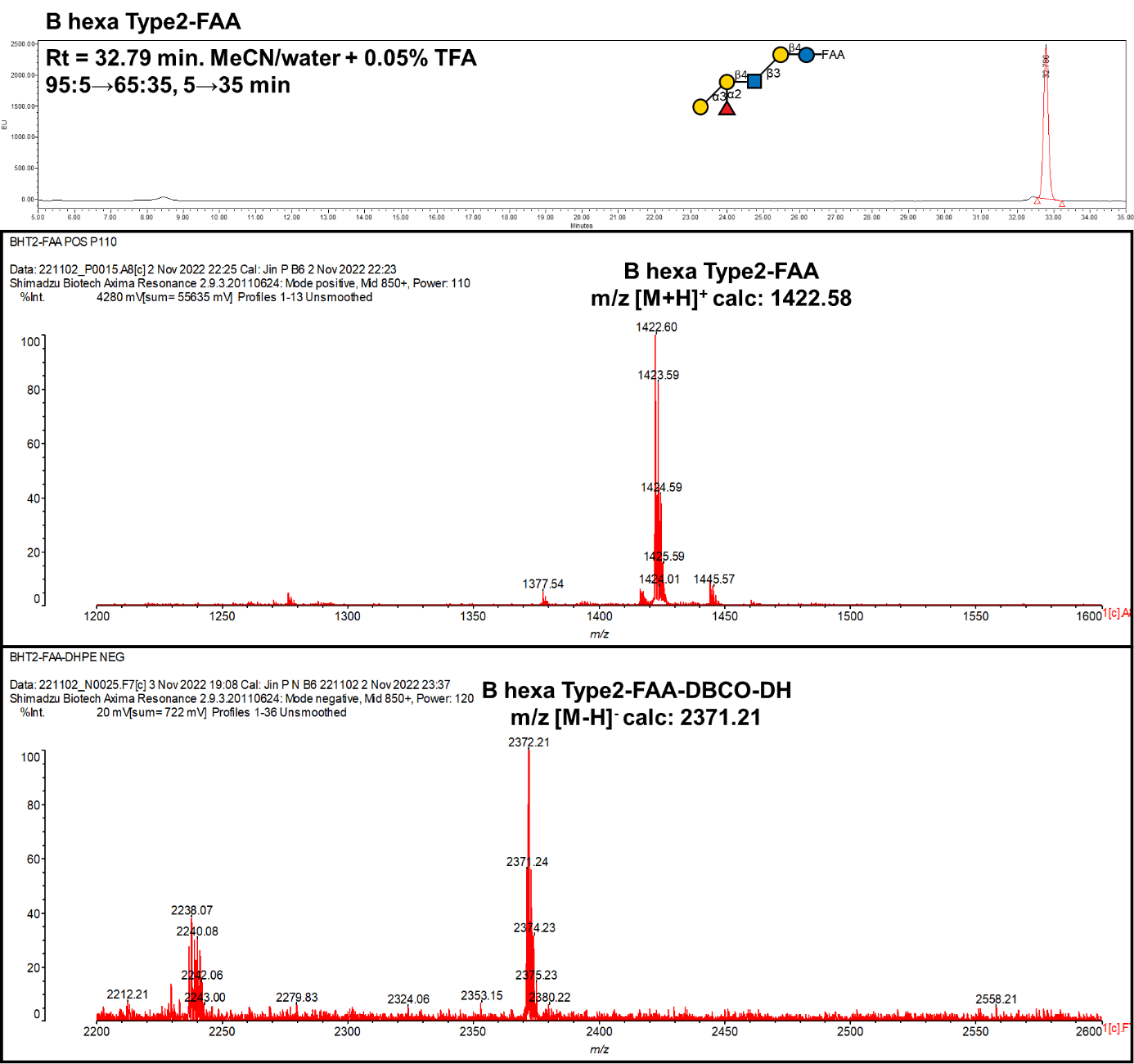

**Probe No 5**

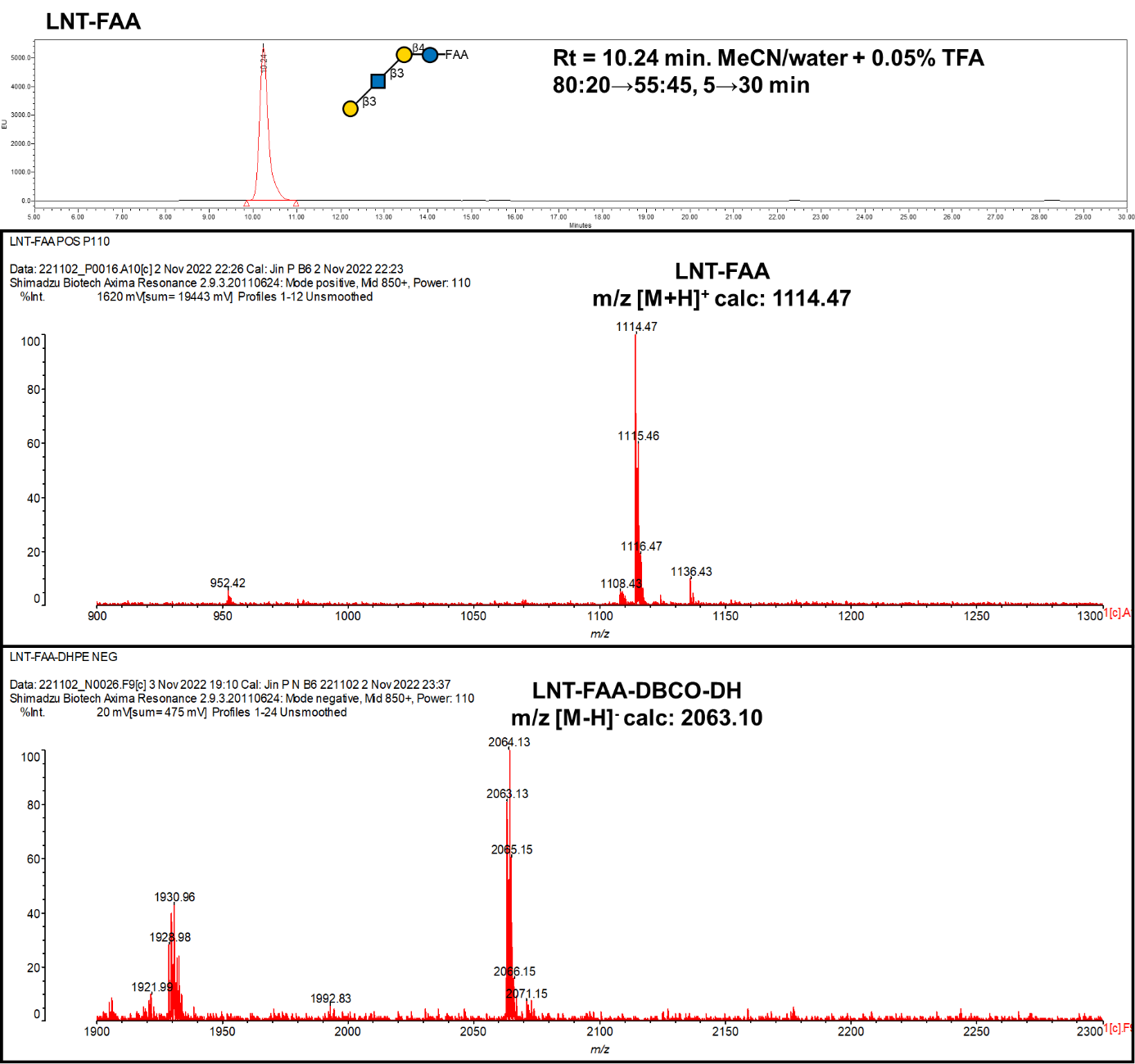

**Probe No 6**

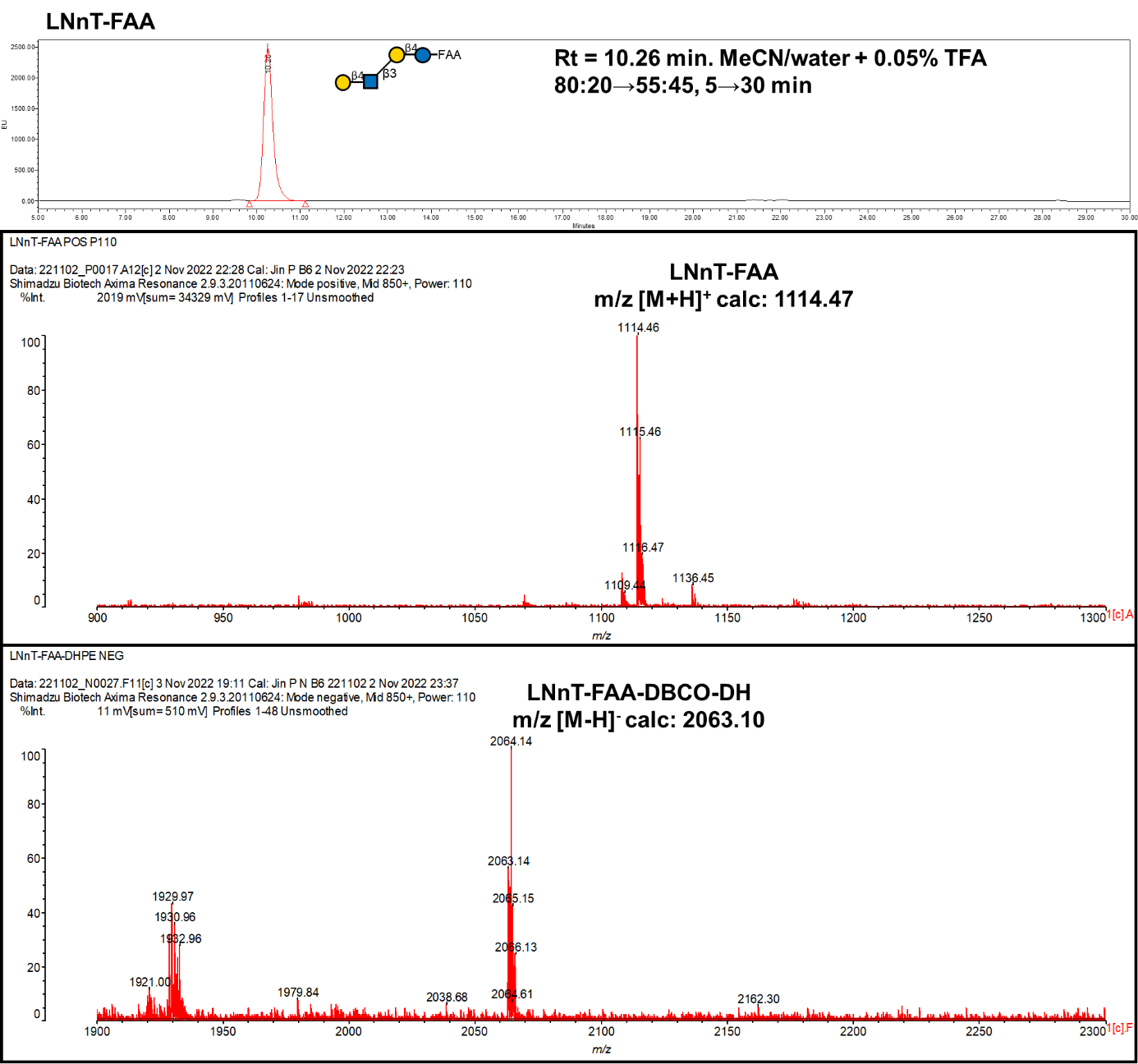

**Probe No 7**

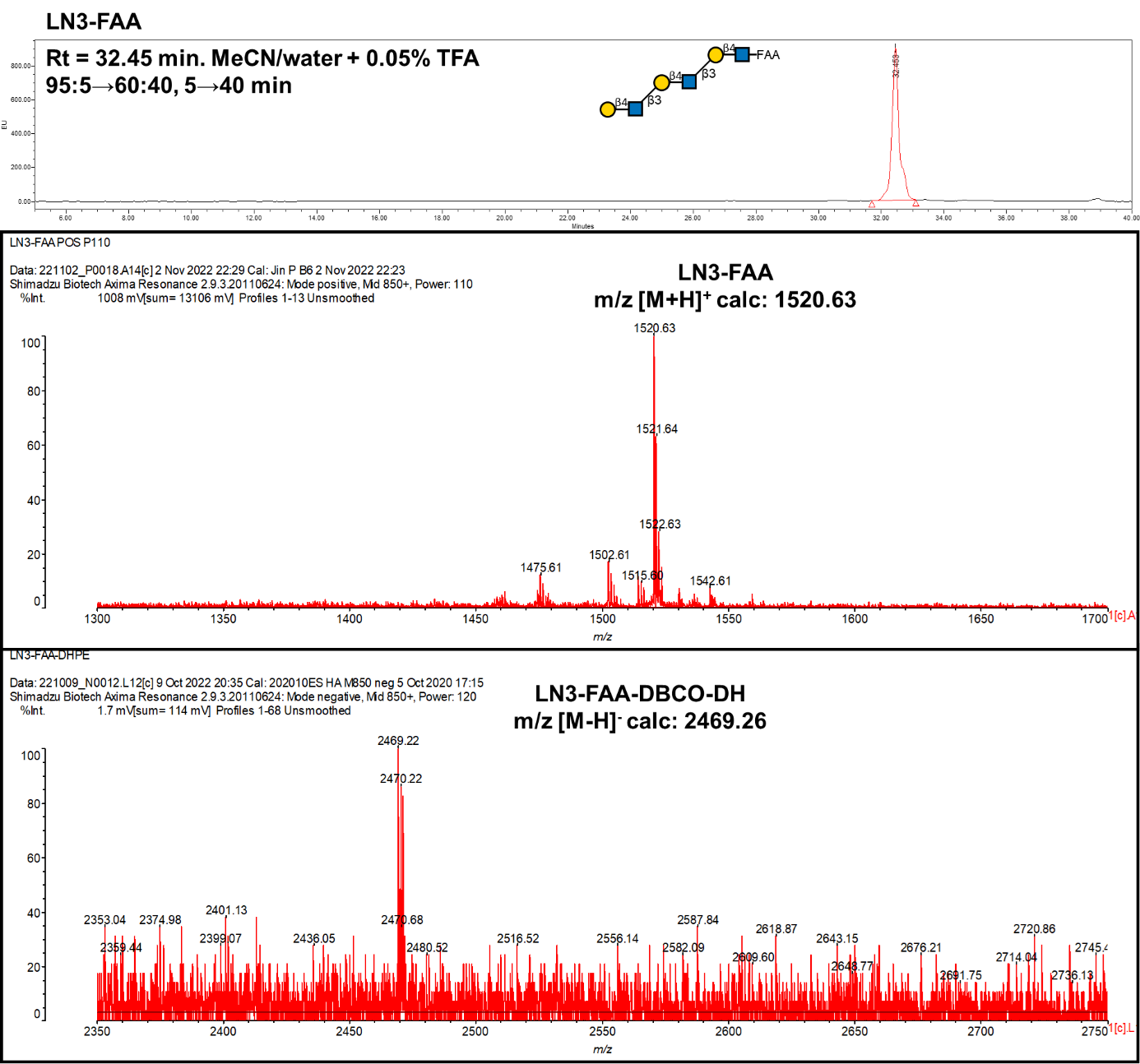

**Probe No 8**

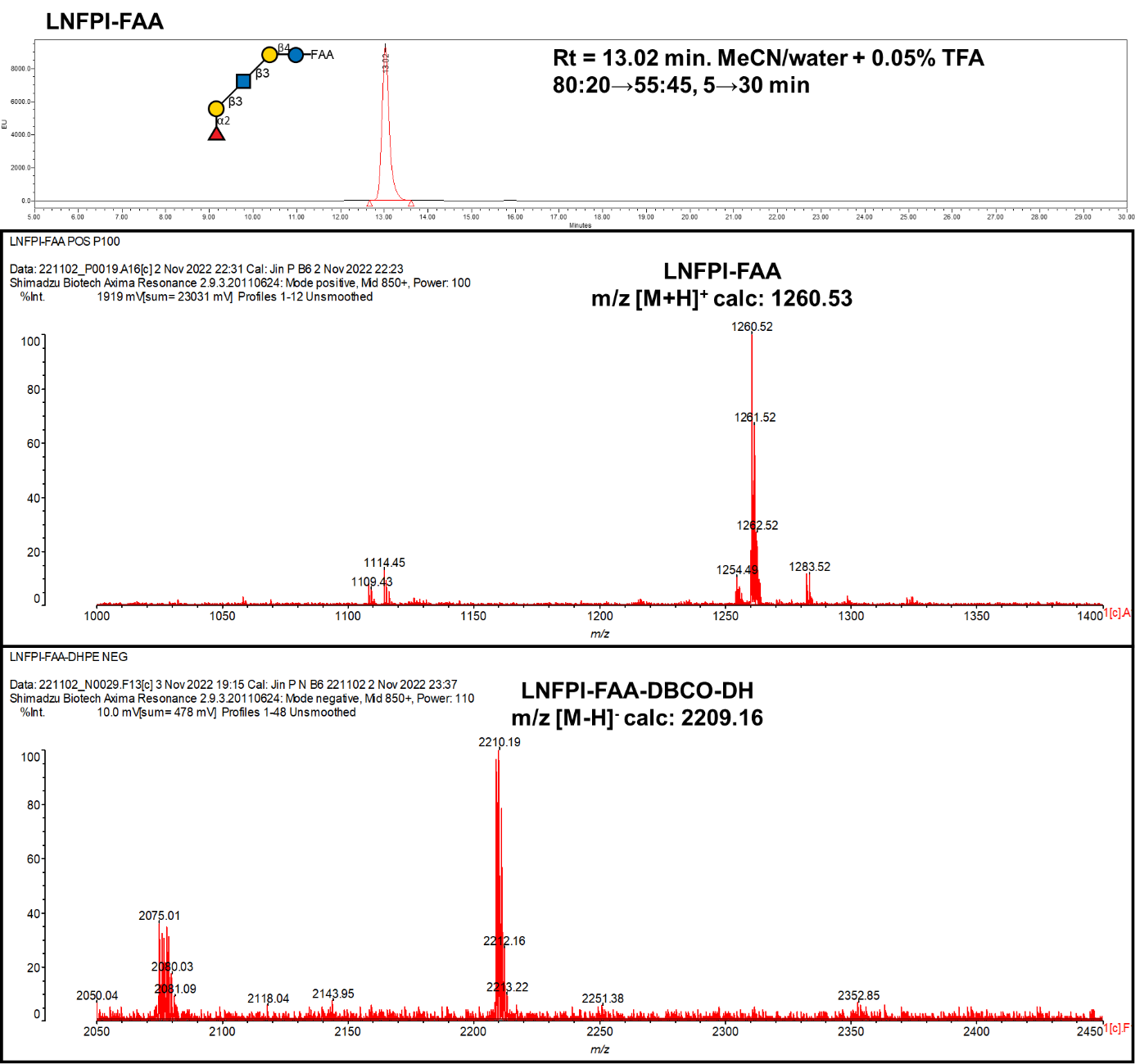

**Probe No 9**

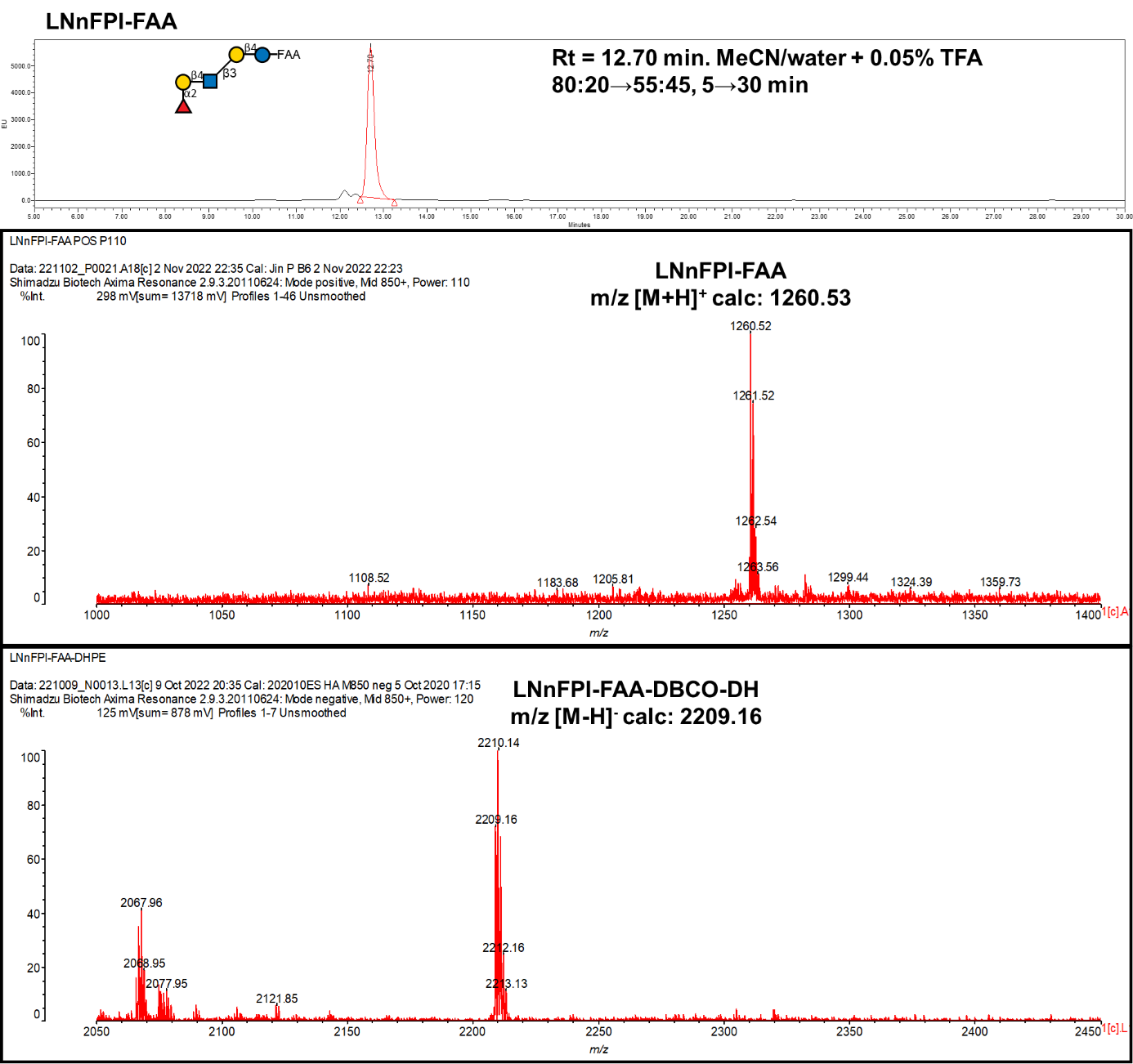

**Probe No 10**

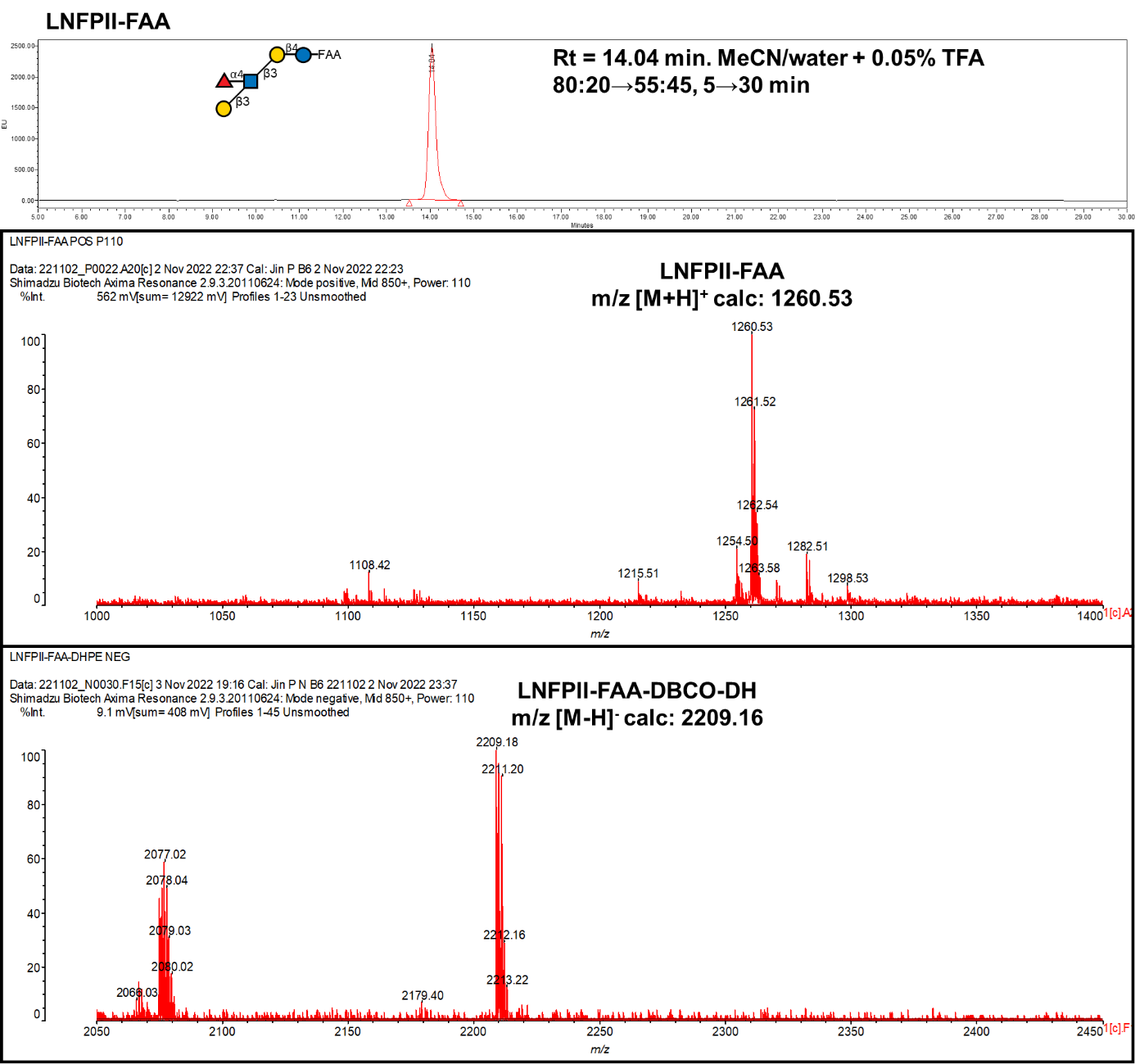

**Probe No 11**

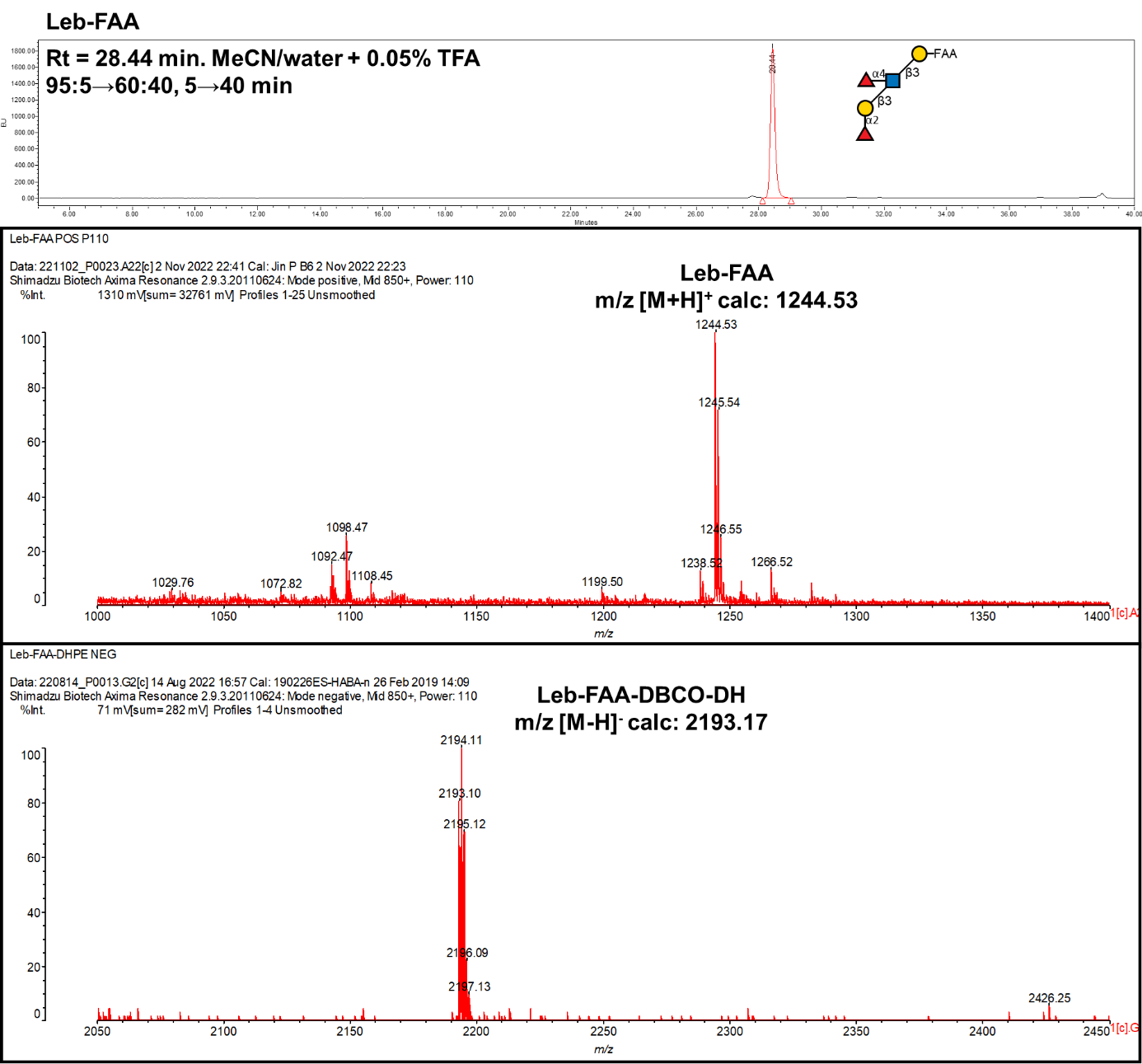

**Probe No 12**

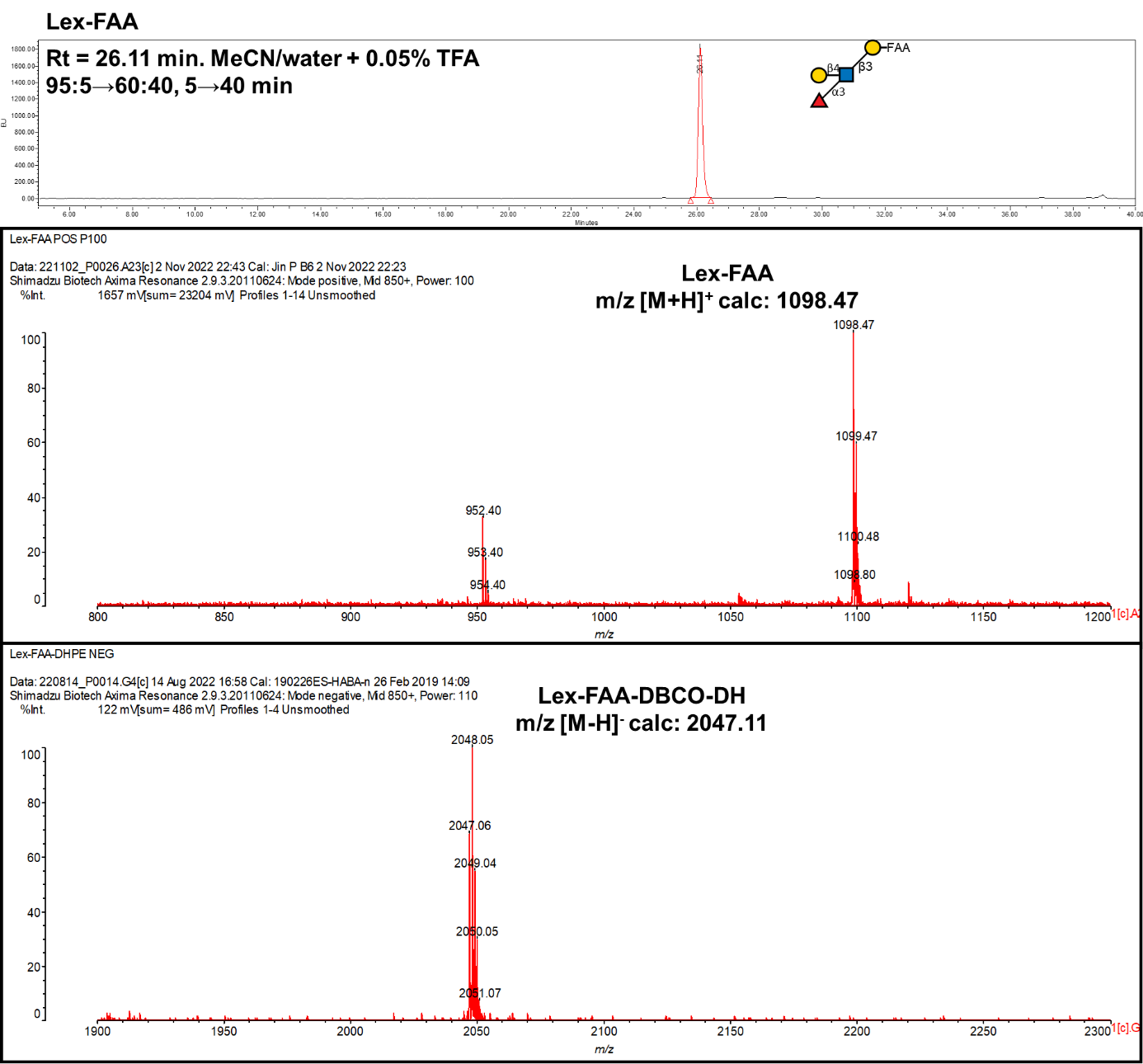

**Probe No 13**

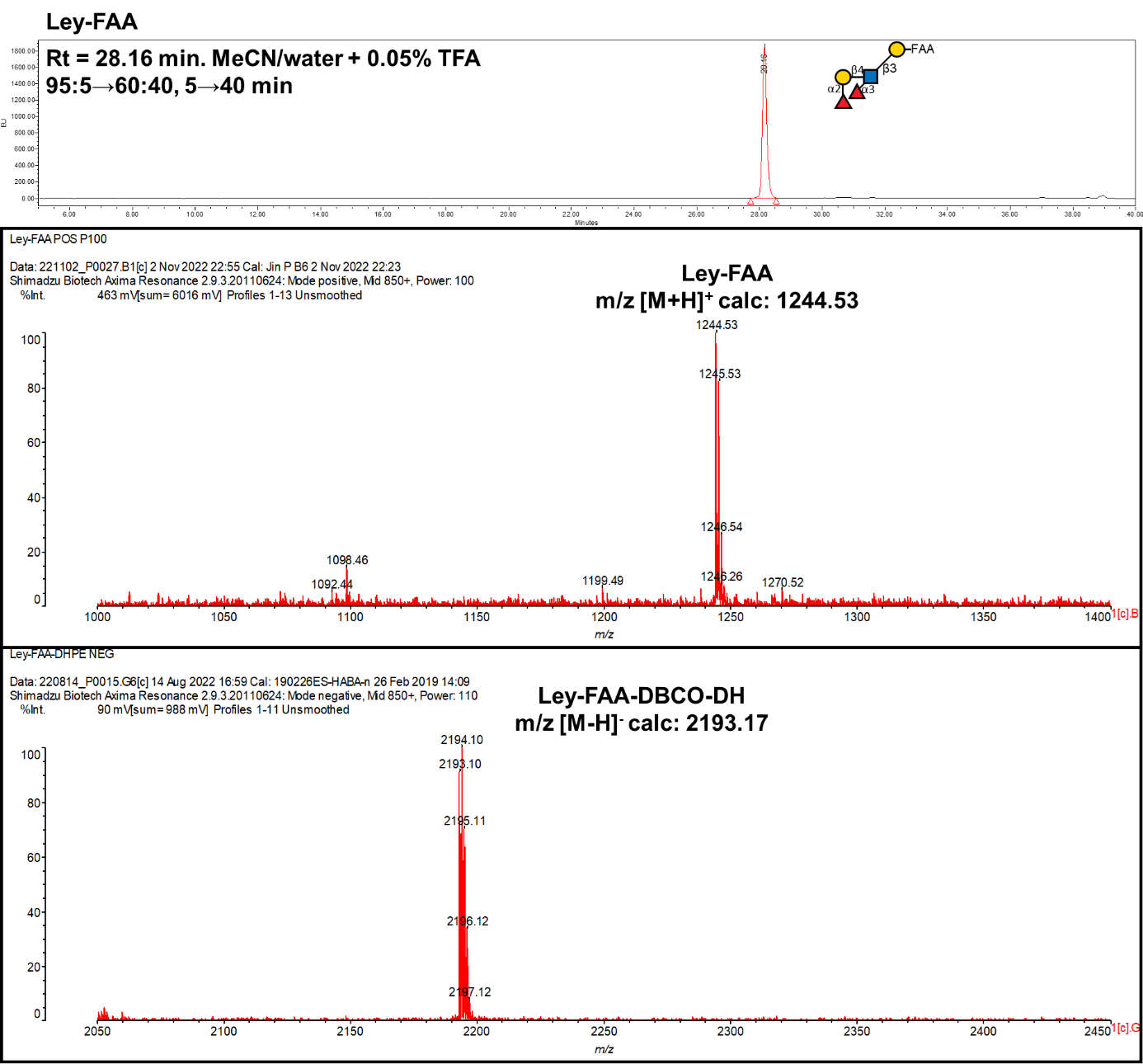

**Probe No 14**

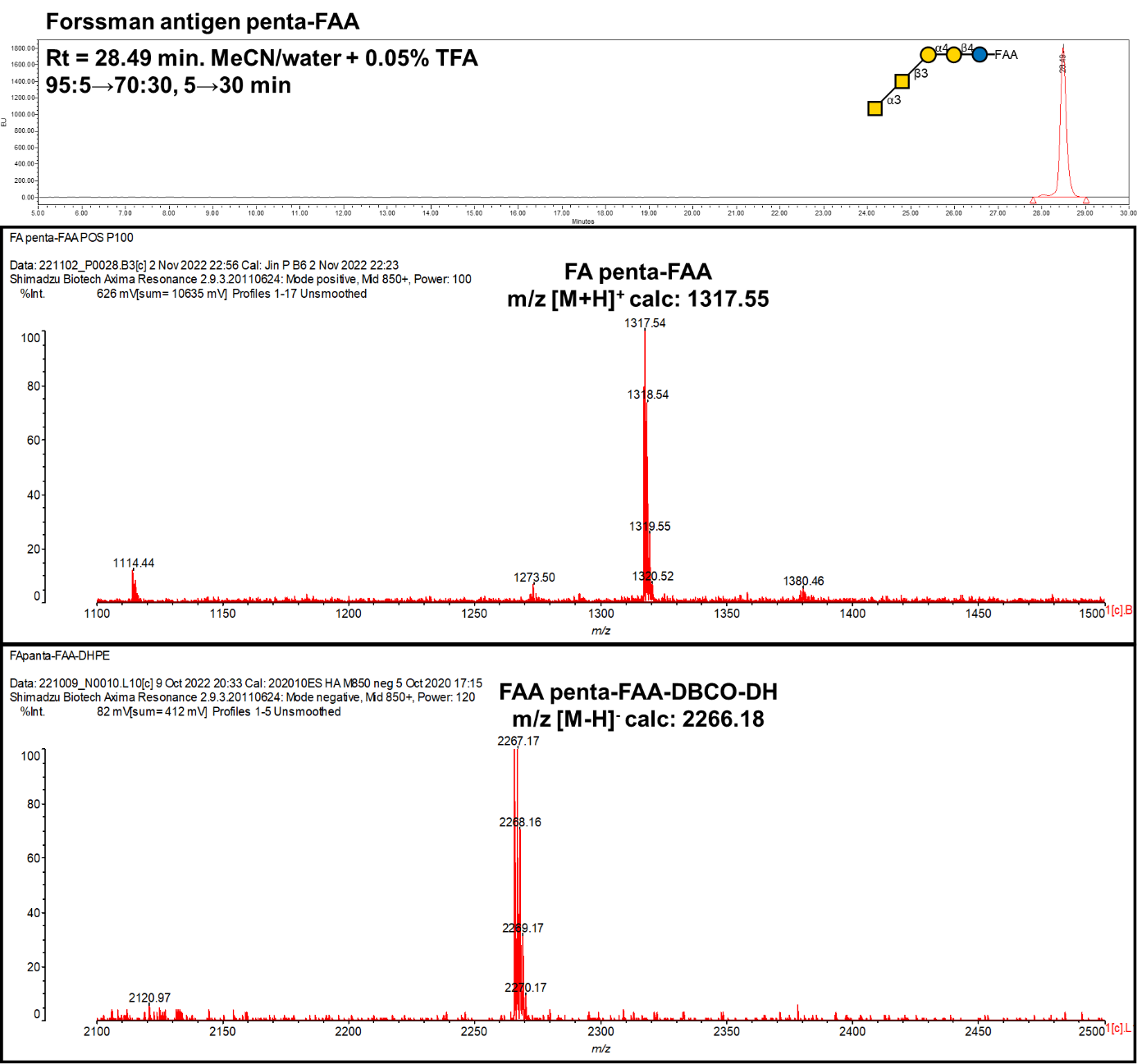

**Probe No 15**

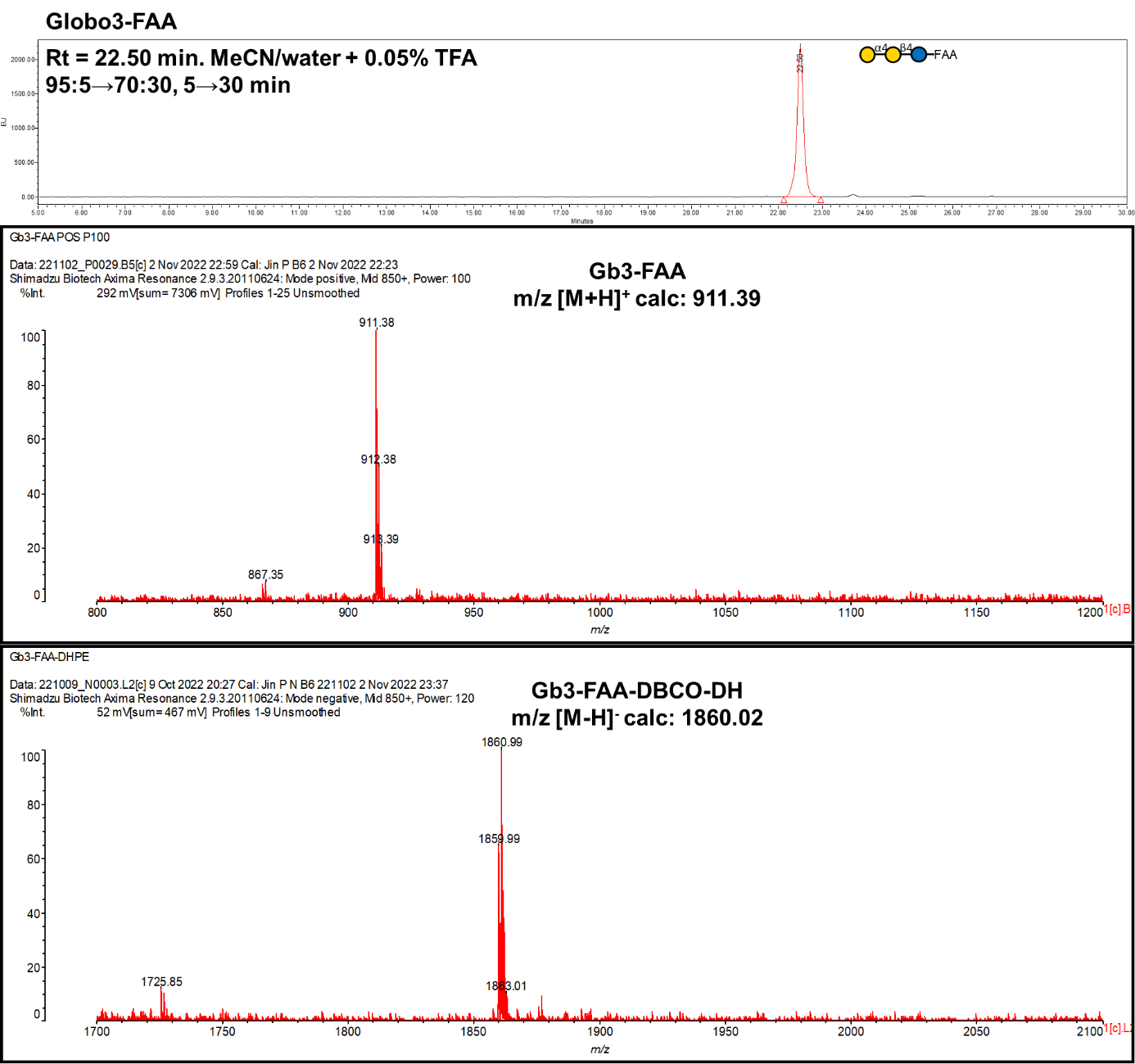

**Probe No 16**

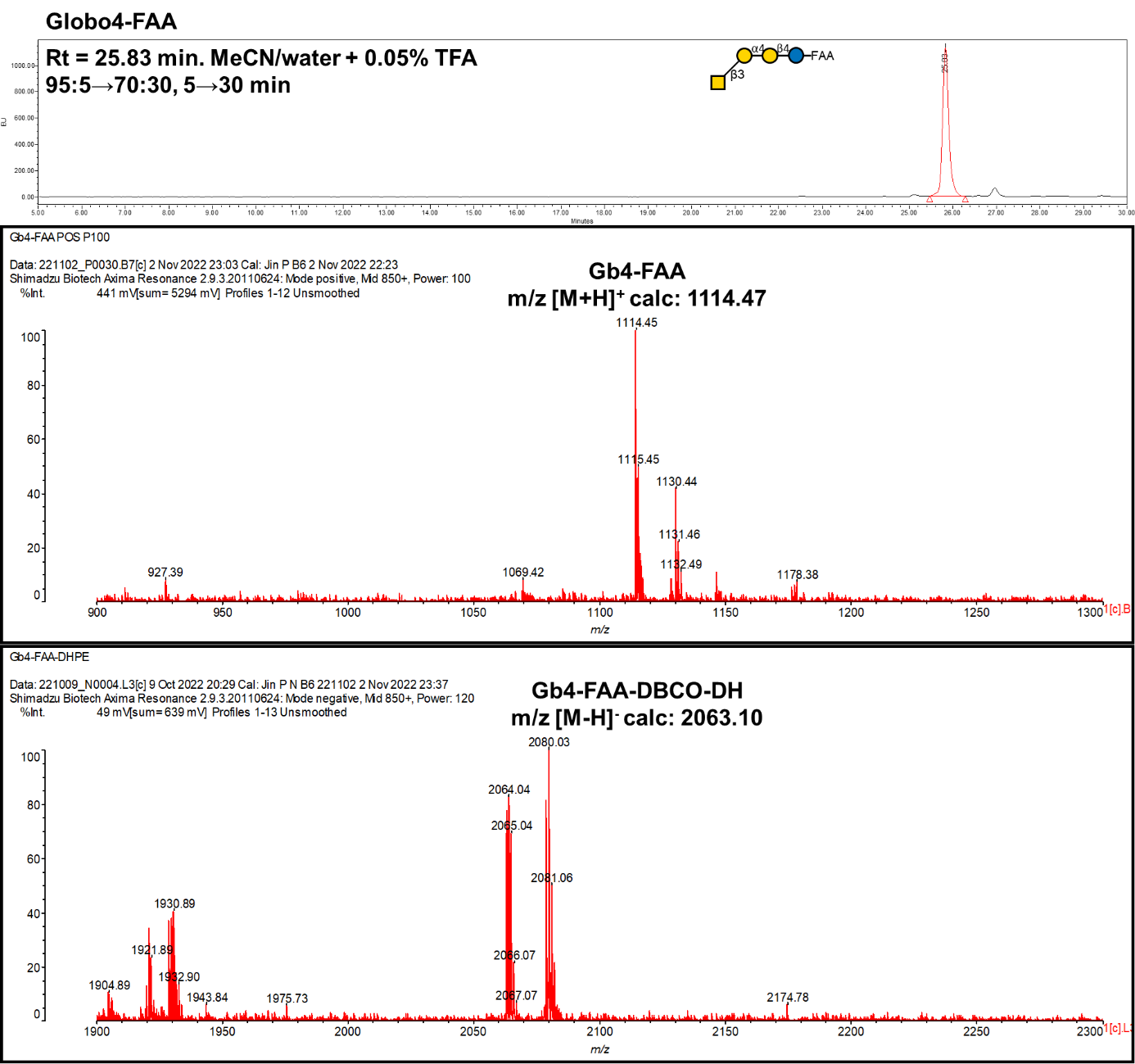

**Probe No 17**

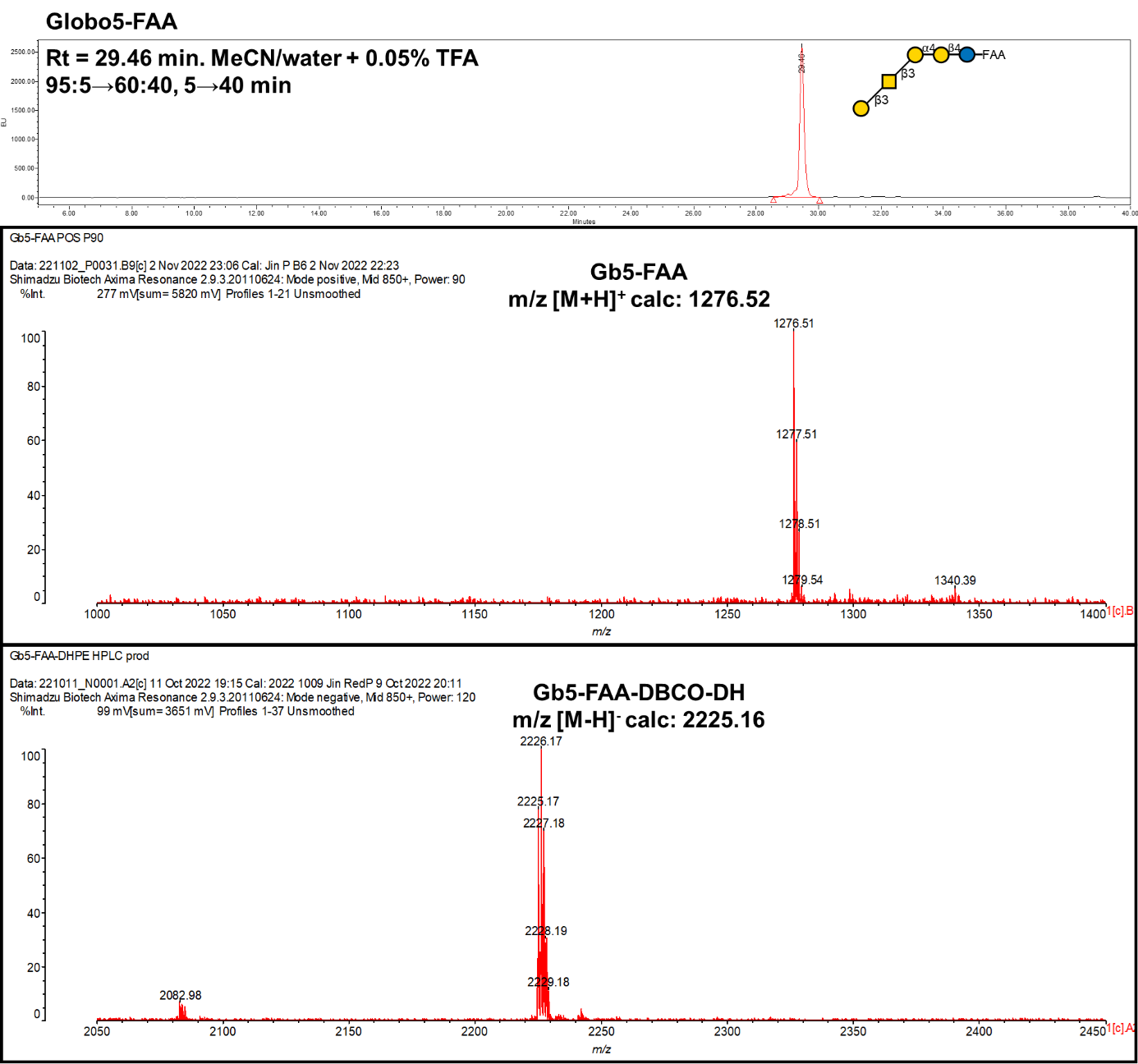

**Probe No 18**

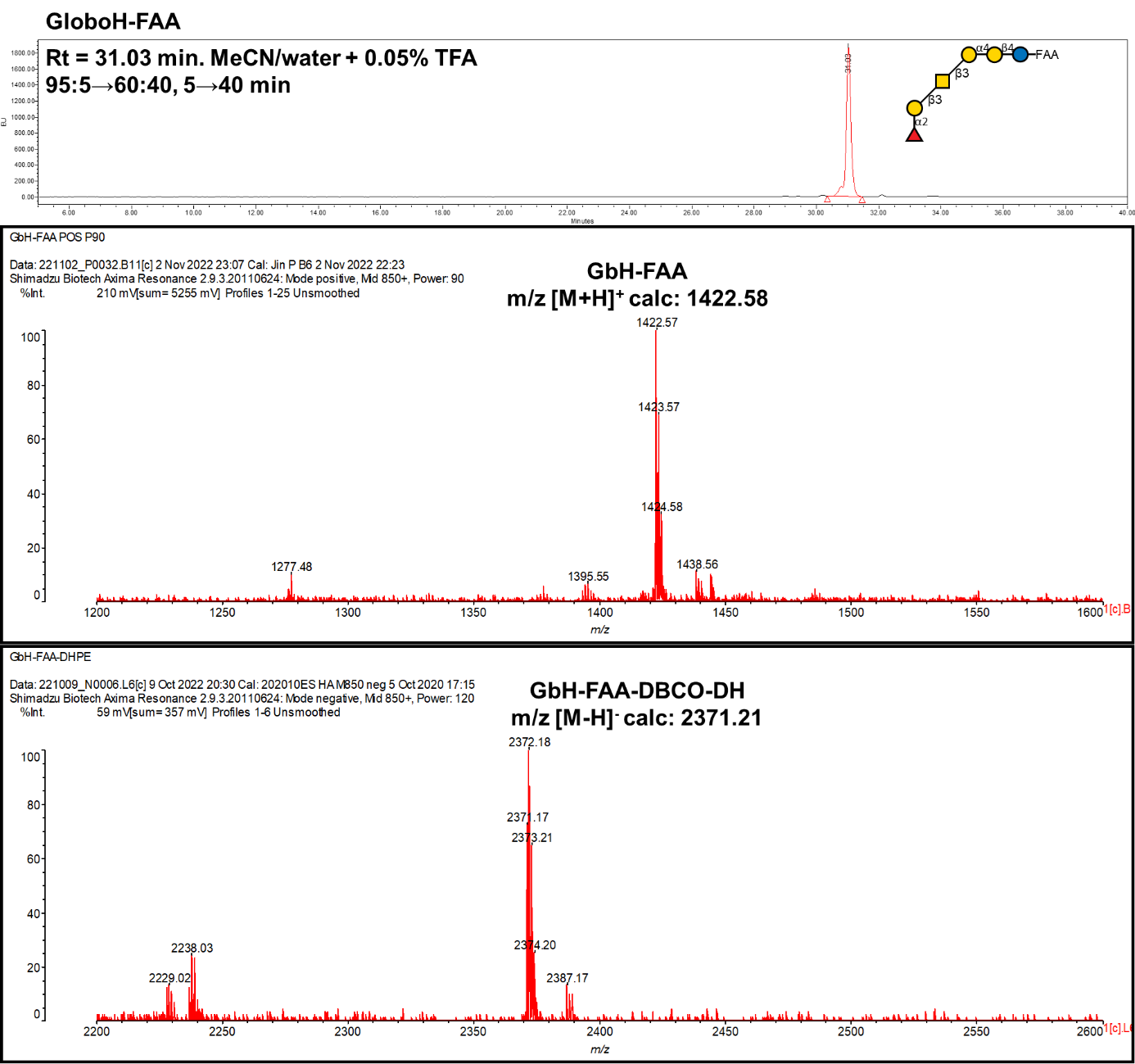

**Probe No 19**

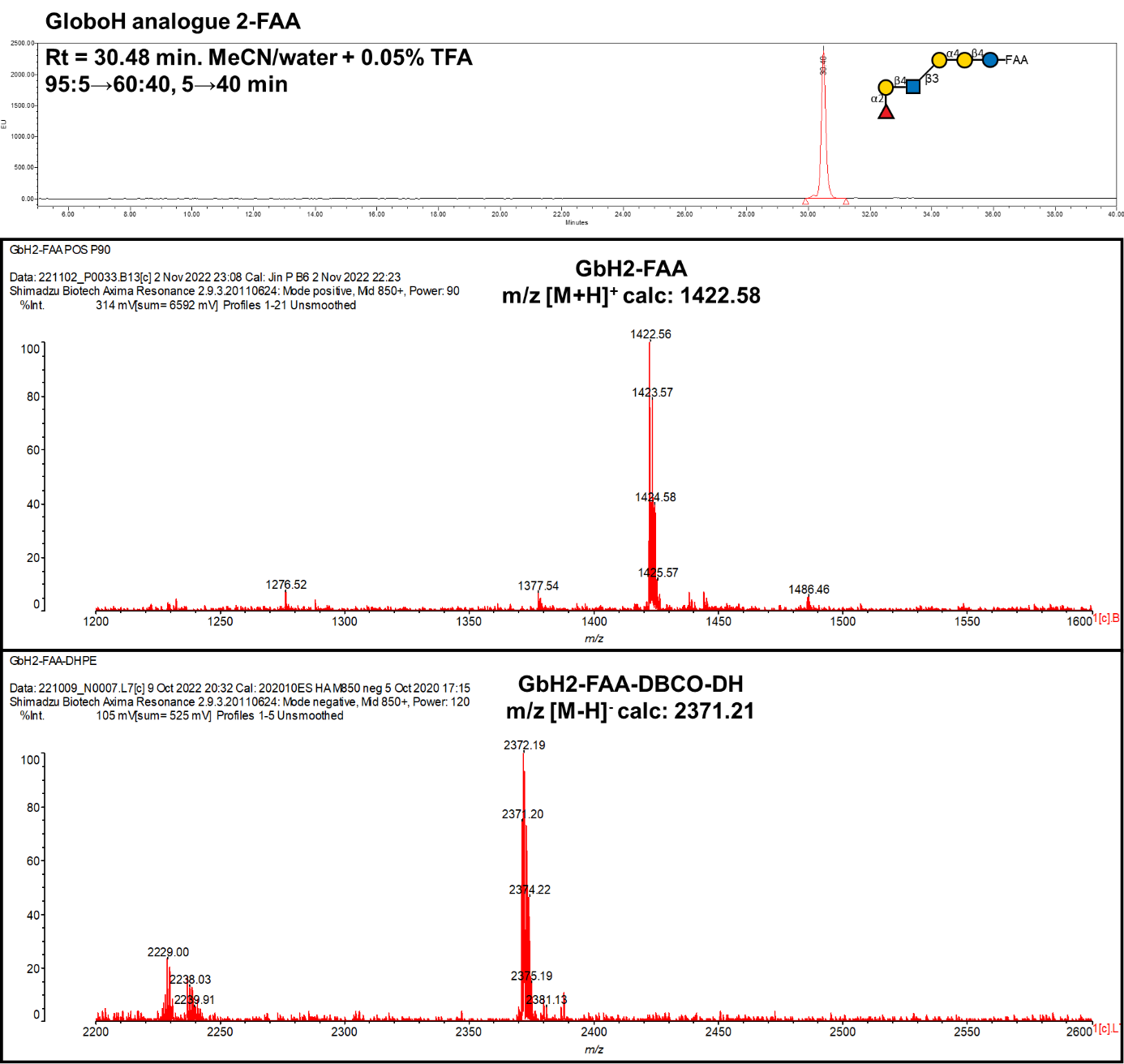

**Probe No 20**

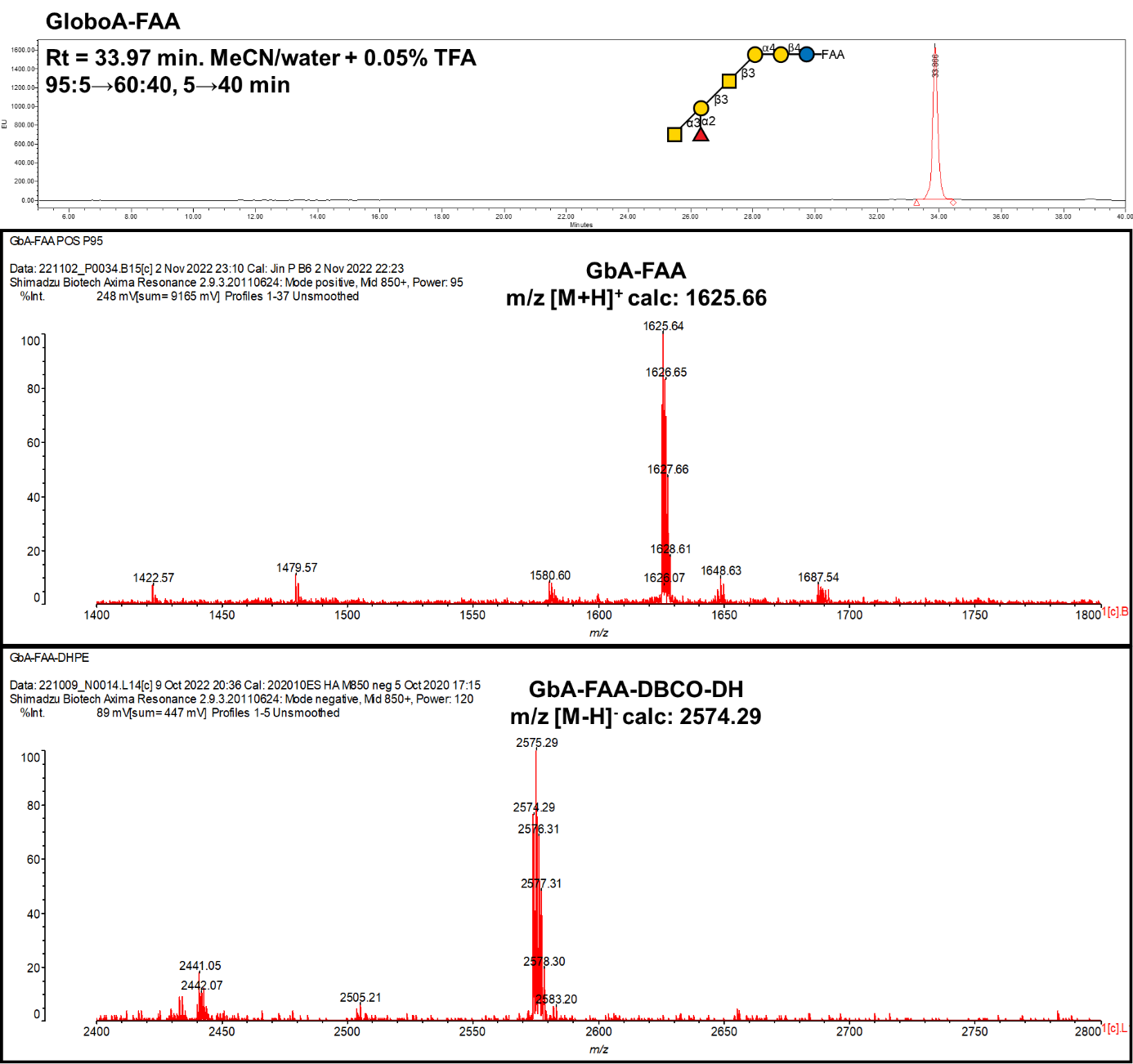

**Probe No 21**

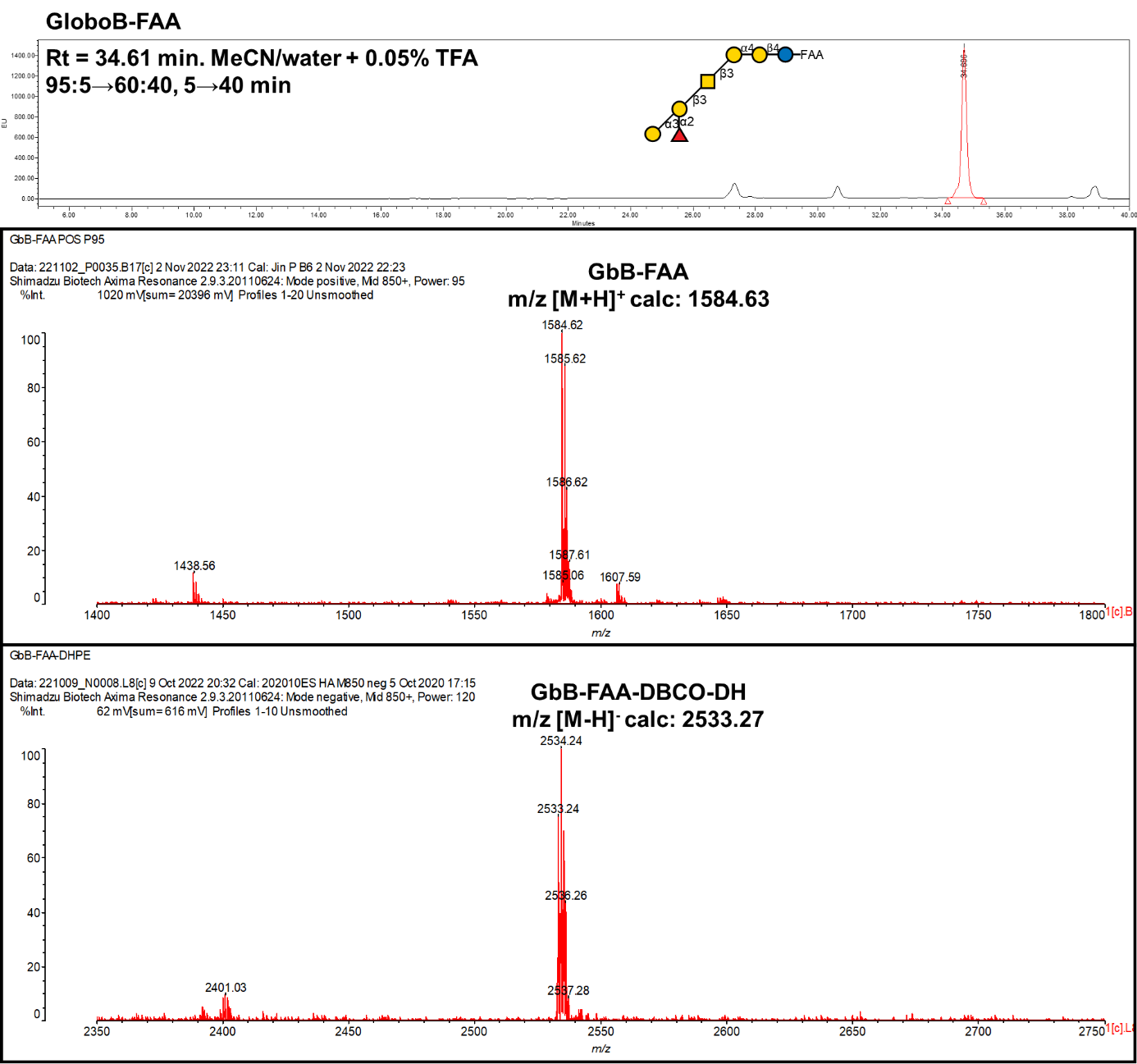

**Probe No 22**

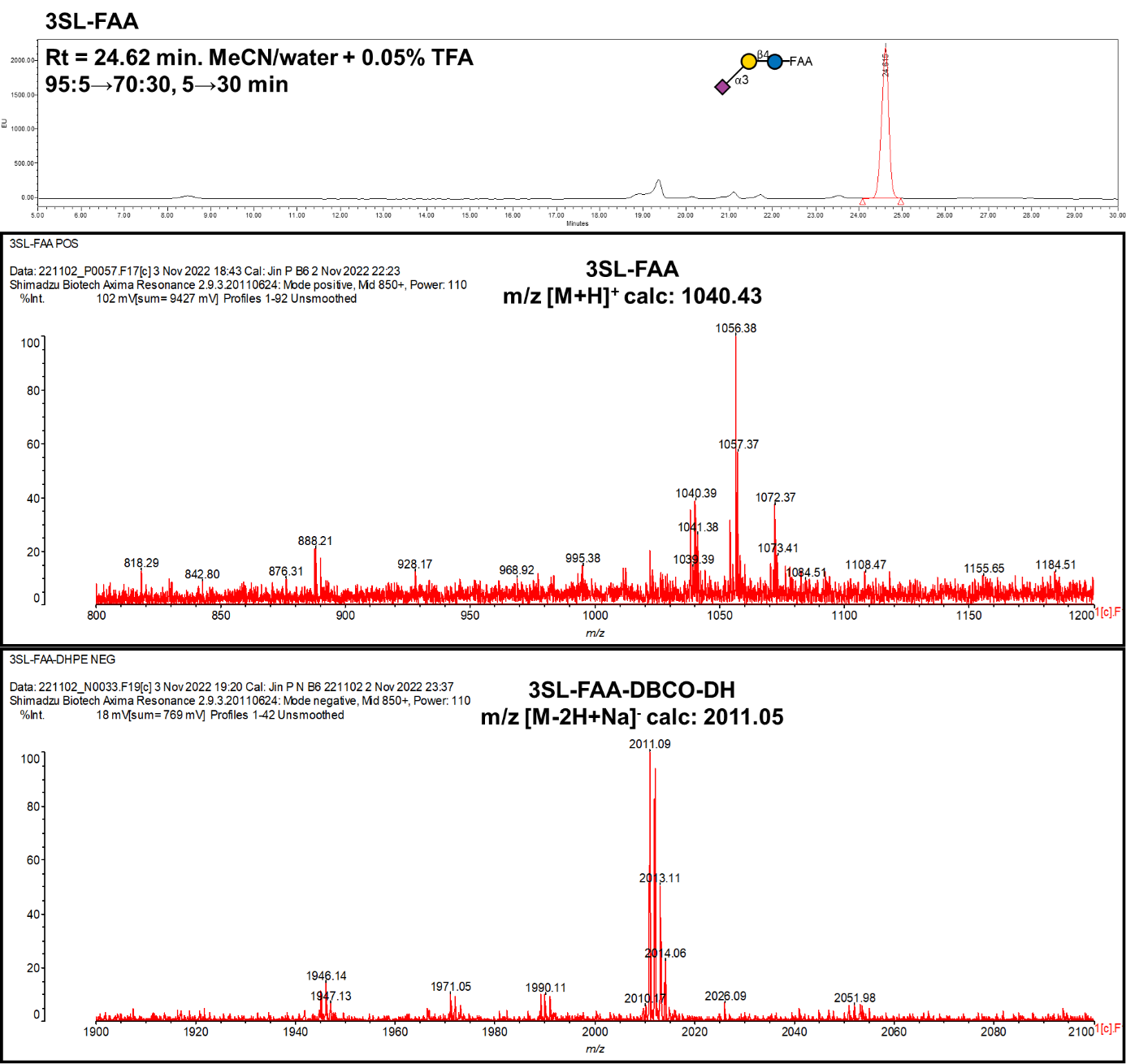

**Probe No 23**

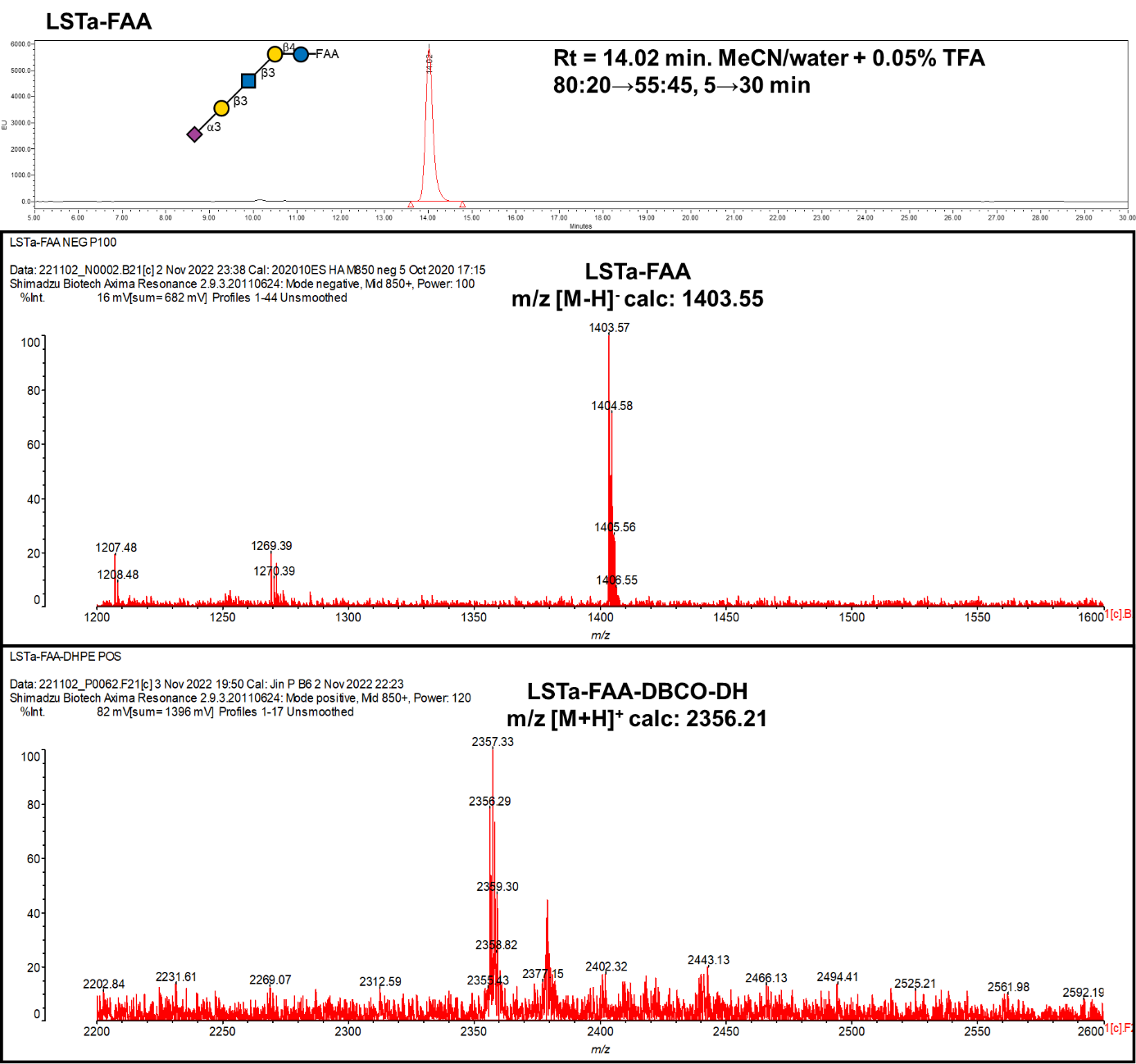

**Probe No 24**

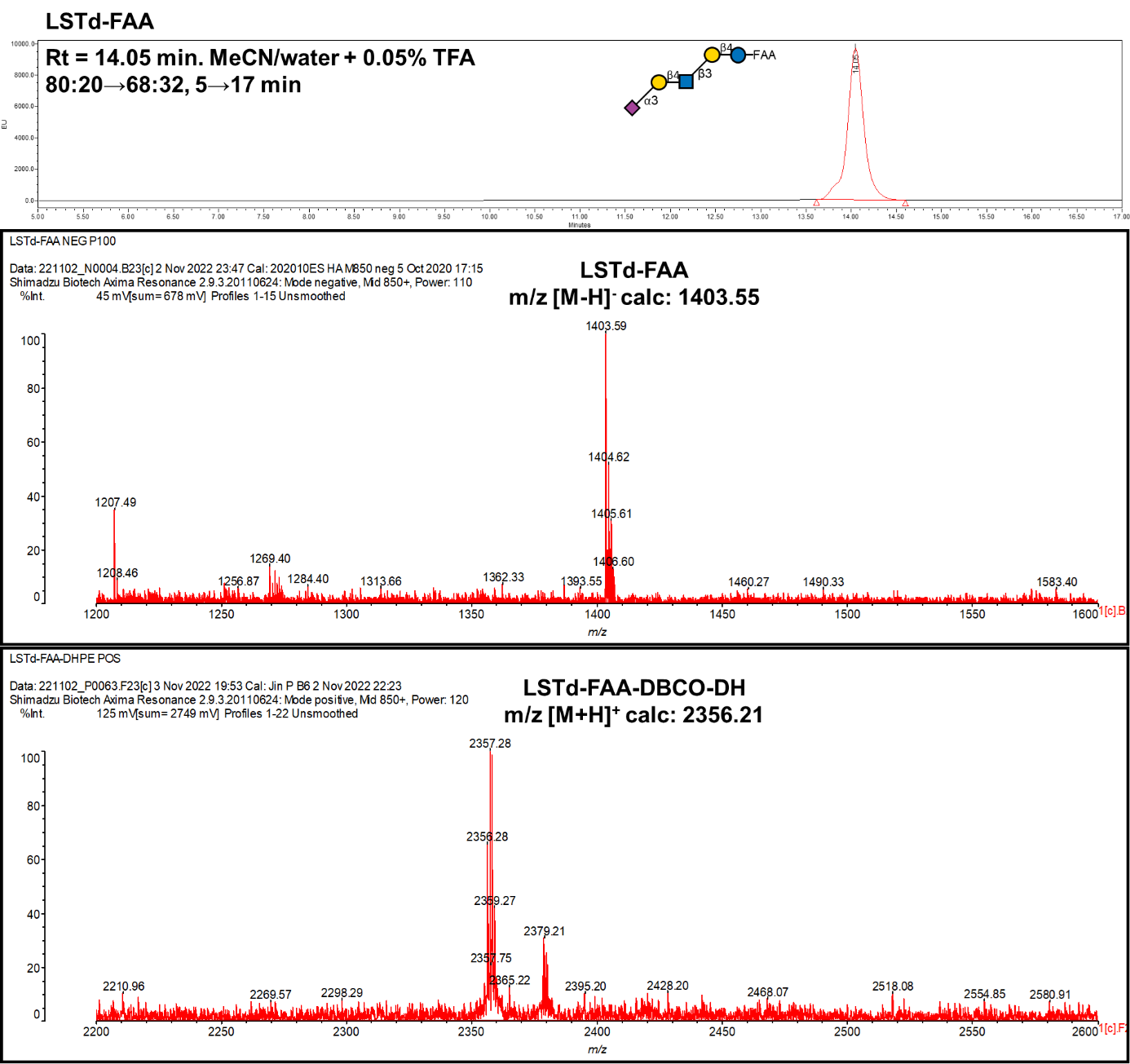

**Probe No 25**

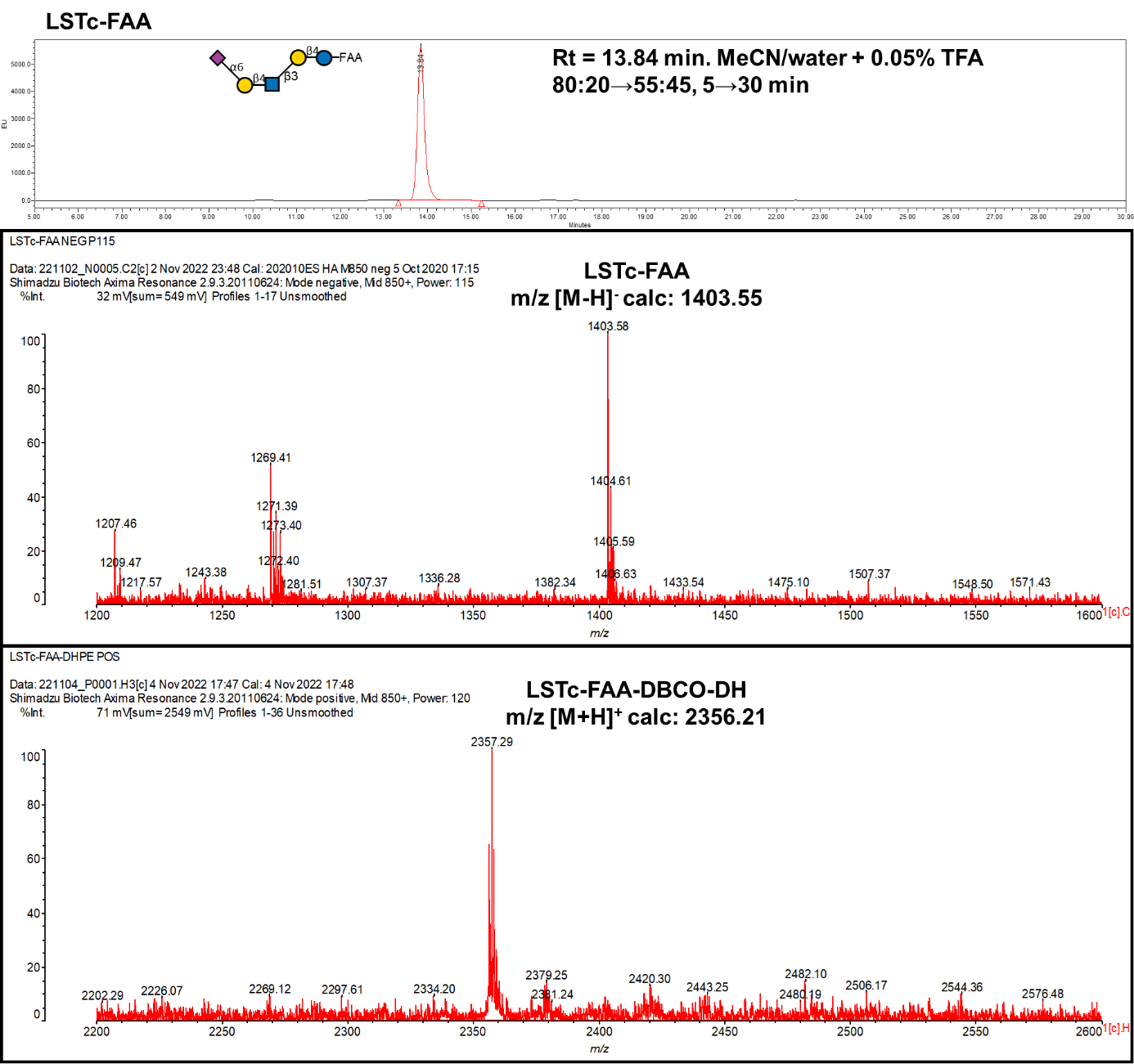

**Probe No 26**

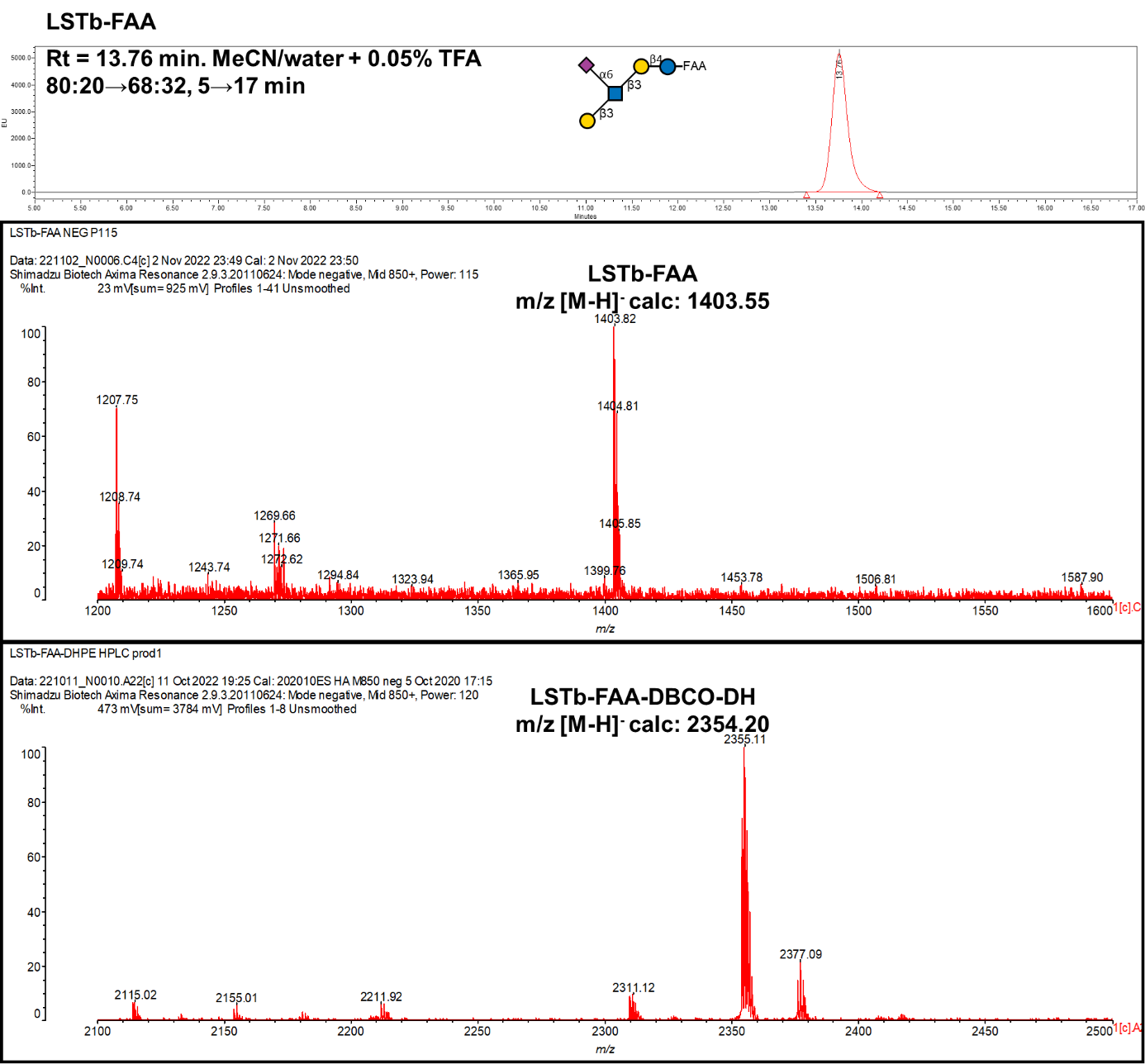

**Probe No 27**

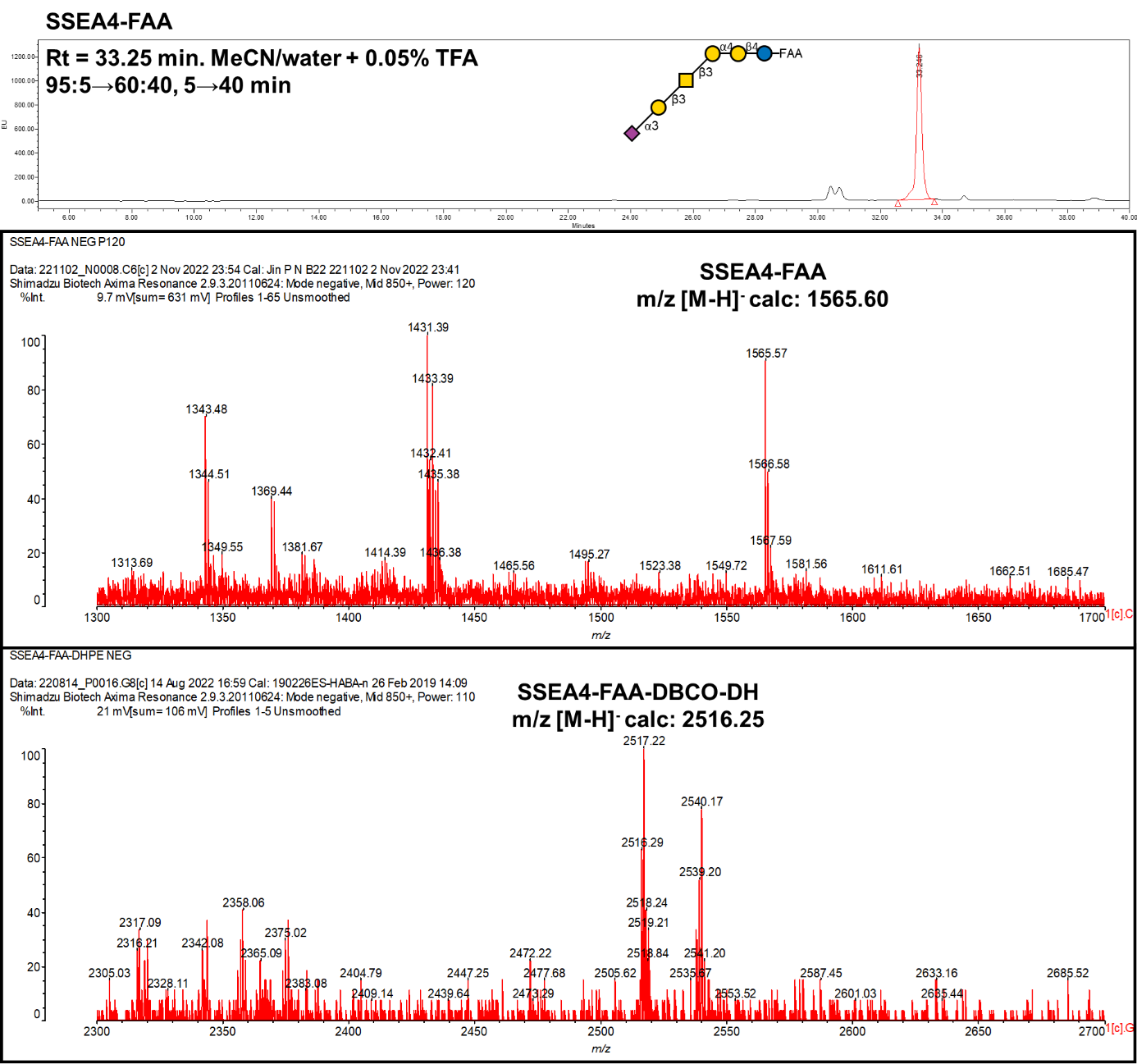

**Probe No 28**

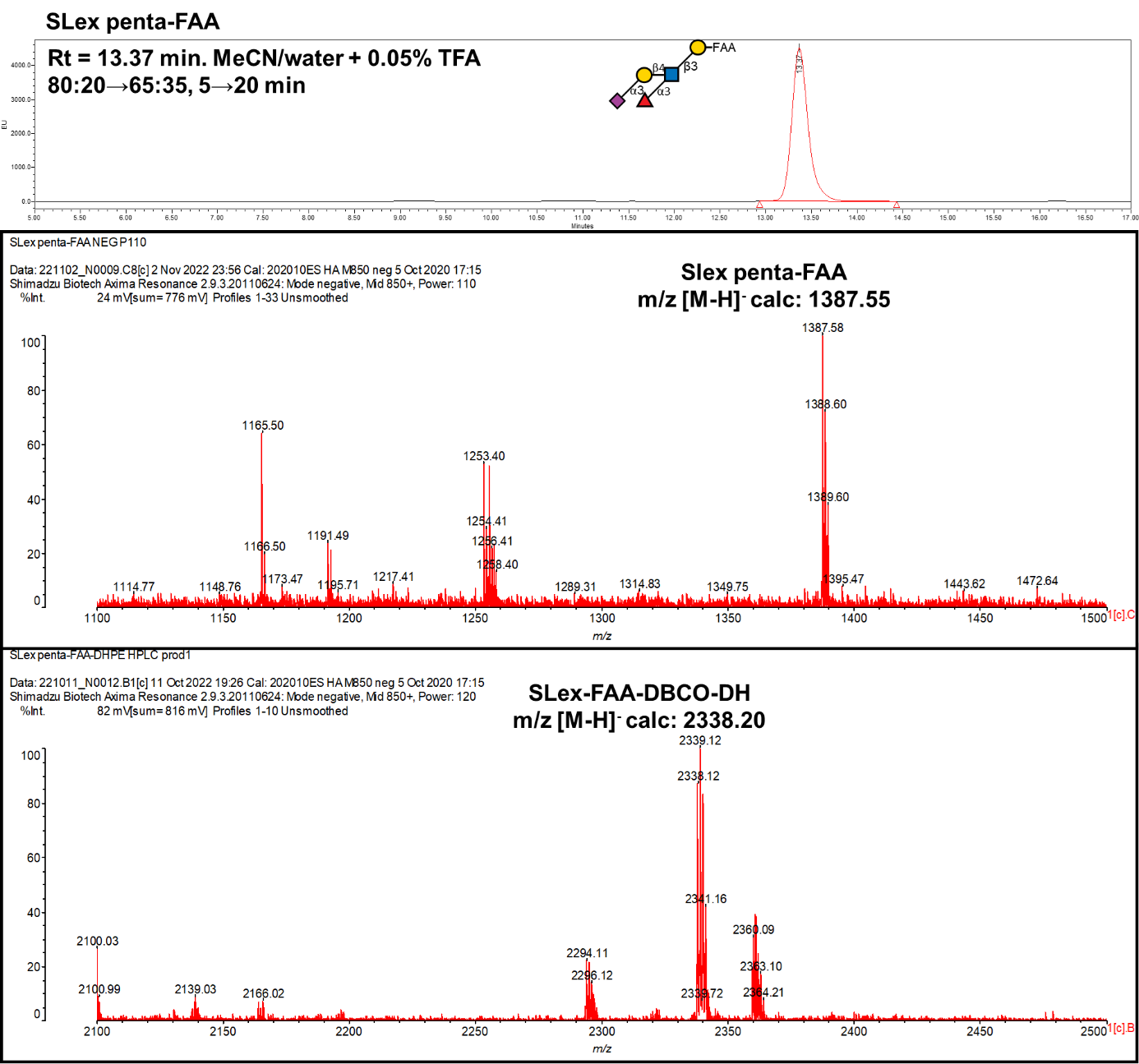

**Probe No 29**

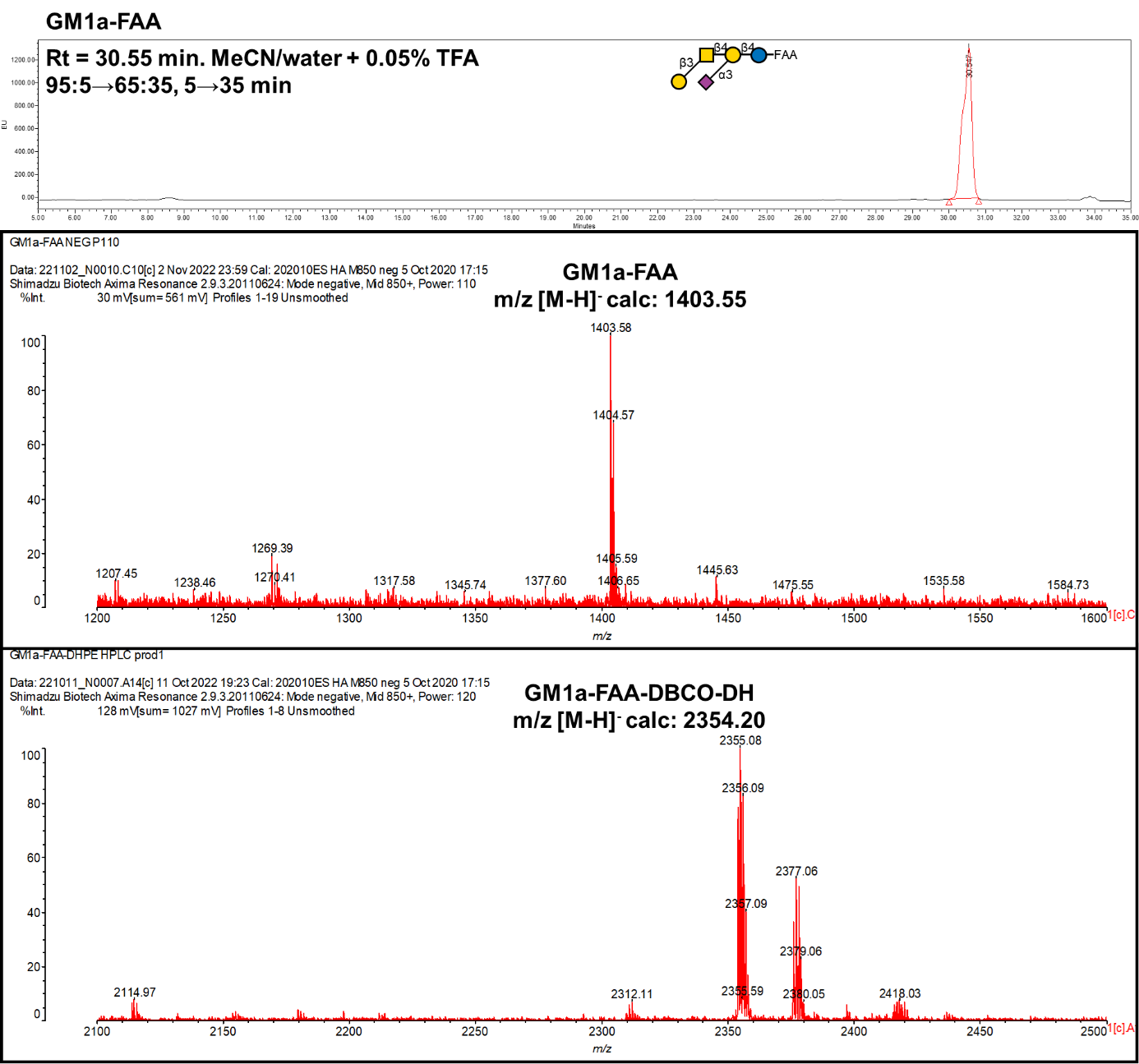

**Probe No 30**

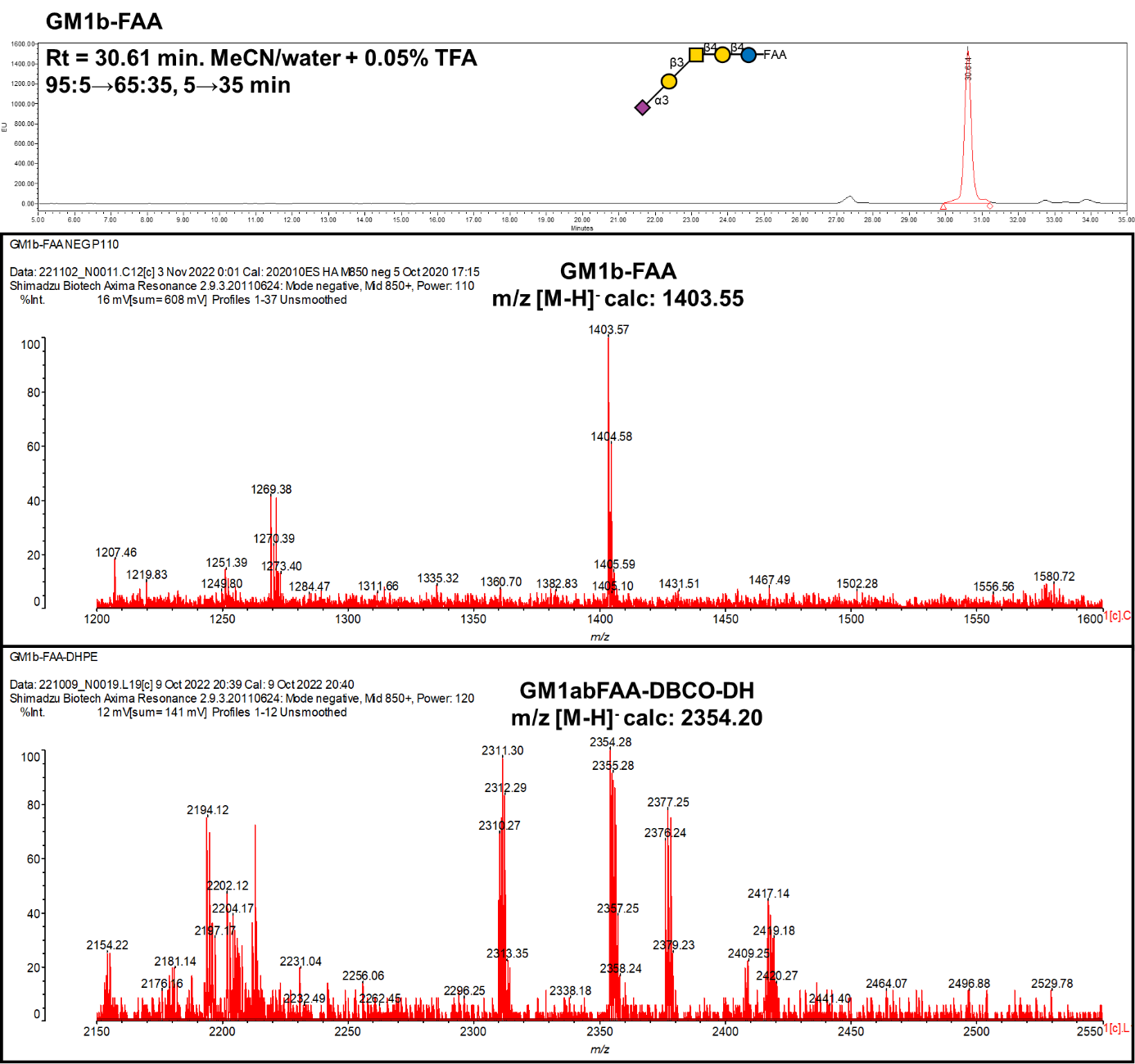

**Probe No 31**

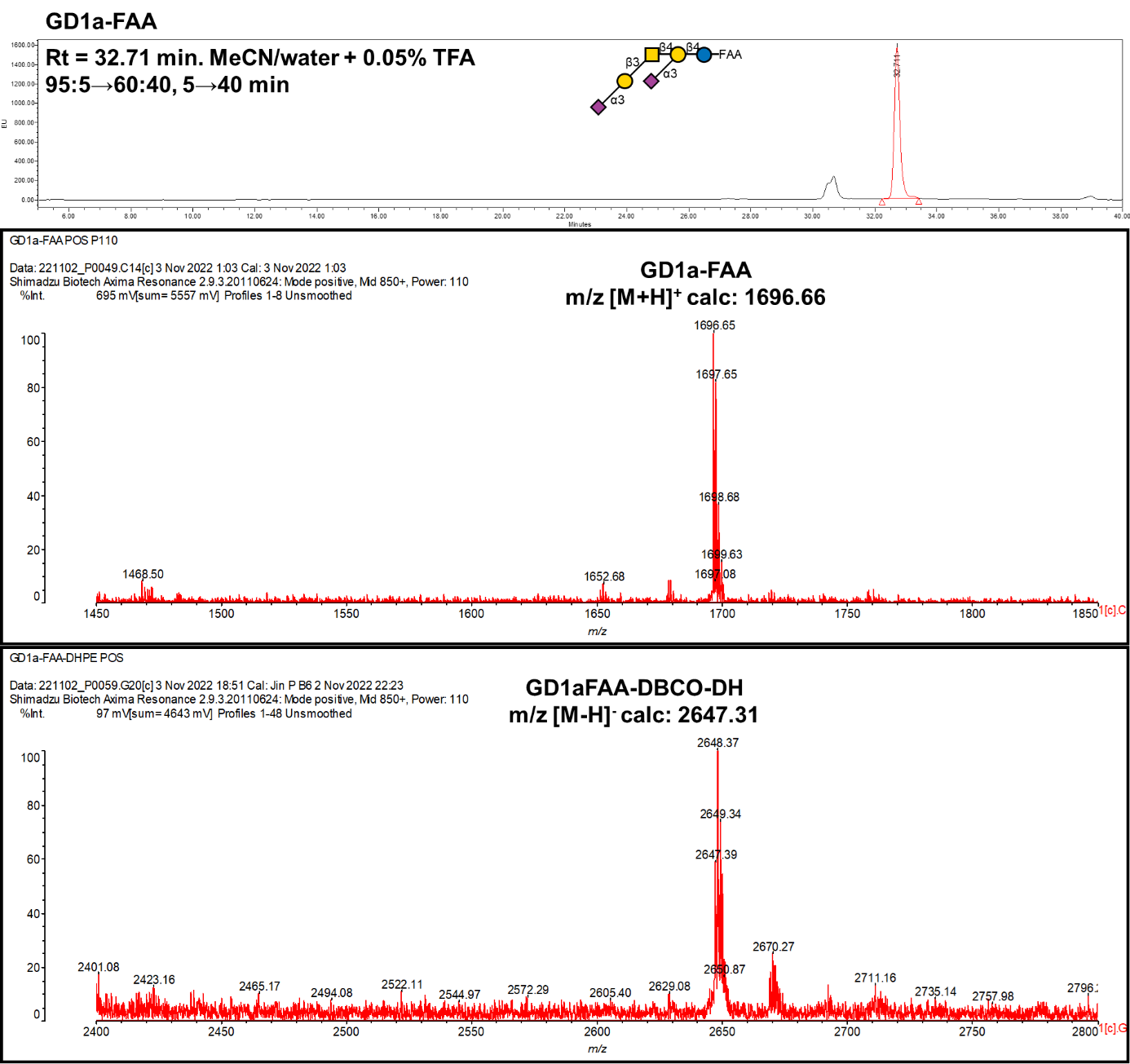

**Probe No 32**

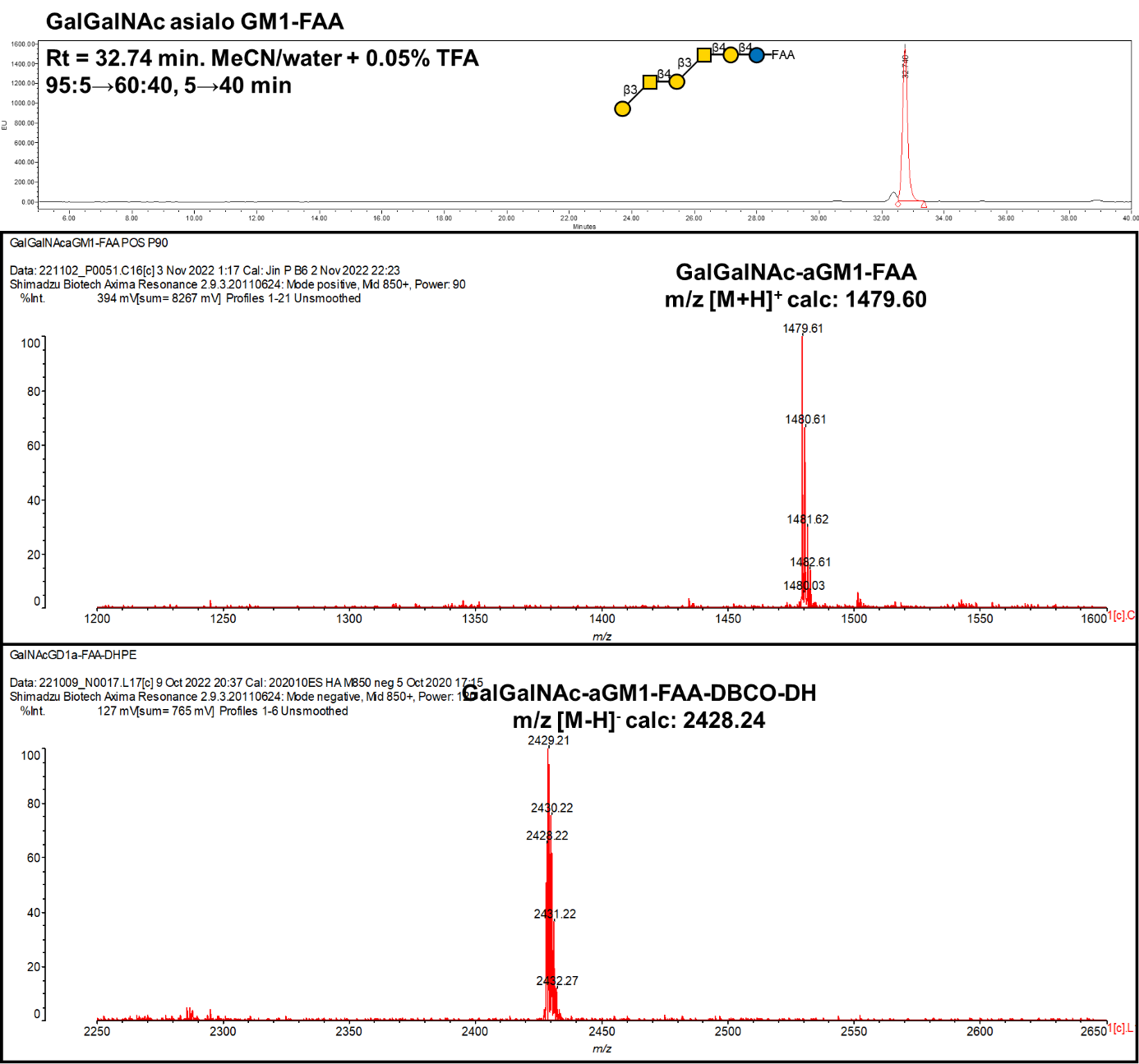

**Probe No 33**

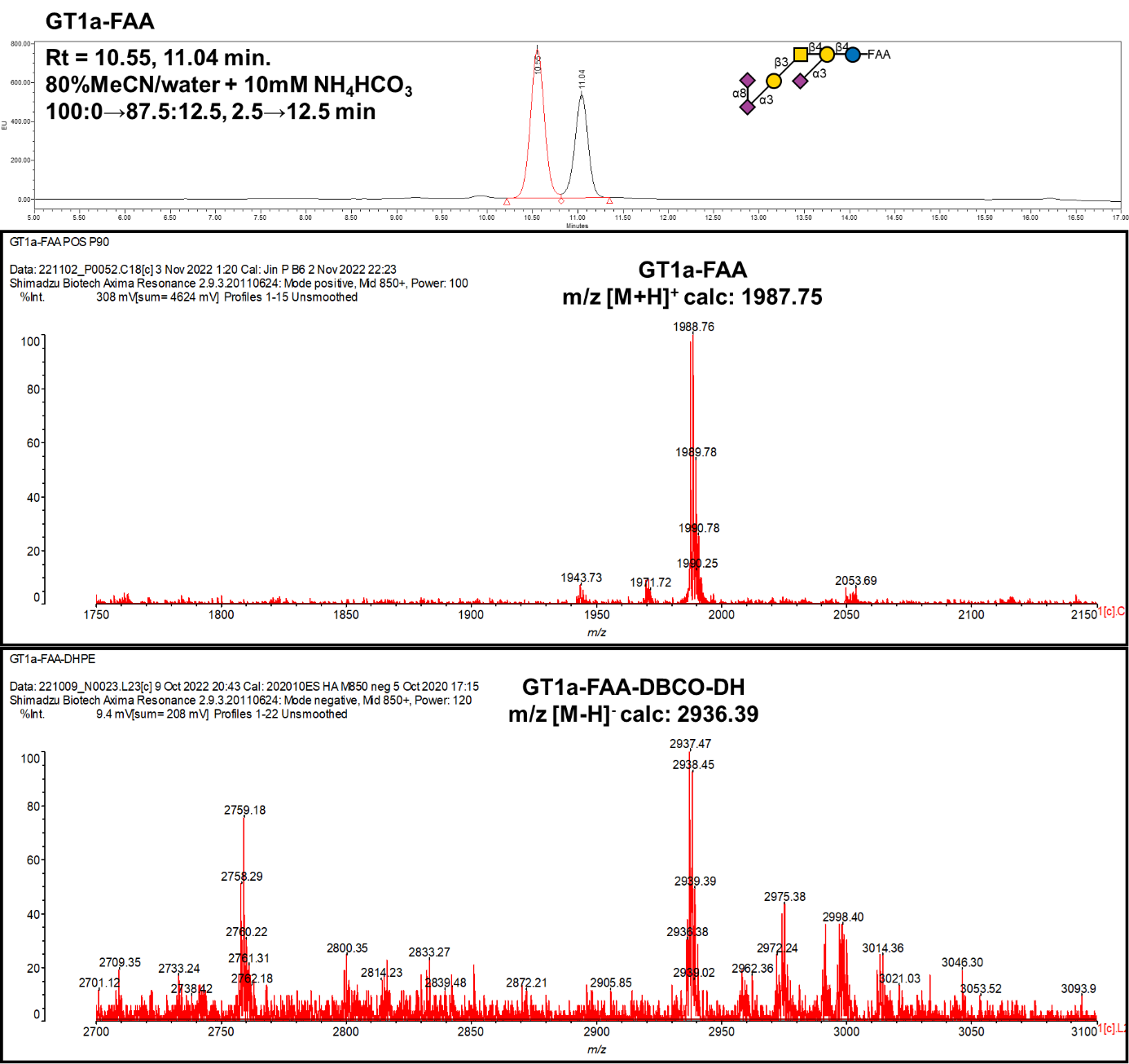

**Probe No 34**

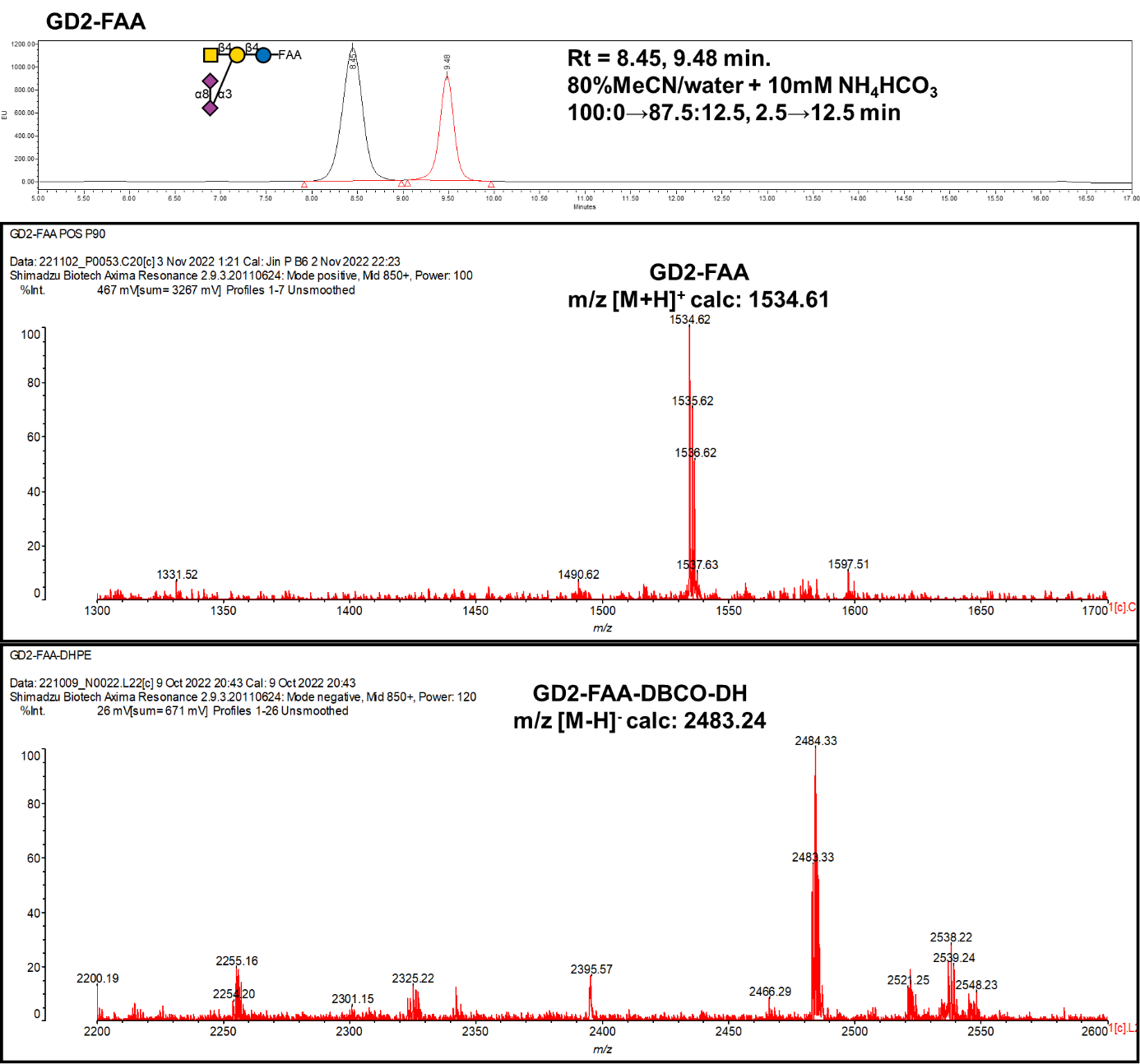

**Probe No 35**

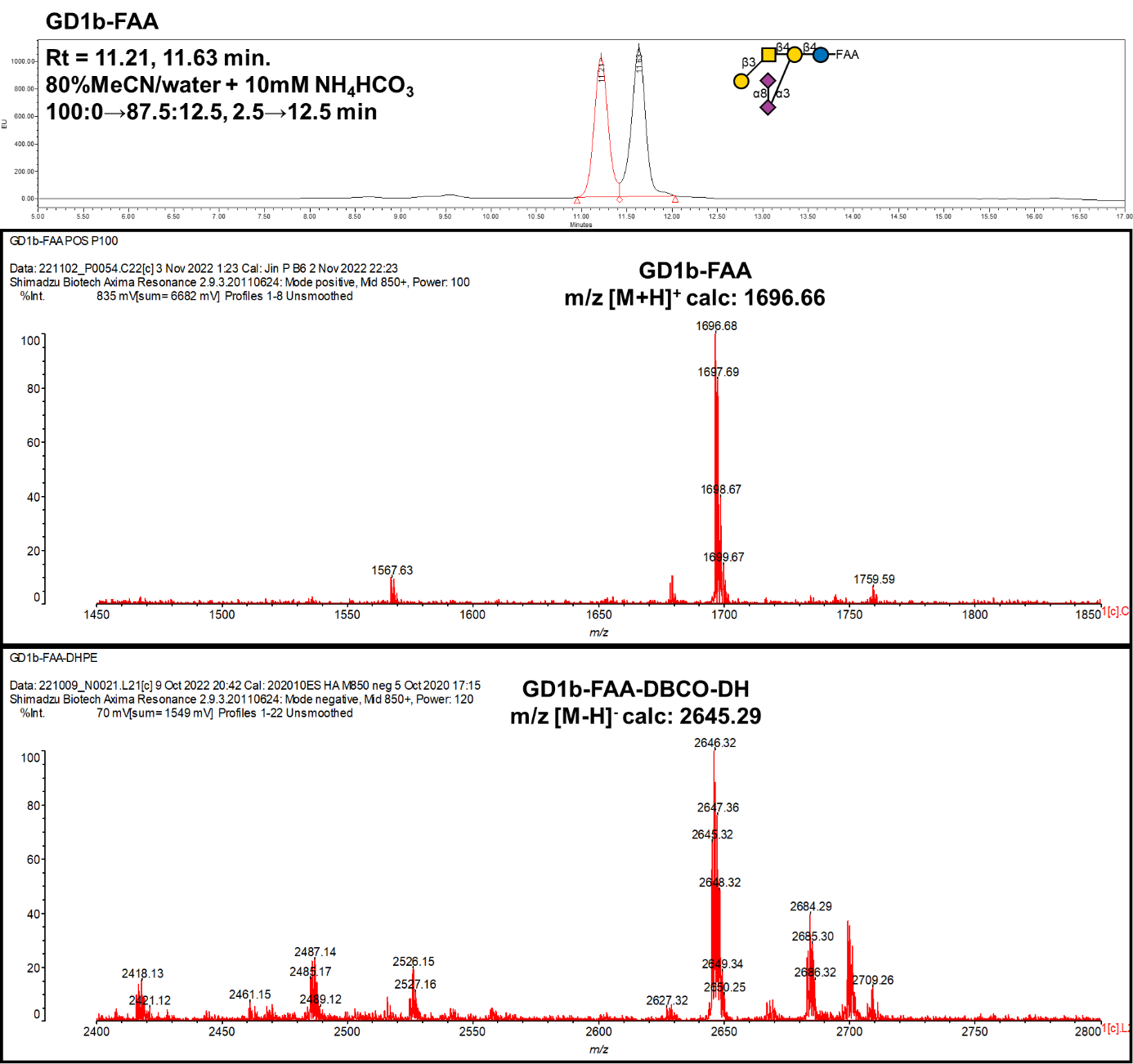

**Probe No 36**

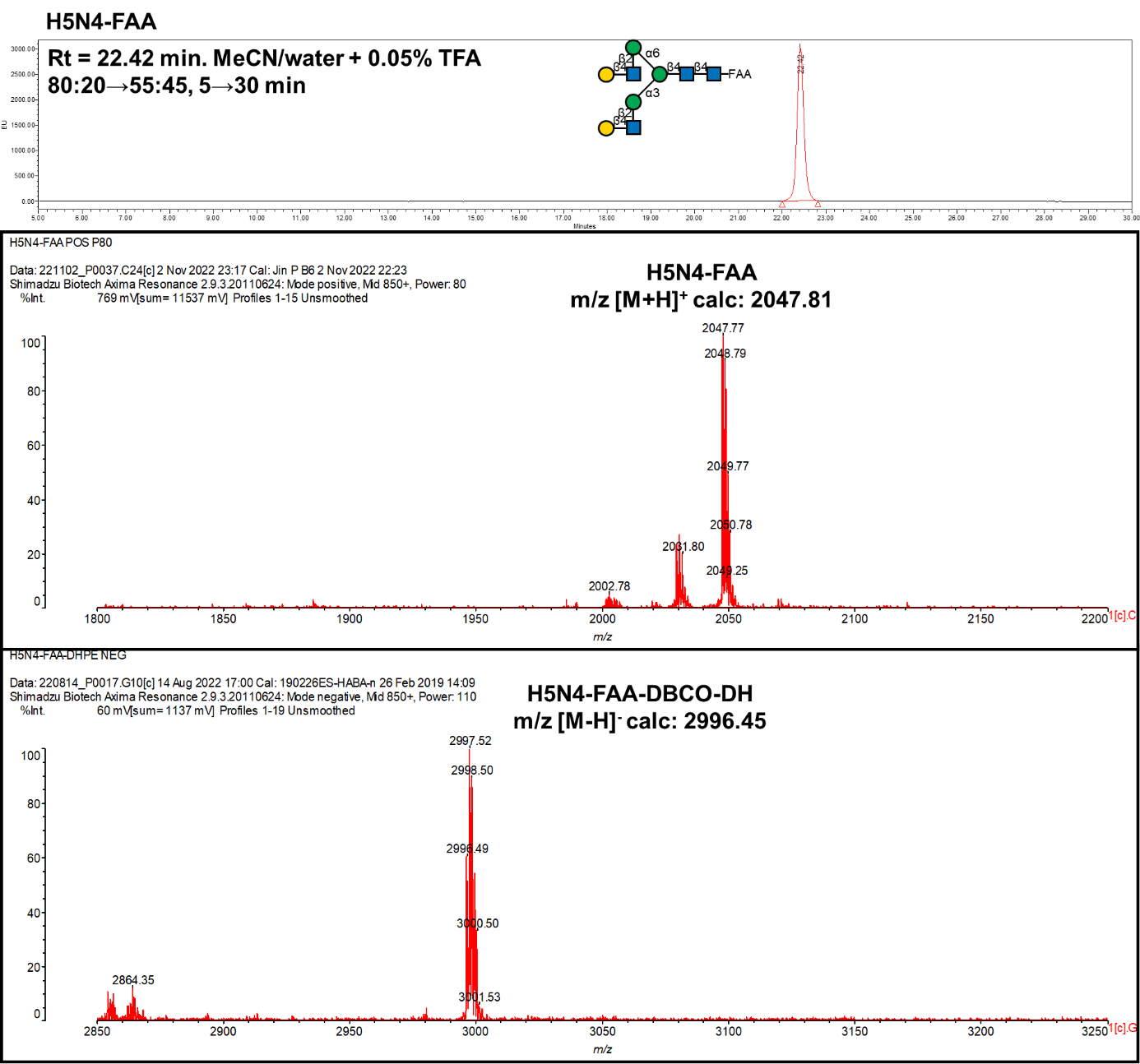

**Probe No 37**

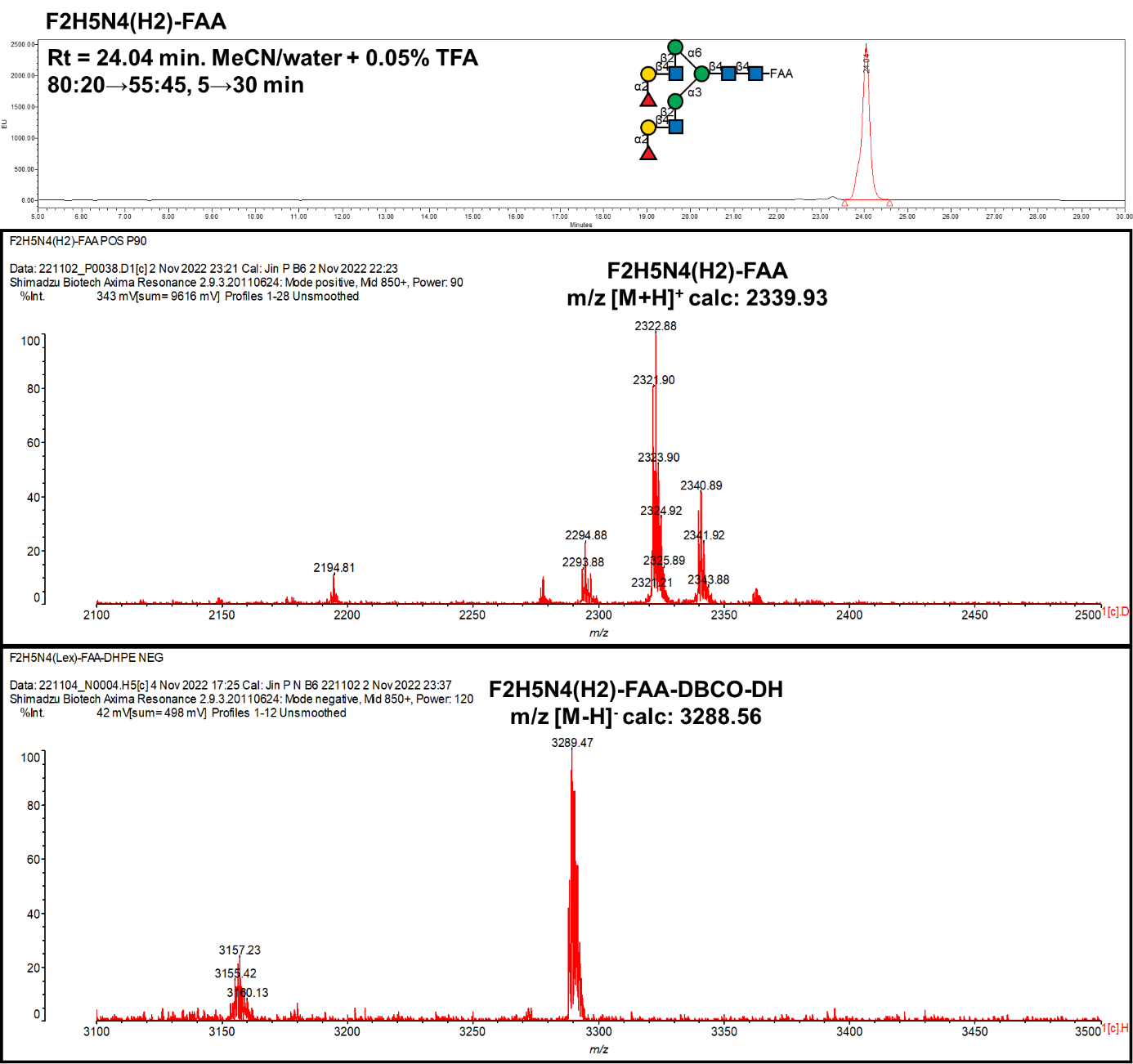

**Probe No 38**

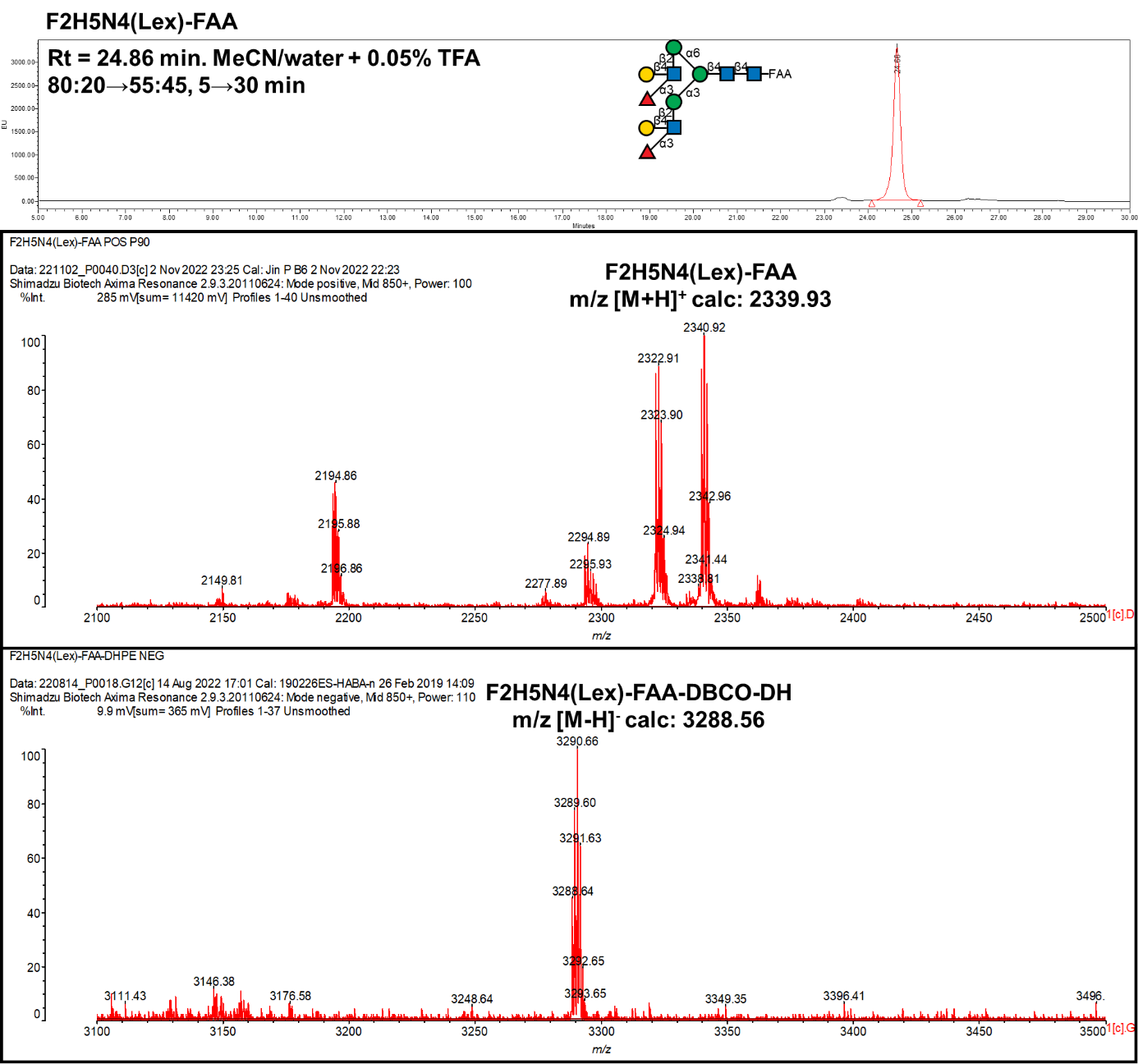

**Probe No 9**

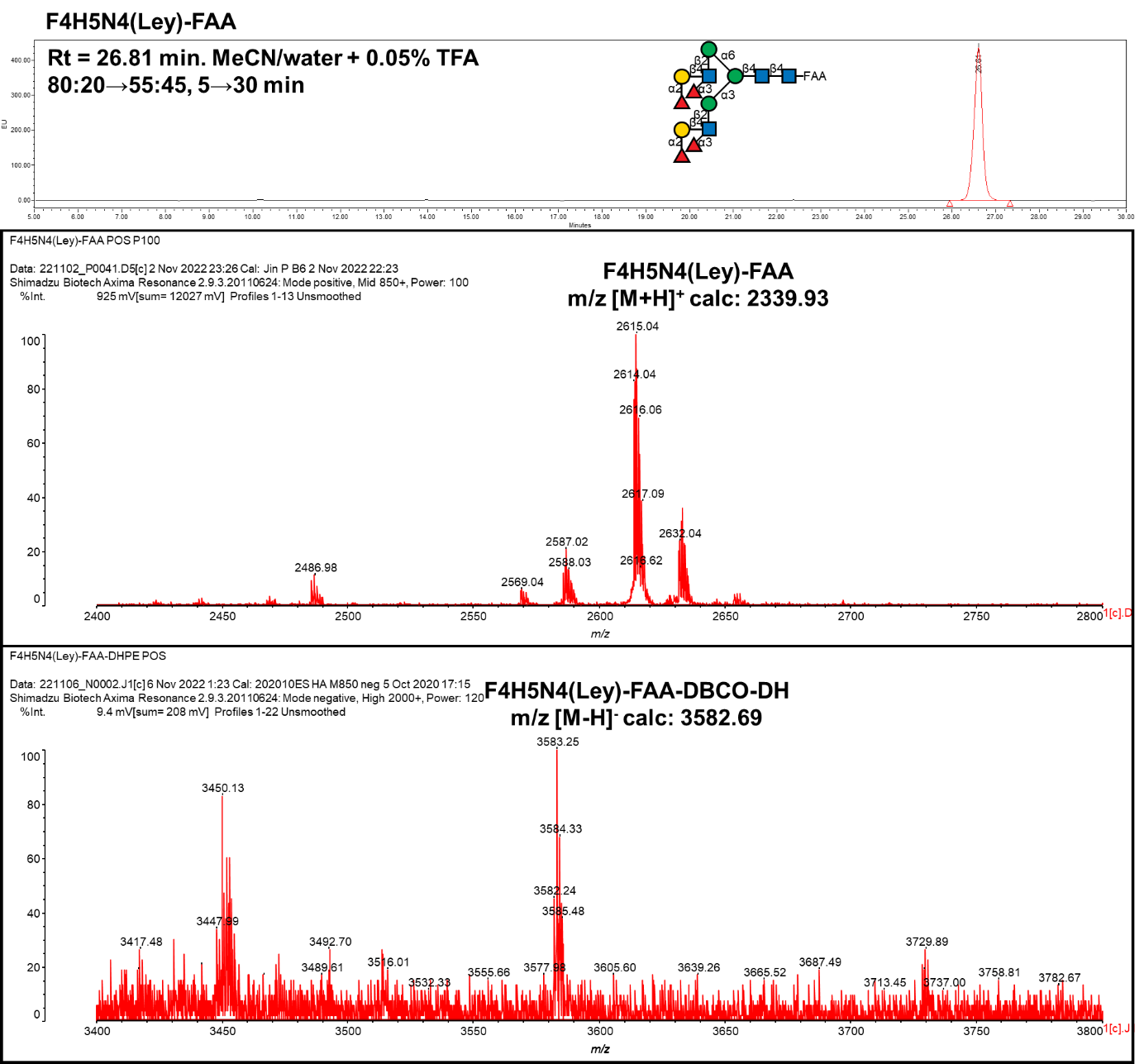

**Probe No 40**

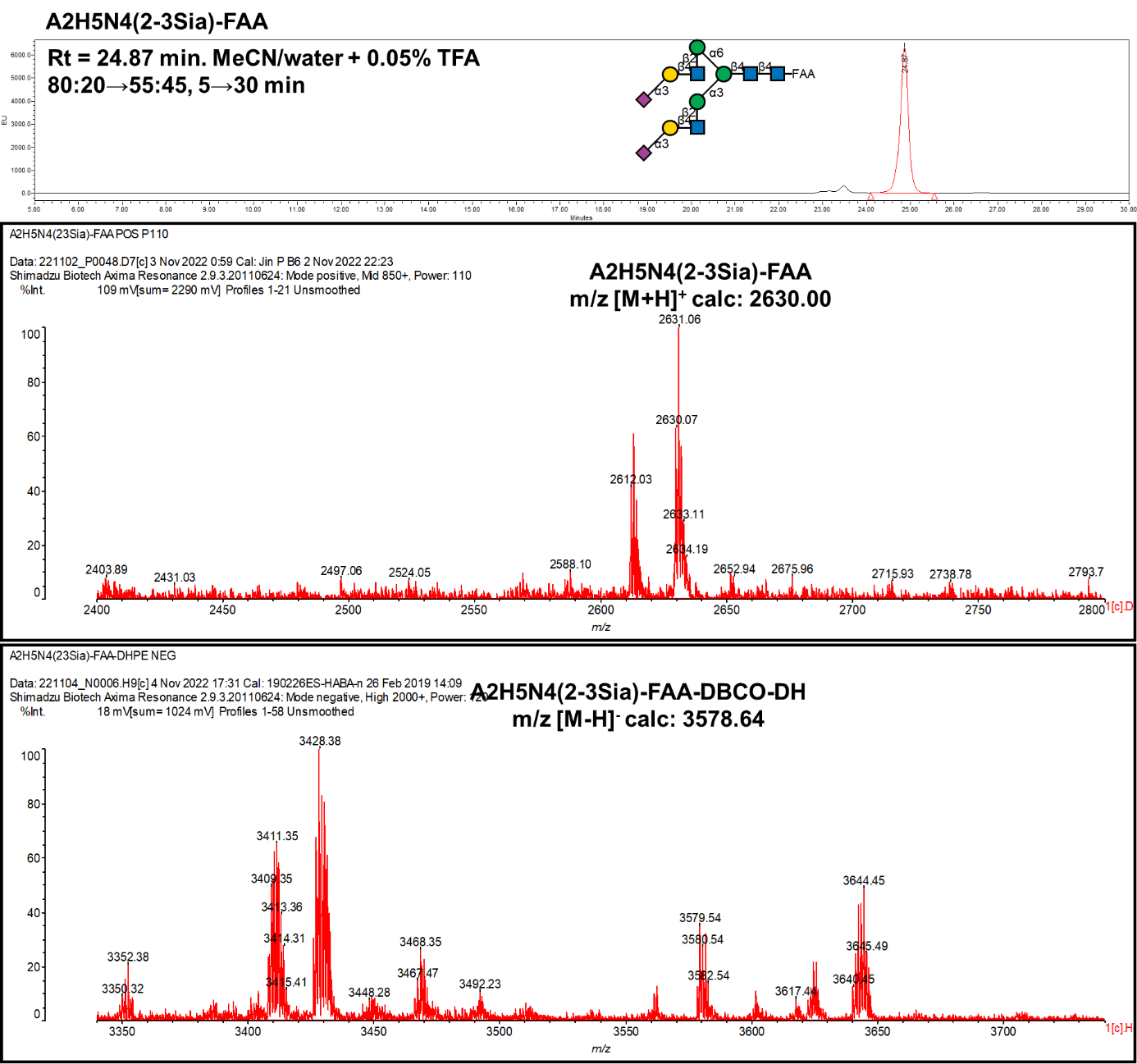

**Probe No 41**

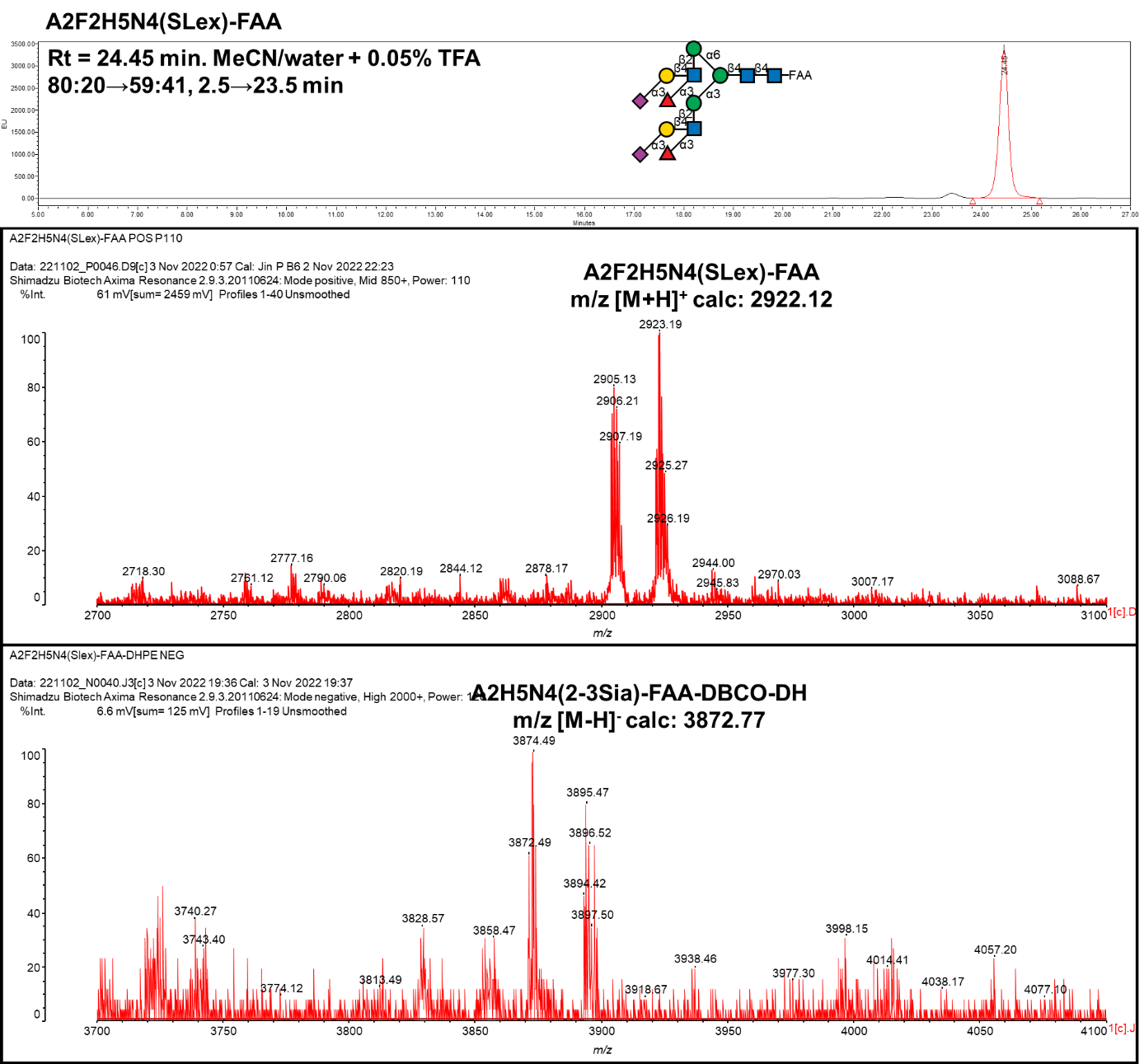

**Probe No 42**

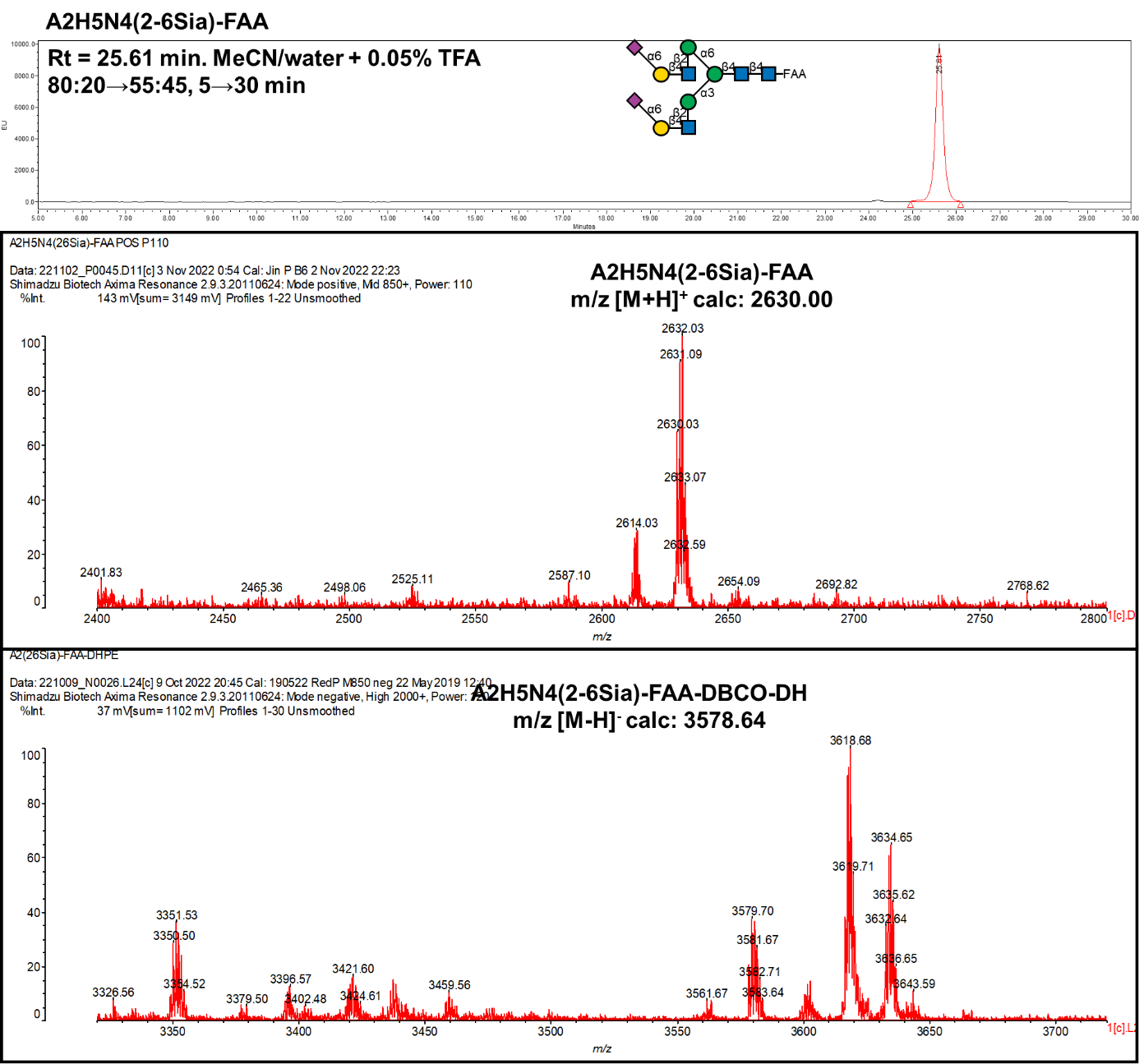

**Probe No 43**

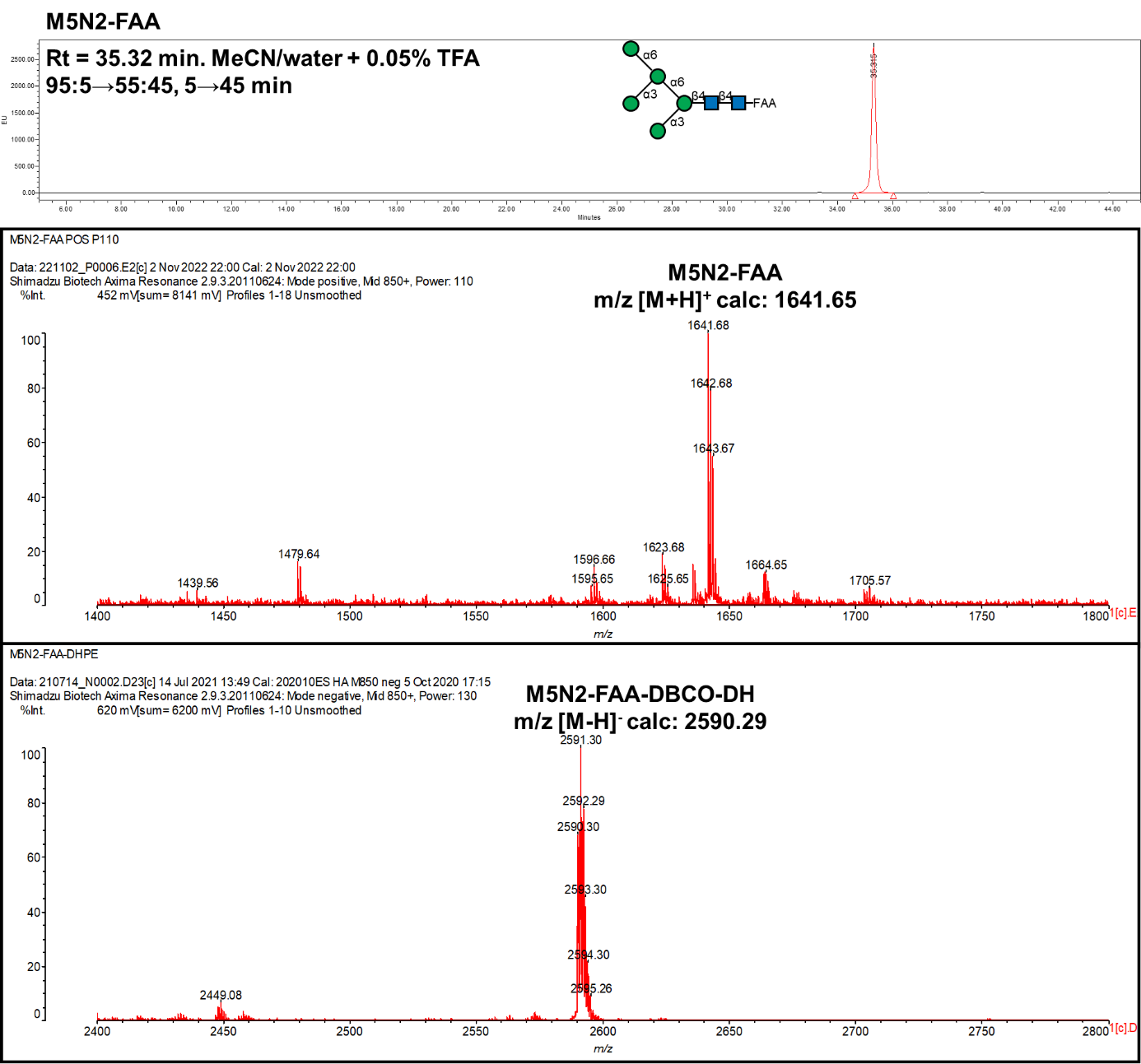

**Probe No 44**

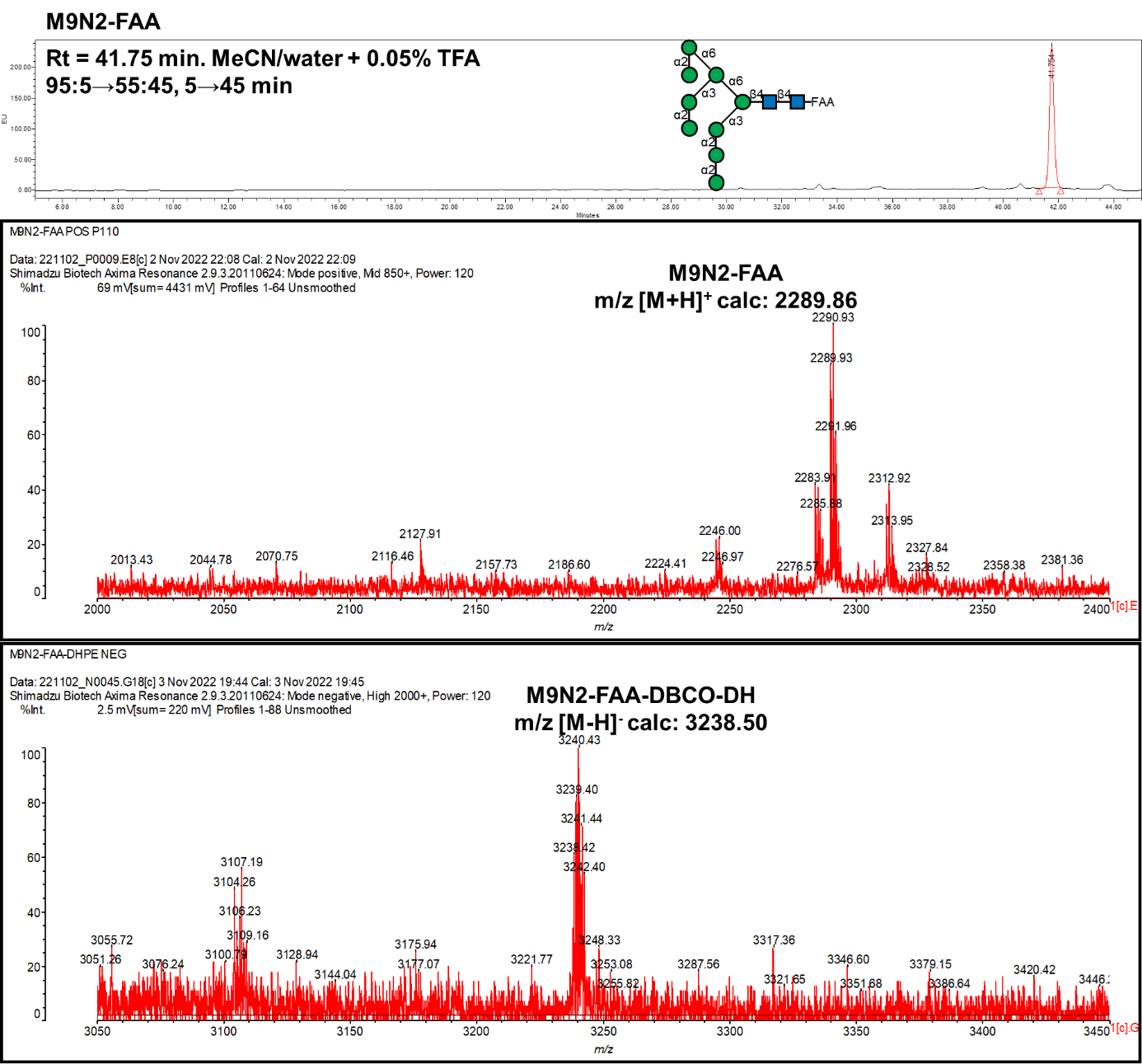

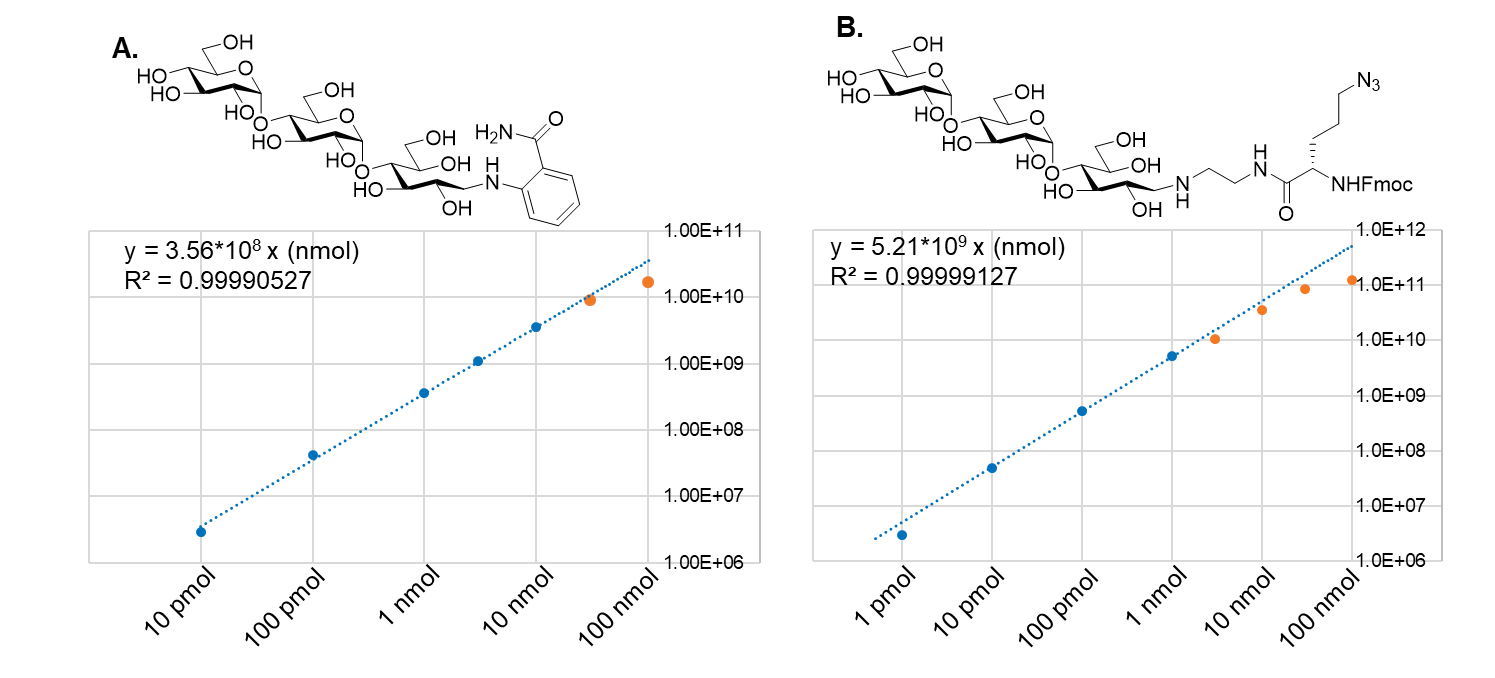

Figure S7. Generation of correlation ‘curve’.

Correlation curve of florescence intensity with amounts (nmol) of tagged glycans using maltotriose as a standard tagged with 2AB in **A** and with FAA in **B**. The blue data points within the linear range were used for quantitation rather than the red data points which were at saturation of florescence detection.

Figure S8. Examples of MALDI spectra of Fmoc deprotected glycan probes.

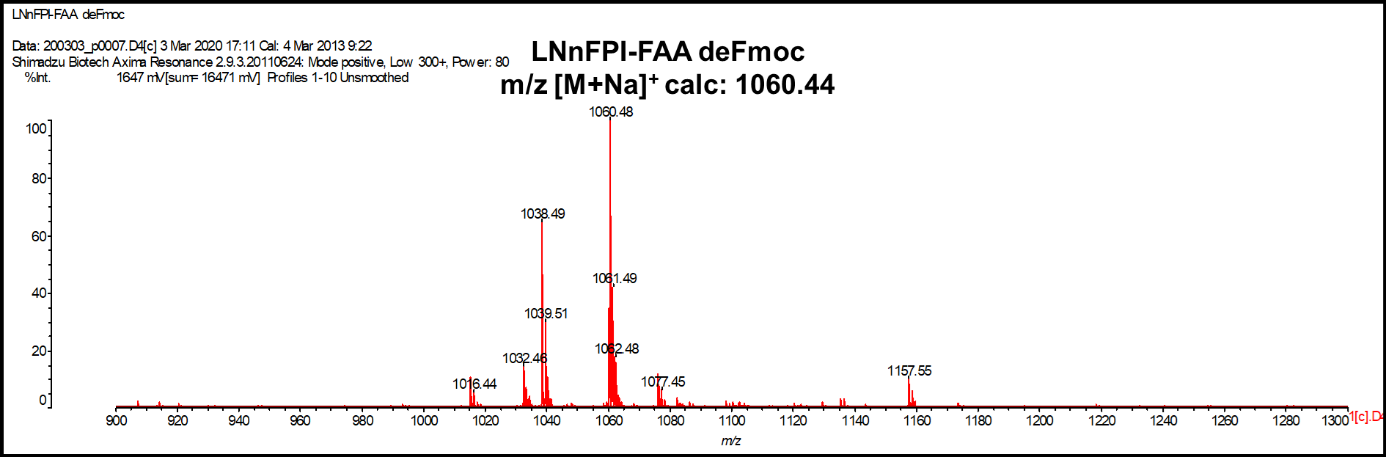

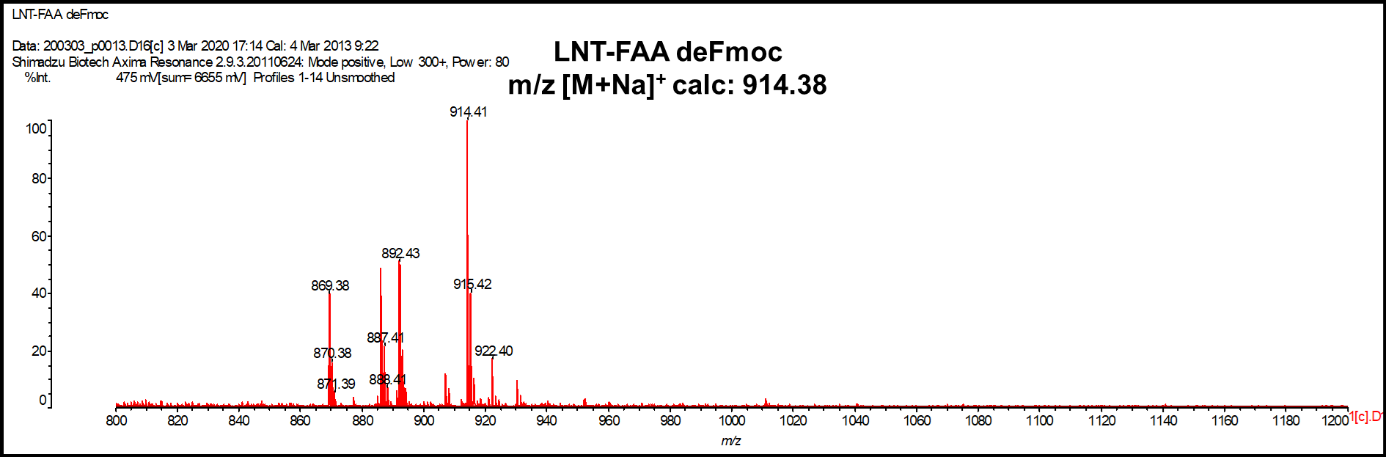

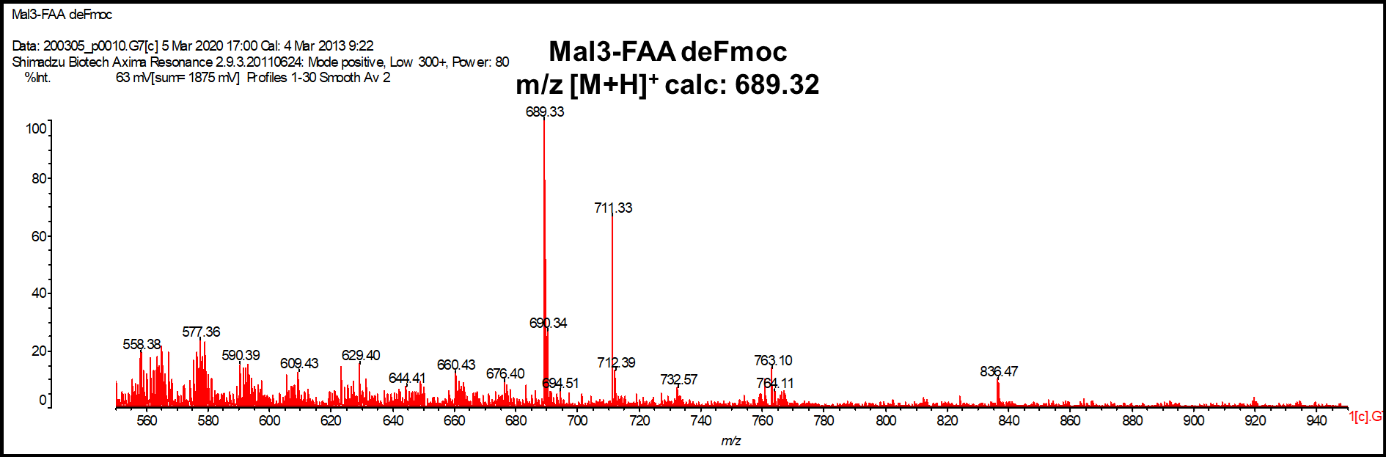

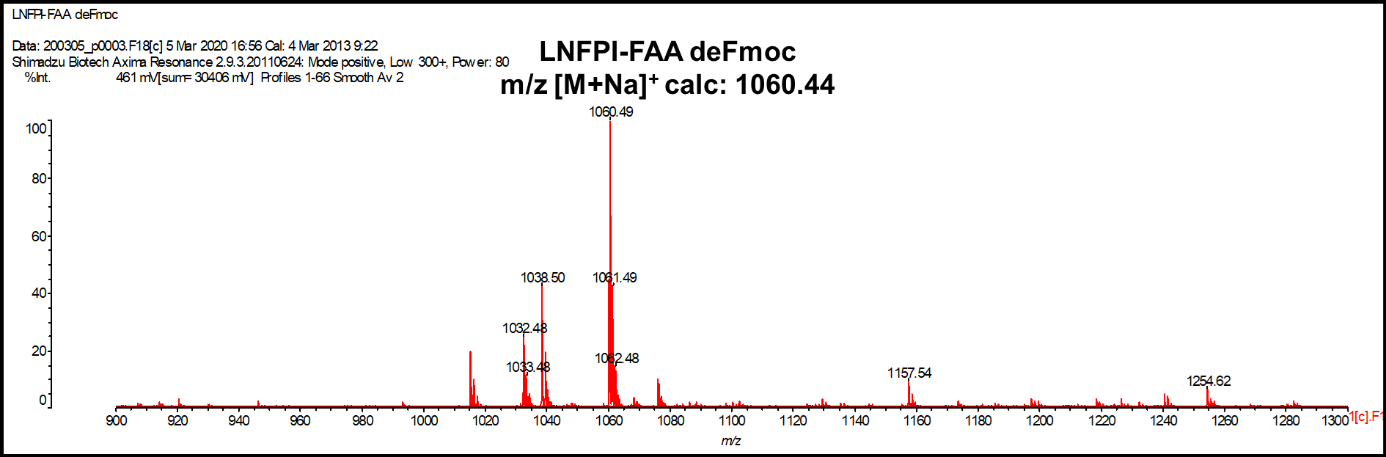

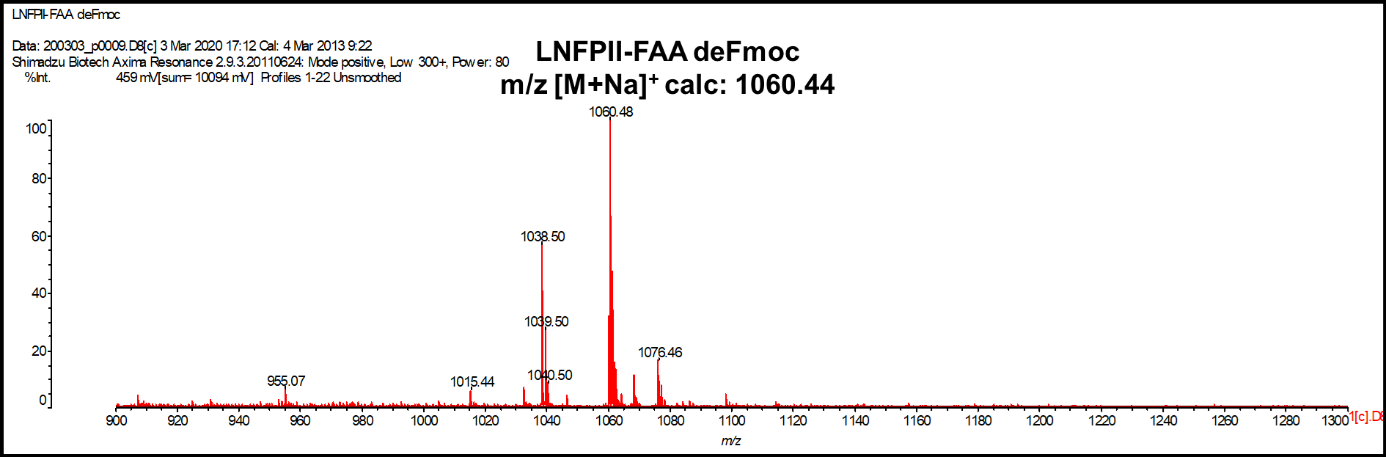

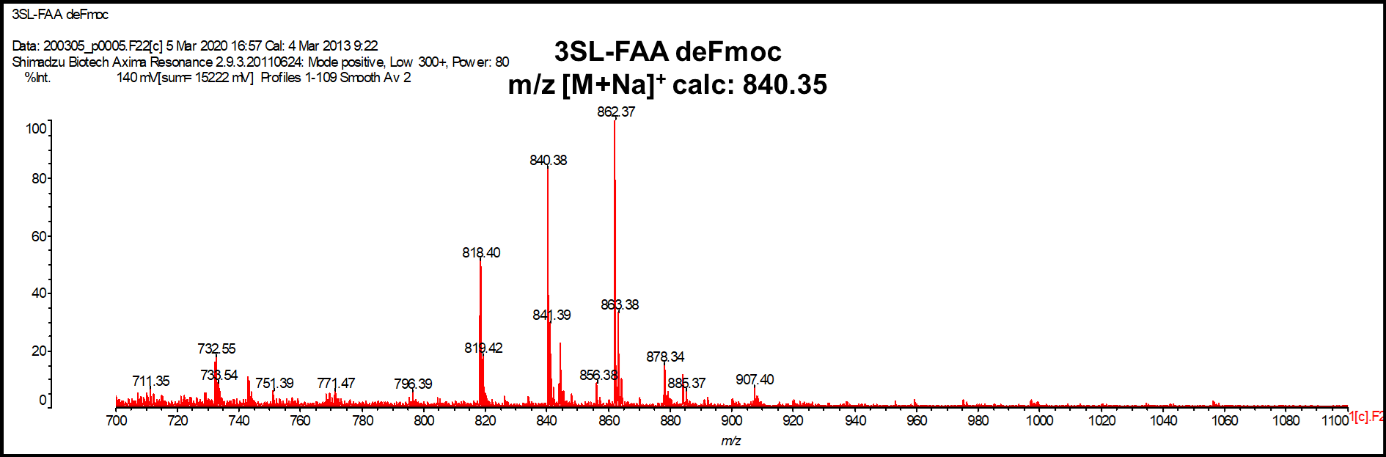

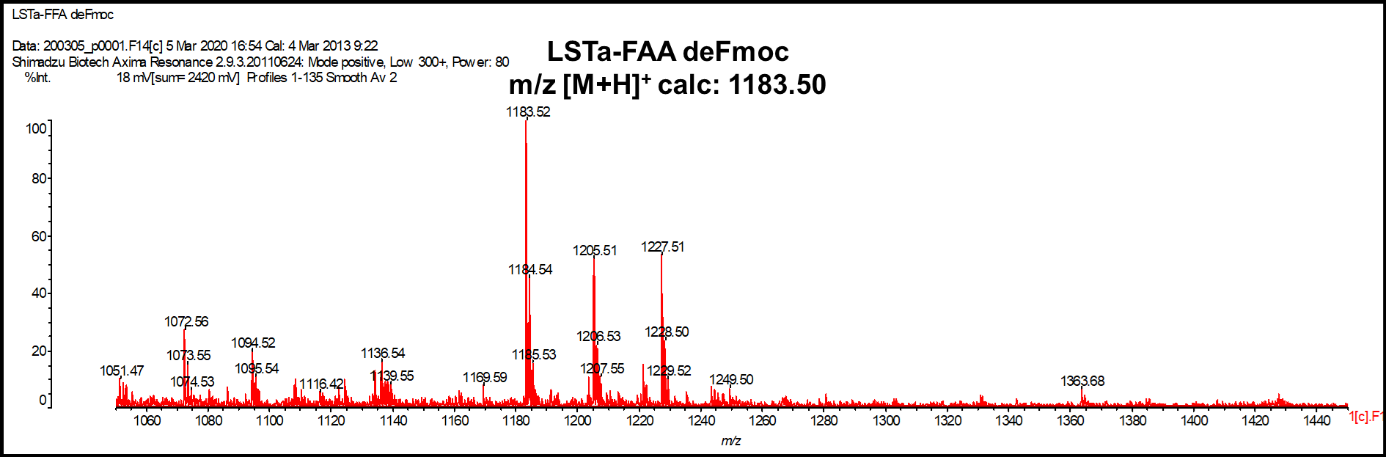

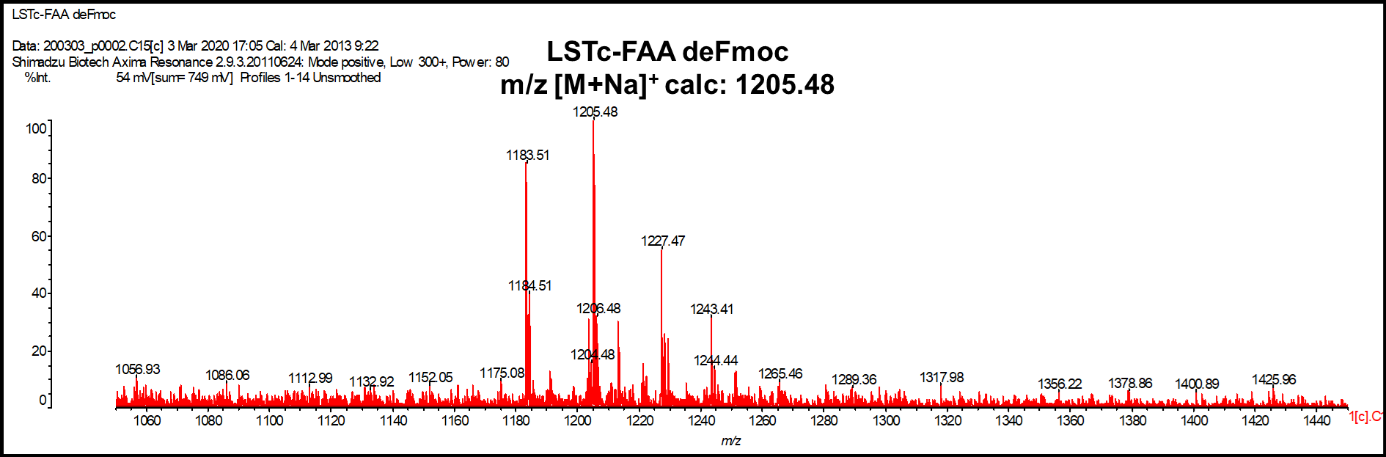

Figure S9. MALDI spectra of F_545_ labeled glycan probes.

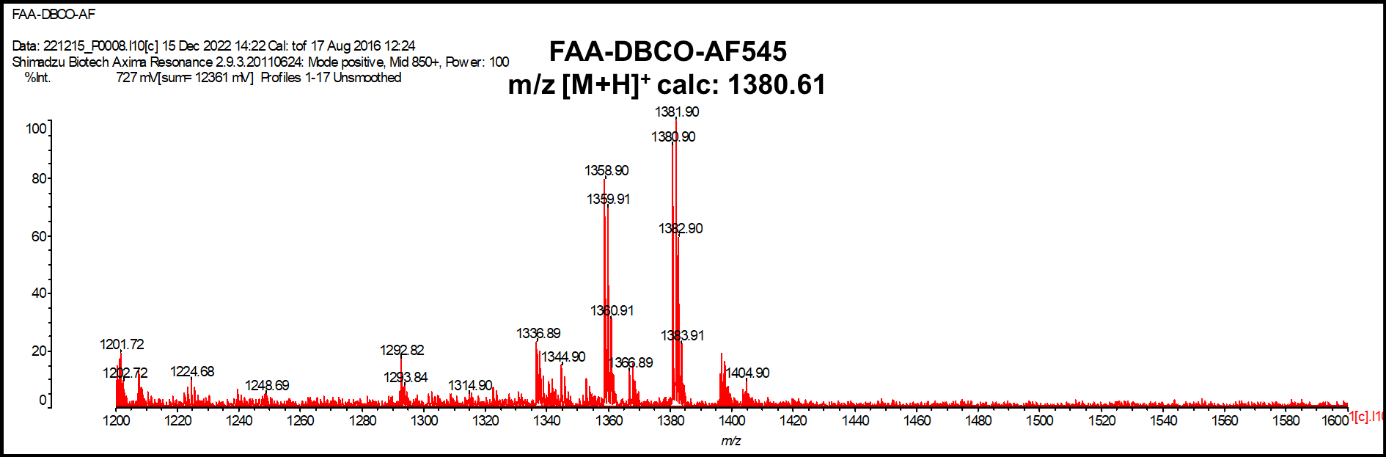

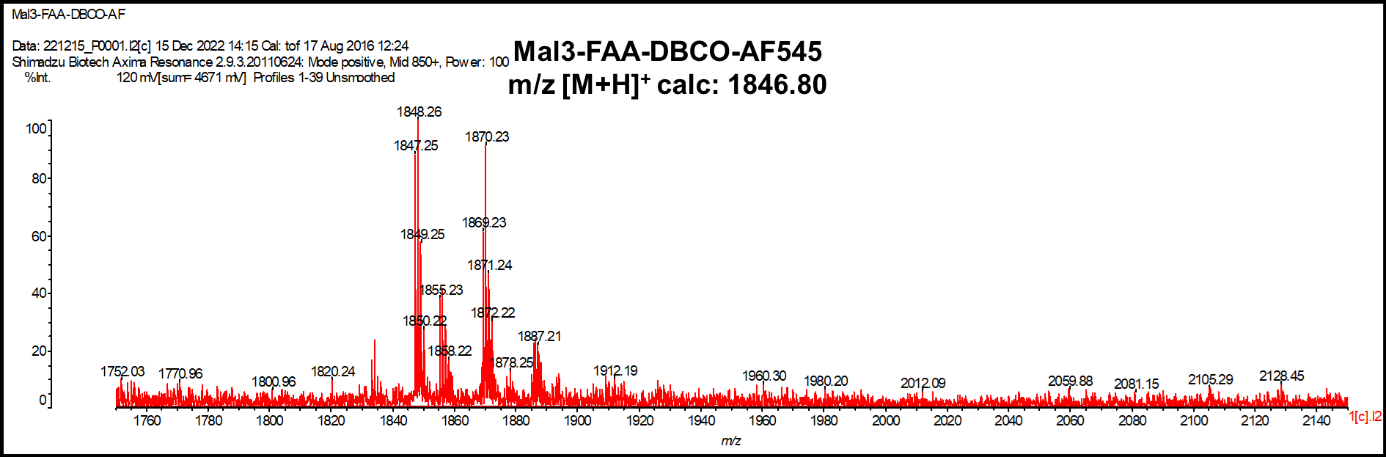

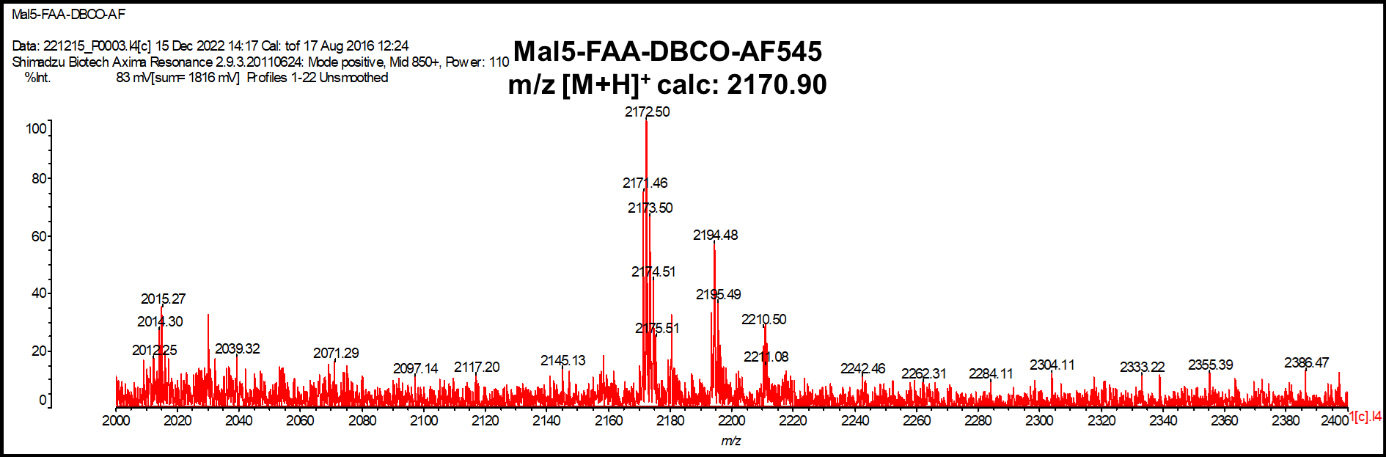

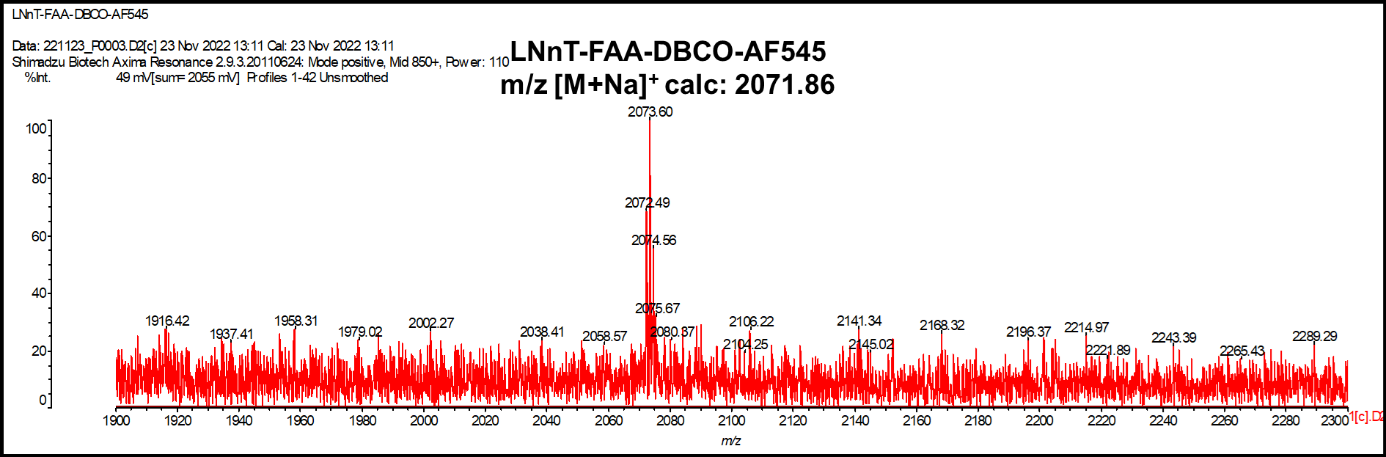

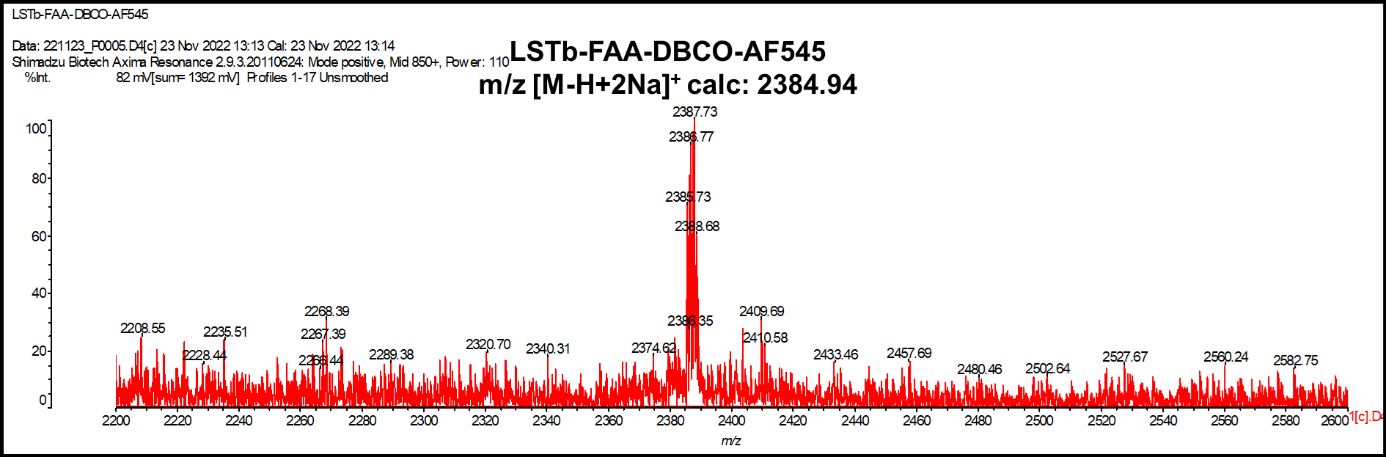

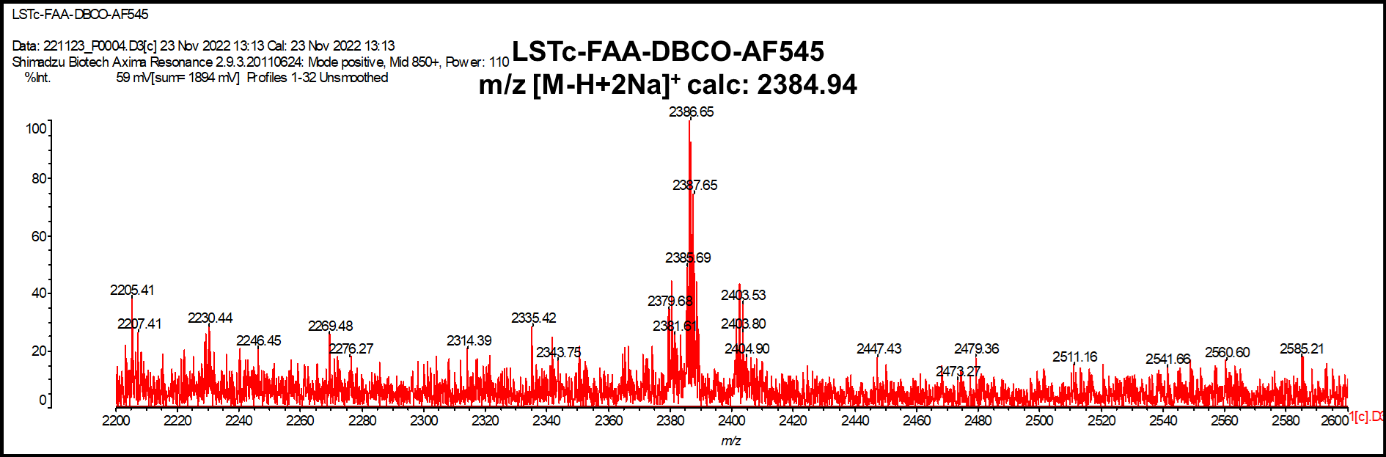

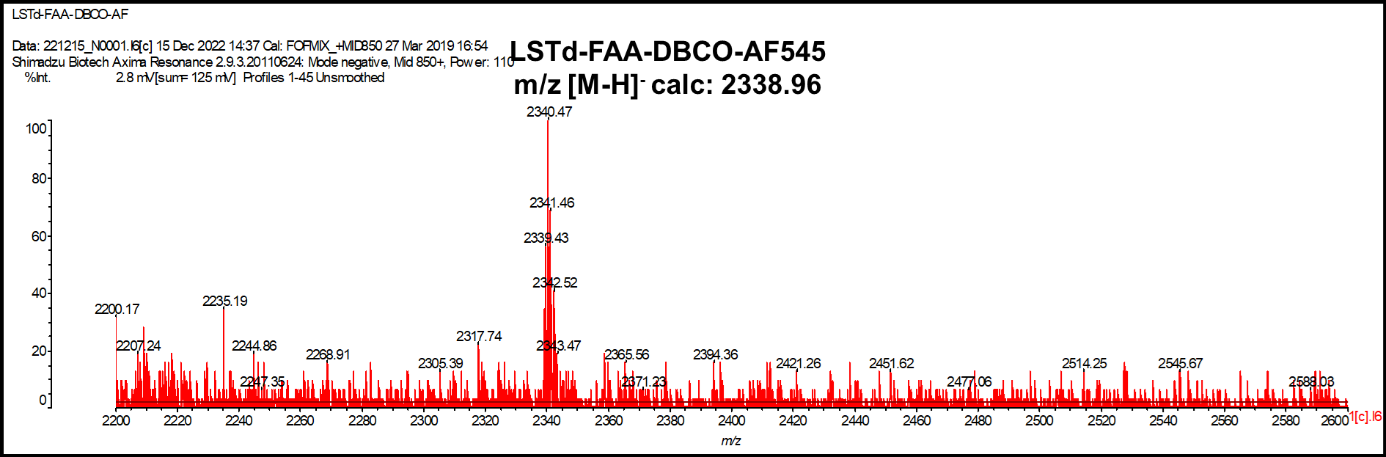

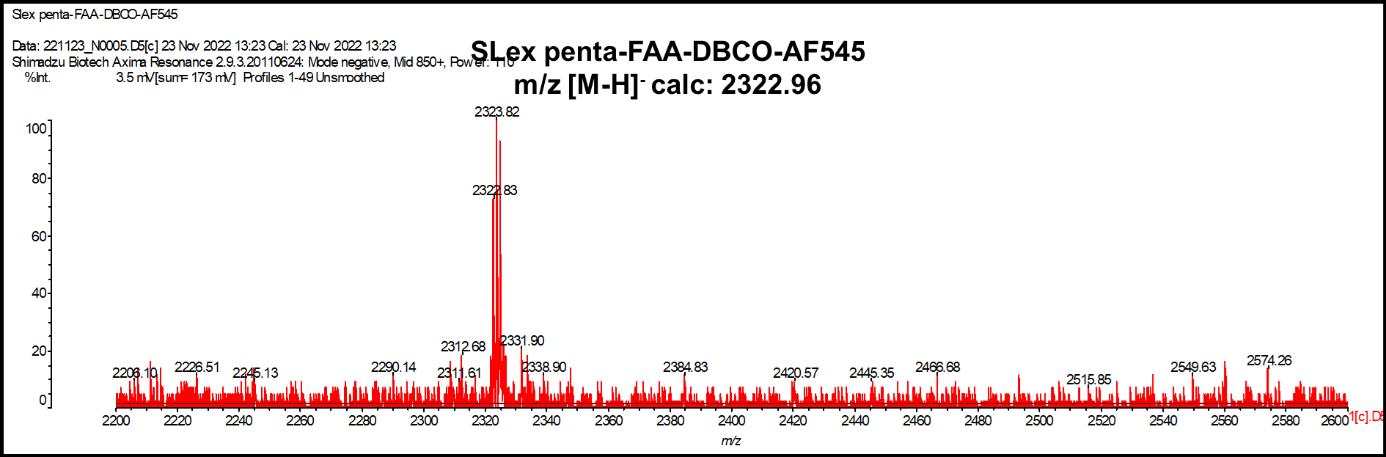

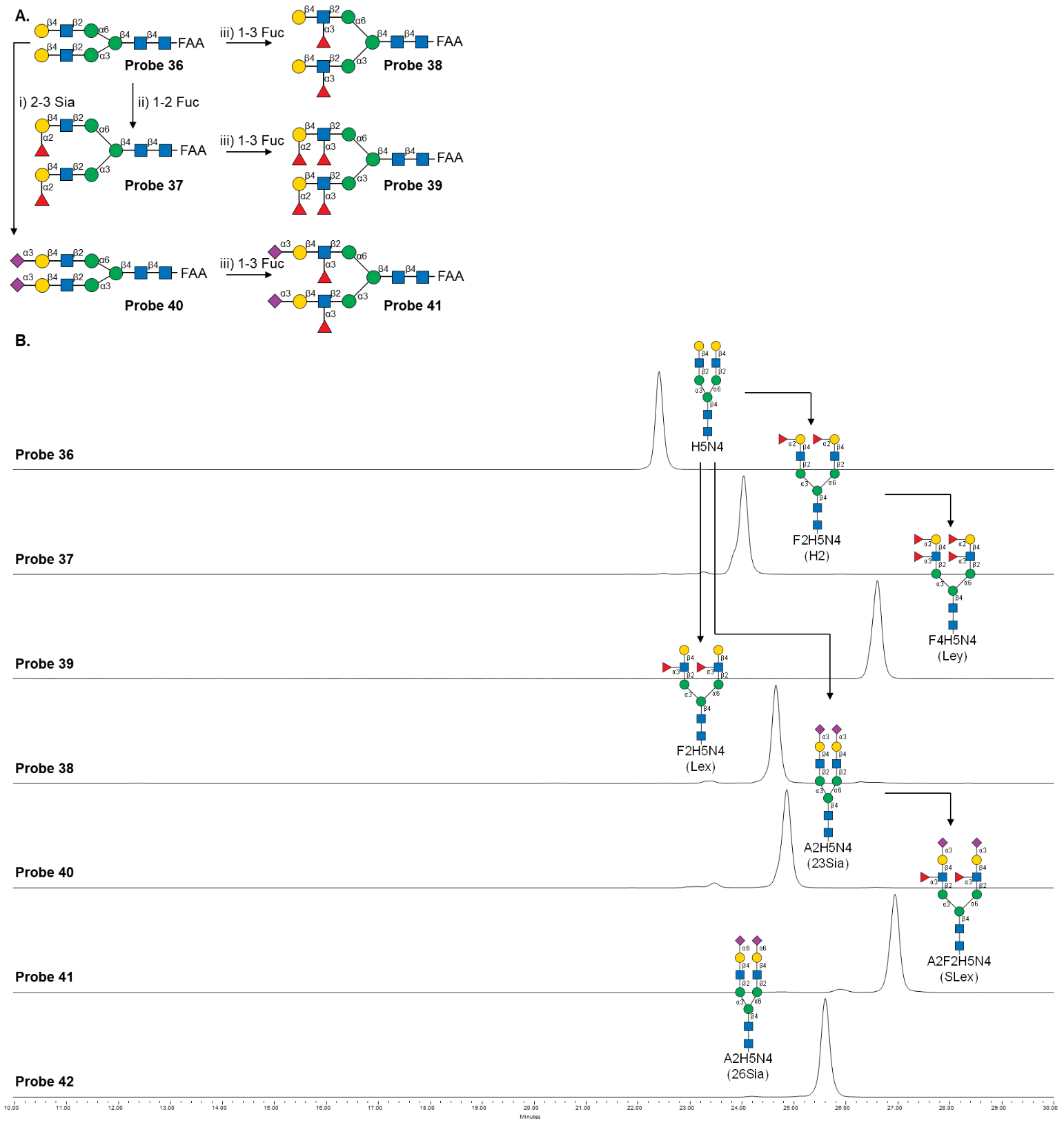

Figure S10. Enzymatic modifications of N glycan probe 36 to generate biantennary *N*-glycan probes 37-41. A) Synthetic schemes: i) Sialylation by 2α→3 sialyltransferase from *Pasteurella multocida*. ii) Fucosylation by 1α→2 fucosyltransferase from *Helicobacter hepaticus*. iii) Fucosylation by 1α→3/4 fucosyltransferase from *Helicobacter mustelae*. B)

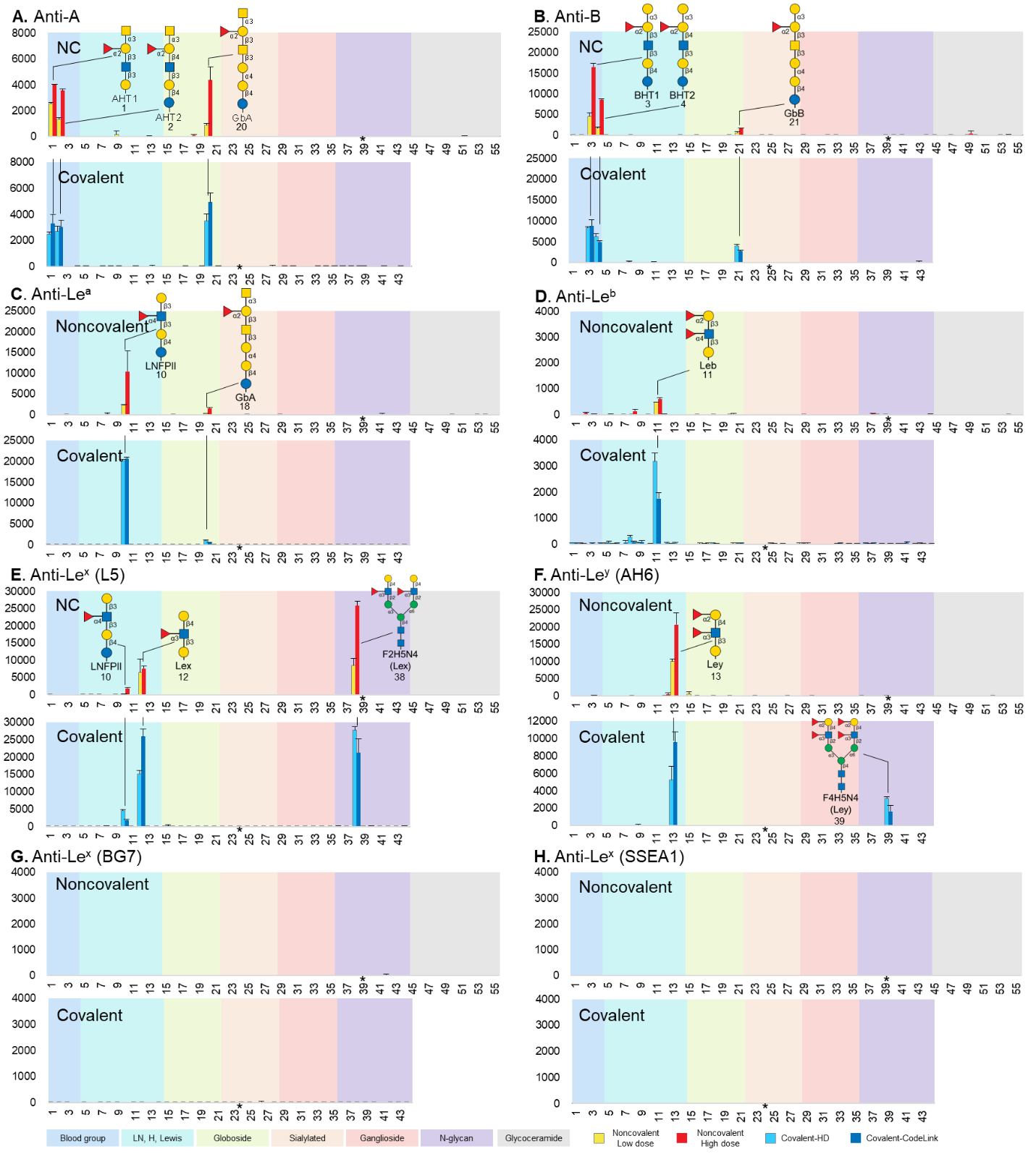

Figure S11. Binding analyses of sequence defined noncovalent and covalent arrays using antibodies.

* Probe 24 on the covalent arrays and probe 39 on the NGL array were misprinted and did not pass quality control.

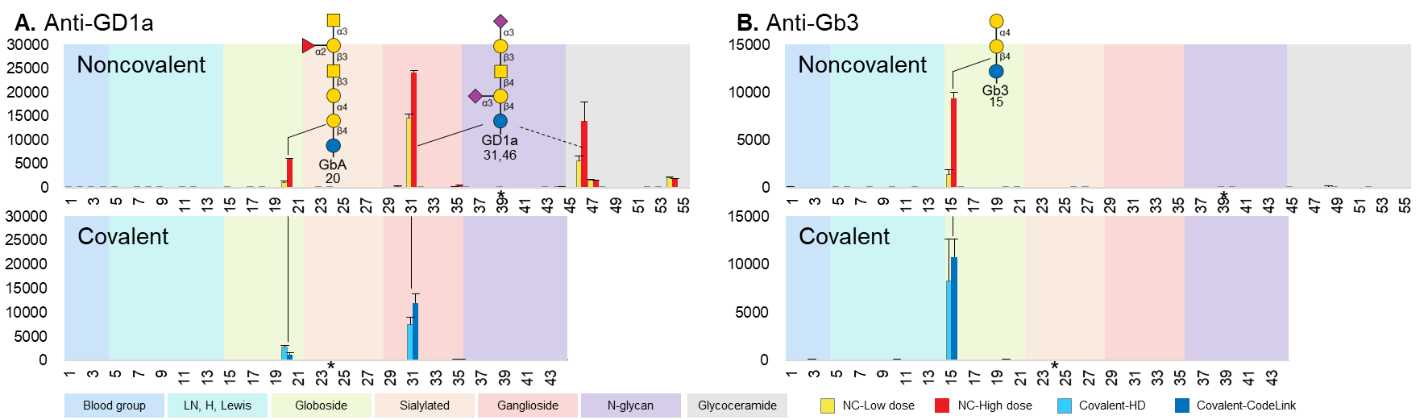

Figure S12. Binding analyses of sequence defined noncovalent and covalent arrays using anti-ganglioside antibodies.

* Probe 24 on the covalent arrays and probe 39 on the NGL array were misprinted and did not pass quality control.

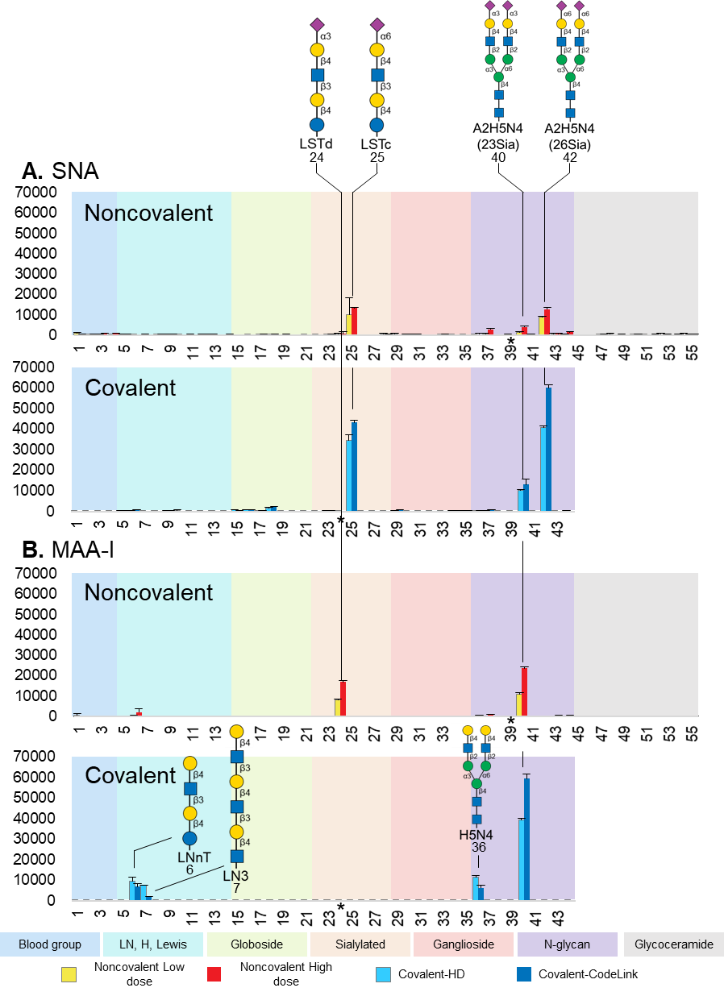

Figure S13. Binding analyses of sequence defined noncovalent and covalent arrays using Sialic acid binding lectins.

* Probe 24 on the covalent arrays and probe 39 on the NGL array were misprinted and did not pass quality control.

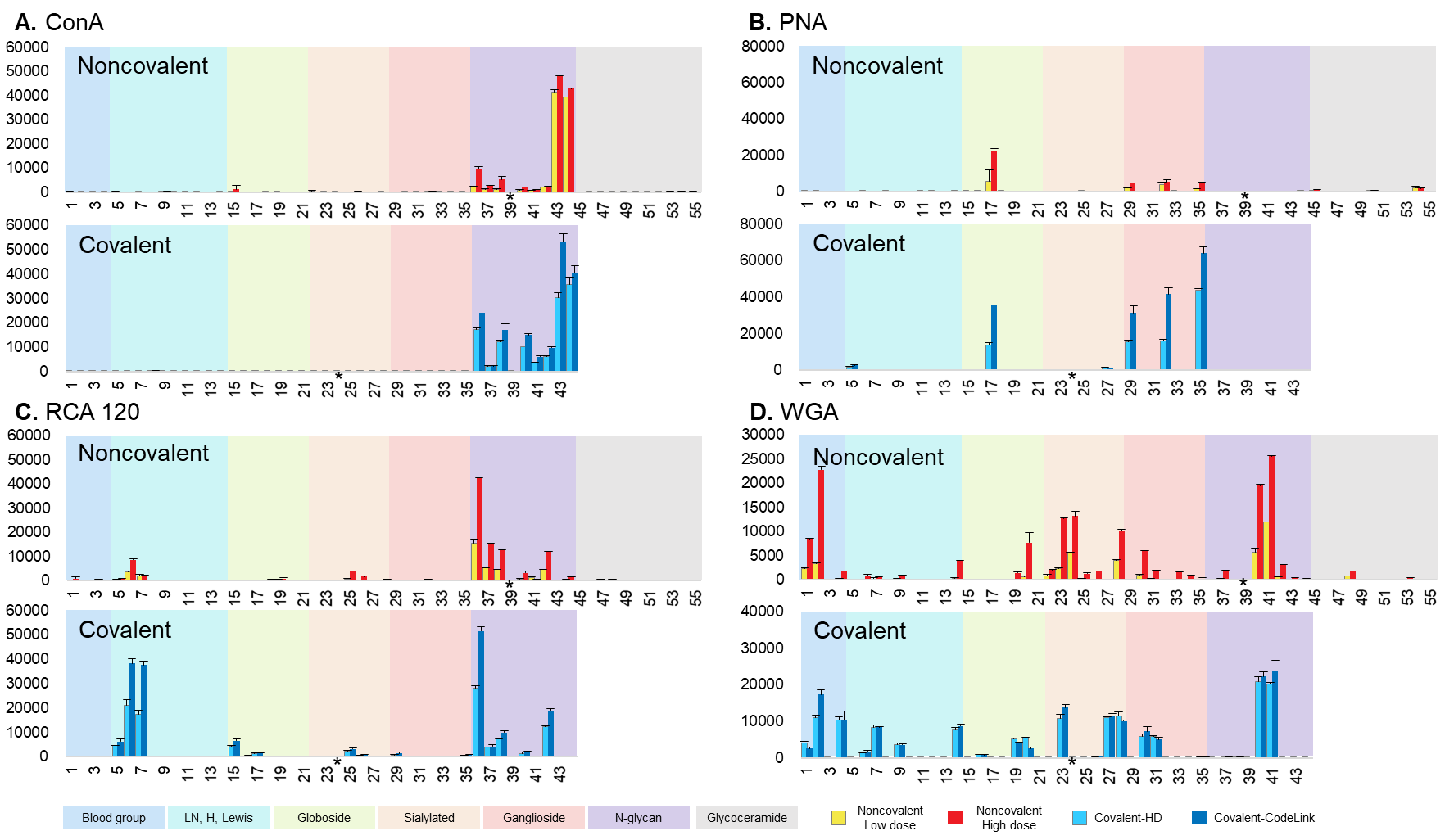

Figure S14. Binding analyses of sequence defined noncovalent and covalent arrays using plant lectins.

* Probe 24 on the covalent arrays and probe 39 on the NGL array were misprinted and did not pass quality control.

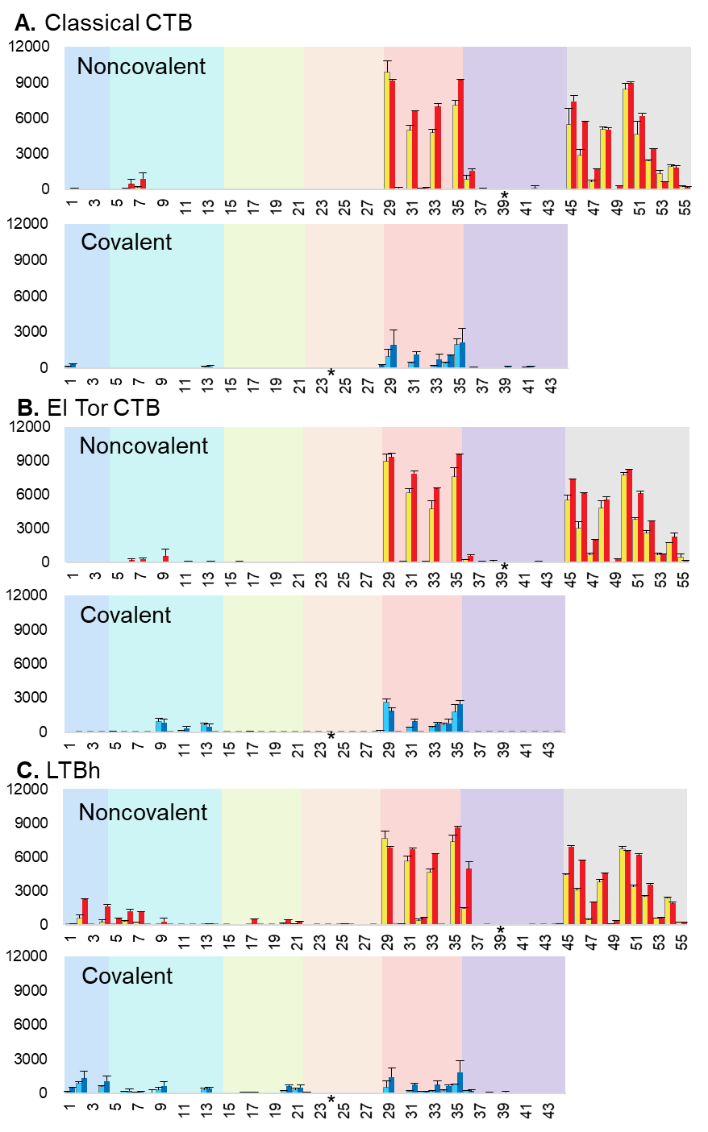

Figure S15. Binding analyses of sequence defined noncovalent and covalent arrays using bacteria toxins at a lower level.

Protein concentration of 0.5 μg/mL was used, in comparison with incubations using 5 μg/mL shown in **Fig 5D-F** in the main text.

* Probe 24 on the covalent arrays and probe 39 on the NGL array were misprinted and did not pass quality control.

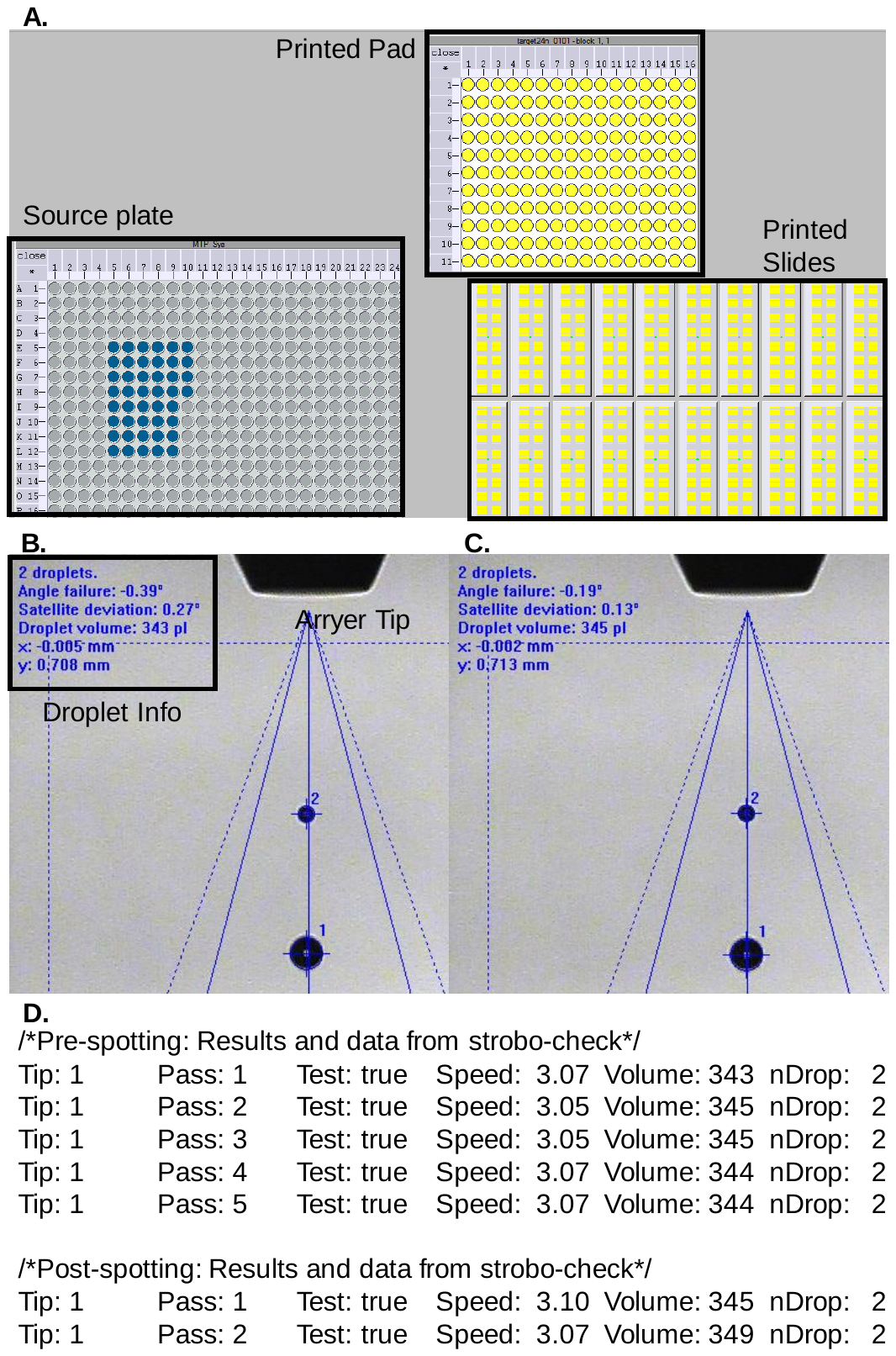

Figure S16. Spotting record of covalent 44 probe FAA glycan array.

A. Arrayer software display monitoring the printing status. The yellow circles indicating successfully printed spots. B. Stroboscope image of the pre-spotting check of the droplets before the printing of probe 24. C. Stroboscope image of the post-spotting check of the droplets after the printing of probe 24. D. Log documents the printing process, including 5 pre-spotting checks and 2 post-spotting checks. During these checks, all droplet volumes remained within ±10% of the set value of 340 pL, and the droplet speed stayed within ±10% of the set value of 3.0 m/s. These results indicate that the printing process passed the self-check program.

MIRAGE compliant Glycan Microarray Document

Supplementary Glycan Microarray Document Based on MIRAGE Guidelines (doi:10.3762/mirage.3)

|  | Description |
| --- | --- |
| 1. Sample: Glycan Binding Sample | |
| Description of Sample | The virus VP1 proteins, siglects, CTBs, antibodies and lectins used in this study are listed in Table S2-6 |
| Sample modifications | Not relevant. |
| Assay protocol | The incubation protocol was included in SI section Microarray analysis of glycan binding. |
| 2. Glycan Library | |
| Glycan description for defined glycans | The sequence information on the 44 covalent probes and 44 NGL probes included in the microarray are in SI section Preparation of sequence defined glycans tagged with FAA |
| Glycan description for undefined glycans | Not relevant. |
| Glycan modifications | The synthesis of covalent and NGL probes were described in SI section Preparation of sequence defined glycans tagged with FAA |
| 3. Printing Surface; e.g., Microarray Slide | |
| Description of surface | Both covalent and noncovalent arrays were prepared in this study. The details are included in SI section Generation of glycan microarrays |
| Manufacturer | The details are included in SI section Generation of glycan microarrays |
| Custom preparation of surface | Not relevant. |
| Noncovalent Immobilization | The details are included in SI section Generation of glycan microarrays |
| 4. Arrayer (Printer) | |
| Description of Arrayer | Nano-Plotter 2.1 (GeSim, Radeberg, Germany). The details are included in SI section Generation of glycan microarrays |
| Dispensing mechanism | Non-contact liquid delivery with four dispensing tips. |
| Glycan deposition | The details are included in SI section Generation of glycan microarrays |
| Printing conditions | The details are included in SI section Generation of glycan microarrays |
| 5. Glycan Microarray with “Map” | |
| Array layout | The array layout of covalent and noncovalent NGL array are described in SI section Generation of glycan microarrays |
| Glycan identification and quality control | The details of glycan sequences are included in SI section Preparation of sequence defined glycans tagged with FAA including characterization on HPLC and MALDI MS. The quality control of the arrayed glycans was performed using antibodies and lectins listed in SI section Glycan binding proteins (GBPs). |
| 6. Detector and Data Processing | |
| Scanning hardware | GenePix 4300A (Molecular Devices, Berkshire, UK) |
| Scanner settings | The scanner settings for covalent and NGL arrays described in SI section Microarray analysis of glycan binding respectively |
| Image analysis software | ScanArray Express software (PerkinElmer LAS, Beaconsfield, UK) for the CLL array and GenePix® Pro 7 (Molecular Devices, Berkshire, UK) for the arrays. The details are included in SI section Microarray analysis of glycan binding |
| Data processing | The gpr files were entered into Excel and processed from there. No particular normalization method or statistical analysis was used. |
| 7. Glycan Microarray Data Presentation | |
| Data presentation | Binding results are presented as histogram charts in the main text and additional figures in SI Figure S11-16  Relative intensities of binding (matrixes) are included in the additional Excel files. |
| 8. Interpretation and Conclusion from Microarray Data | |
| Data interpretation | No software or algorithms were used to interpret processed data. |
| Conclusions | Different glycan binding systems have different preferences for the covalent and noncovalent arrays as summarised in Table S7. |

Appended Excel files

Excel Table 1. Probelist of covalent sequence defined array

Included in additional Excel file.

Excel Table 2. Probelist of noncovalent sequence defined array

Included in additional Excel file.

Excel Table 3. Binding data matrix of antibodies and lectins on sequence defined array

Included in additional Excel file.

Excel Table 4. Binding data matrix of GBPs on sequence defined array

Included in additional Excel file.

List of references

(1) Liu, Y.; Childs, R. A.; Palma, A. S.; Campanero-Rhodes, M. A.; Stoll, M. S.; Chai, W.; Feizi, T. Neoglycolipid-Based Oligosaccharide Microarray System: Preparation of NGLs and Their Noncovalent Immobilization on Nitrocellulose-Coated Glass Slides for Microarray Analyses. In *Carbohydrate Microarrays: Methods and Protocols*, Chevolot, Y. Ed.; Humana Press, 2012; pp 117-136.

(2) Liu, L.; Prudden, A. R.; Bosman, G. P.; Boons, G. J. Improved isolation and characterization procedure of sialylglycopeptide from egg yolk powder. *Carbohydrate research* **2017**, *452*, 122-128.

(3) Bigge, J. C.; Patel, T. P.; Bruce, J. A.; Goulding, P. N.; Charles, S. M.; Parekh, R. B. Nonselective and Efficient Fluorescent Labeling of Glycans Using 2-Amino Benzamide and Anthranilic Acid. *Analytical Biochemistry* **1995**, *230* (2), 229-238. DOI: <https://doi.org/10.1006/abio.1995.1468>

(4) Song, X.; Xia, B.; Stowell, S. R.; Lasanajak, Y.; Smith, D. F.; Cummings, R. D. Novel fluorescent glycan microarray strategy reveals ligands for galectins. *Chemistry & biology* **2009**, *16 1*, 36-47.

(5) Li, L.; Liu, Y.; Ma, C.; Qu, J.; Calderon, A. D.; Wu, B.; Wei, N.; Wang, X.; Guo, Y.; Xiao, Z.; et al. Efficient chemoenzymatic synthesis of an N-glycan isomer library. *Chemical Science* **2015**, *6* (10), 5652-5661, 10.1039/C5SC02025E. DOI: 10.1039/C5SC02025E

(6) Gao, C.; Wei, M.; McKitrick, T. R.; McQuillan, A. M.; Heimburg-Molinaro, J.; Cummings, R. D. Glycan Microarrays as Chemical Tools for Identifying Glycan Recognition by Immune Proteins. *Front Chem* **2019**, *7*, 833. DOI: 10.3389/fchem.2019.00833

(7) Blixt, O.; Head, S.; Mondala, T.; Scanlan, C.; Huflejt, M. E.; Alvarez, R.; Bryan, M. C.; Fazio, F.; Calarese, D.; Stevens, J.; et al. Printed covalent glycan array for ligand profiling of diverse glycan binding proteins. *Proc Natl Acad Sci U S A* **2004**, *101* (49), 17033-17038. DOI: 10.1073/pnas.0407902101

(8) Wu, Z.; Liu, Y.; Ma, C.; Li, L.; Bai, J.; Byrd-Leotis, L.; Lasanajak, Y.; Guo, Y.; Wen, L.; Zhu, H.; et al. Identification of the binding roles of terminal and internal glycan epitopes using enzymatically synthesized N-glycans containing tandem epitopes. *Organic & Biomolecular Chemistry* **2016**, *14* (47), 11106-11116, 10.1039/C6OB01982J. DOI: 10.1039/C6OB01982J

(9) Streit, A.; Yuen, C. T.; Loveless, R. W.; Lawson, A. M.; Finne, J.; Schmitz, B.; Feizi, T.; Stern, C. D. The Le(x) carbohydrate sequence is recognized by antibody to L5, a functional antigen in early neural development. *J Neurochem* **1996**, *66* (2), 834-844. DOI: 10.1046/j.1471-4159.1996.66020834.x

(10) Young, W. W.; Portoukalian, J.; Hakomori, S. Two monoclonal anticarbohydrate antibodies directed to glycosphingolipids with a lacto-N-glycosyl type II chain. *Journal of Biological Chemistry* **1981**, *256* (21), 10967-10972. DOI: <https://doi.org/10.1016/S0021-9258(19)68541-8>

(11) Galustian, C.; Childs, R. A.; Stoll, M.; Ishida, H.; Kiso, M.; Feizi, T. Synergistic interactions of the two classes of ligand, sialyl-Lewis(a/x) fuco-oligosaccharides and short sulpho-motifs, with the P- and L-selectins: implications for therapeutic inhibitor designs. *Immunology* **2002**, *105* (3), 350-359. DOI: 10.1046/j.1365-2567.2002.01369.x

(12) Olsvik, O.; Wahlberg, J.; Petterson, B.; Uhlén, M.; Popovic, T.; Wachsmuth, I. K.; Fields, P. I. Use of automated sequencing of polymerase chain reaction-generated amplicons to identify three types of cholera toxin subunit B in Vibrio cholerae O1 strains. *Journal of Clinical Microbiology* **1993**, *31* (1), 22-25. DOI: doi:10.1128/jcm.31.1.22-25.1993

(13) Lebens, M.; Holmgren, J. Structure and arrangement of the cholera toxin genes in Vibrio cholerae O139. *FEMS Microbiology Letters* **1994**, *117* (2), 197-202. DOI: 10.1111/j.1574-6968.1994.tb06764.x (acccessed 6/15/2023).

(14) Sandkvist, M.; Hirst, T. R.; Bagdasarian, M. Alterations at the carboxyl terminus change assembly and secretion properties of the B subunit of Escherichia coli heat-labile enterotoxin. *Journal of Bacteriology* **1987**, *169* (10), 4570-4576. DOI: doi:10.1128/jb.169.10.4570-4576.1987

(15) Bojar, D.; Meche, L.; Meng, G.; Eng, W.; Smith, D. F.; Cummings, R. D.; Mahal, L. K. A Useful Guide to Lectin Binding: Machine-Learning Directed Annotation of 57 Unique Lectin Specificities. *ACS Chemical Biology* **2022**, *17* (11), 2993-3012. DOI: 10.1021/acschembio.1c00689

(16) Streit, A.; Yuen, C.-T.; Loveless, R. W.; Lawson, A. M.; Finne, J.; Schmitz, B.; Feizi, T.; Stern, C. D. The Lex Carbohydrate Sequence Is Recognized by Antibody to L5, a Functional Antigen in Early Neural Development. *Journal of Neurochemistry* **1996**, *66* (2), 834-844. DOI: <https://doi.org/10.1046/j.1471-4159.1996.66020834.x>

(17) Li, L.; Guan, W.; Zhang, G.; Wu, Z.; Yu, H.; Chen, X.; Wang, P. G. Microarray analyses of closely related glycoforms reveal different accessibilities of glycan determinants on N-glycan branches. *Glycobiology* **2019**, *30* (5), 334-345. DOI: 10.1093/glycob/cwz100 (acccessed 6/14/2023).

(18) Larkin, M.; Ahern, T. J.; Stoll, M. S.; Shaffer, M.; Sako, D.; O'Brien, J.; Yuen, C. T.; Lawson, A. M.; Childs, R. A.; Barone, K. M.; et al. Spectrum of sialylated and nonsialylated fuco-oligosaccharides bound by the endothelial-leukocyte adhesion molecule E-selectin. Dependence of the carbohydrate binding activity on E-selectin density. *J Biol Chem* **1992**, *267* (19), 13661-13668.
